# Cytotoxicity and Characterization of 3D-Printable Resins Using a Low-Cost Printer for Muscle-based Biohybrid Devices

**DOI:** 10.1101/2024.10.31.621115

**Authors:** Ashlee S. Liao, Kevin Dai, Alaeddin Burak Irez, Anika Sun, Michael J. Bennington, Saul Schaffer, Bhavya Chopra, Ji Min Seok, Rebekah Adams, Yongjie Jessica Zhang, Victoria A. Webster-Wood

**Author notes:** Email address (Victoria A. Webster-Wood), URL: https://www.engineering.cmu.edu/borg (Victoria A. Webster-Wood). These authors contributed equally (Kevin Dai, Alaeddin Burak Irez). These authors contributed equally (Anika Sun, Michael J. Bennington).

## Abstract

Biohybrid devices integrate biological and synthetic materials. The selection of an appropriate synthetic material to interface with living cells and tissues is crucial due to cellular chemical and mechanical sensitivities. As such, the stiffness of the material and its biocompatibility while in direct contact must be considered. In this study, the material properties and biocompatibility of six commercially available, 3D printable resins (three rigid and three elastomeric) were assessed for their suitability for biohybrid actuators. To characterize the material, uniaxial tension and compression tests with post-hoc Hookean and Yeoh model analyses were conducted for both nonsterile and sterilized (ethanol-soaking or autoclaved) samples. The mechanical properties of the elastomeric resins were minimally impacted by the different sterilization techniques. However, both Phrozen AquaGray 8K and Liqcreate Bio-Med Clear rigid resins were significantly softer in tensile tests after sterilization, and AquaGray became far more ductile. Asiga DentaGUIDE was much more stable in its mechanical properties than the other rigid resins. It was also shown that long-term exposure to saline solutions leads to a decrease in the Young’s moduli of these rigid resins before any sterilization has occurred. The print fidelity was also assessed for nonsterile and sterilized samples via manual scoring to determine the impacts of the sterilization processes on the part fidelity. Sterilization techniques had a minimal impact on print fidelity for both elastomeric and ridged resins with two exceptions. In both Formlabs Silicone 40A IPA/BuOAc post-treatment and Phrozen AquaGrey 8K groups, ethanol/UV-sterilization caused more degradation compared to autoclave-sterilization. In addition to the material analyses, cytotoxicity analyses using calcein AM and ethidium homodimer-1 fluorescence markers were conducted by directly culturing C2C12, a common myoblast cell line used in bioactuators, with sterilized resin samples. Of the elastomeric resins, only Formlabs Silicone 40A was shown to have minimal impacts on cell viability. For the rigid resins, Asiga DentaGUIDE, Liqcreate Bio-Med Clear, and ethanol-sterilized Phrozen AquaGray 8K demonstrated minimal impacts on cell viability. Based on these analyses, Asiga DentaGUIDE and Formlabs Silicone 40A demonstrate potential for applications in biohybrid muscle-based actuators when using low-cost 3D printers.

## 1. Introduction

Biohybrid devices demonstrate potential for biomedical applications, such as regenerative medicine and pharmaceutical screening, due to the integration of synthetic and organic materials (Ricotti et al., 2017; Wang et al., 2022). Biohybrid devices have been developed to form a contractile heart tube (Bliley et al., 2022), create gas-permeable, cell-laden membranes towards a biohybrid lung (Wiegmann et al., 2016), and integrate directly with peripheral nerves for regenerative bioelectronics (Rochford et al., 2023). Since biohybrid devices are reliant on the close relationship between synthetic and organic materials, selecting the appropriate nonliving materials to interface with the living materials is key to developing a functional system (Liao et al., 2024). When selecting an appropriate synthetic material, key parameters to consider are material mechanical properties and biocompatibility (Liao et al., 2024).

The rigidity of a material contributes to the stiffness of the microenvironment in biohybrid devices, which is known to affect cellular maturation and behavior (Janmey et al., 2020). Matching the stiffness of the scaffold or structure to the needs of the tissue is therefore critical. Native extracellular matrix (ECM) varies greatly in stiffness throughout the body, from 1-3 kPa in the brain to 15-40 GPa in bone (Yi et al., 2022). Deviations from the native stiffness can cause a mechanical stimulus, inducing differing cellular responses such as changes to cell morphologies, differentiation, and cellular adhesion (Janmey et al., 2020). Muscle, which is of particular interest in biohybrid devices, is known to be mechanosensitive with their proliferation and differentiation especially affected by environmental stiffness (Janmey et al., 2020; Boontheekul et al., 2007). Myotube striation was shown to be affected by the flexibility of the culture substrate (Engler et al., 2004). If the substrate stiffness deviates from the stiffness of normal muscle (passive Young’s modulus of ≈ 12 kPa), myotubes fail to form properly striated fibers (Engler et al., 2004). In biohybrid actuators, these stiffnesses need to be carefully considered since muscle is commonly used to produce force (Liao et al., 2024). Not only are the stiffnesses important to ensure proper maturation of the cellular components, but synthetic materials can also serve as stiff structural supports (Finkel et al., 2023) or flexible skeletons capable of motion (Raman et al., 2017). Therefore, the stiffness of any interfacing artificial materials must be carefully considered based on their supportive functions to muscle in biohybrid actuators.

With the advent of additive manufacturing, 3D-printable resins provide a source of materials with a wide range of mechanical properties, enabling the rapid production of customized medical and biohybrid devices (select resins listed in Table 1). Several commercially available resins are now available that have been advertised to be biocompatible (Liao et al. (2024); Guttridge et al. (2022), and a subset can be found in Table 2). However, ISO 10993-1:2018 requires that tests must be designed and implemented to assess biocompatibility in only specific material use cases instead of a general assessment to be interpreted for all potential uses (International Organization for Standardization, 2018). Based on these standards, before any commercial resins are used for biohybrid applications, their biocompatibility should be reevaluated for their intended application (Liao et al., 2024). Therefore, for the specific use in biohybrid actuators, these resins should be tested in direct contact with the cultured muscle cells, when fabricated and cured using accessible manufacturing hardware.

**Table 1:** Mechanical properties of commercially available, 3D printable resins as reported by their respective manufacturers. ^†^For DentaGUM, Choi et al. (2021) measured the elastic modulus and the tensile strength. For Liqcreate Bio-Med Clear, the manufacturer’s reported values are for samples *before steam sterilization and **after steam sterilization at 121 °C).

| Resin | Tensile Modulus (MPa) | Ultimate Tensile Strength (MPa) | Elongation at Break | Shear Modulus (MPa) | Shore |
| --- | --- | --- | --- | --- | --- |
| Formlabs Silicone 40A | Not Listed | 5.0 | 230% | Not Listed | 40A |
| 3Dresyns Bioflex A10 MB | <1 | <1 | >>300% | Not Listed | 10A |
| Asiga DentaGUM | Not Listed<br>1.84<sup>†</sup> | Not Listed<br>3.41<sup>†</sup> | Not Listed | Not Listed | Not Listed |
| Asiga DentaGUIDE | Not Listed | Not Listed | Not Listed | Not Listed | Not Listed |
| Phrozen AquaGray 8K | 1196-1915 | 21-35 | 12-20% | Not Listed | 75-82D |
| Liqcreate Bio-Med Clear | 1700*<br>2200** | 78*<br>75** | 5-10% | 2200*<br>2500** | 85D |

**Table 2:** General overview of the use cases of six commercially-available, 3D-printable resins.

| Resin | Intended Use | Advertised Biocompatibility | Material Class |
| --- | --- | --- | --- |
| Formlabs Silicone 40A | Various, including medical | Pending | Elastomeric |
| 3Dresyns Bioflex A10 MB | Biomedical applications | Biocompatible | Elastomeric |
| Asiga DentaGum | 3D Printed Gingiva Mask, Not for medical use | Not Listed | Elastomeric |
| Asiga DentaGuide | 3D-Printed Dental Surgical Guides, Short-term use (<30 days) | CE Class I | Rigid |
| Phrozen AquaGray 8K | General 3D Printing | Not Listed | Rigid |
| Liqcreate Bio-Med Clear | Medical Devices | ISO 10993-5:2009<br>ISO 10993-10:2021<br>ISO 10993-23:2021 | Rigid |

In addition to a diverse selection of commercially available resins, the rapid development of additive manufacturing technologies has led to various printers that range in printing feature resolution and cost. Currently, commercial 3D resin printers, which commonly use stereolithography (SLA) or digital light processing (DLP), can achieve resolutions on the order of tens of microns and a cost on the order of $1000s USD (Formlabs, 2024c; Lalwani, 2020). For example, Asiga’s DLP printer, MAX X27, can achieve a 27 μm resolution (Asiga, 2024b) at a cost of $9,990 as of April 2024 (Systems, 2019), and Formlabs’ SLA printer for healthcare, Form 3B+, can achieve a 25 μm XY resolution with an 85 μm laser spot size (Formlabs, 2024b) at a cost of $4,429 as of April 2024 (Formlabs, 2024a)). With the increase in accessibility of resin printers entering the market for hobbyist crafters, the cost of high-resolution printers has drastically decreased to less than $1,000 with the use of a liquid crystal display (LCD), also often referred to as masked stereolithography (MSLA) (Formlabs, 2024c; Lalwani, 2020). For example, Phrozen’s Sonic Mini 8K LCD printer can achieve a 22 μm resolution at a cost of $485 (Phrozen, a). However, these lower-cost printers commonly require more calibration and tuning per resin (Formlabs, 2024c). Therefore, the printer type and calibration conditions may affect the final cured resin product. In particular, if resins are under-cured due to the capabilities of the printer and curing station, residual monomers are likely to be present and are known to be cytotoxic. Thus, the biocompatibility and material properties must be reevaluated based on not only the resin type but also the printer, curing, and wash approach used.

In this work, the cytotoxicity, mechanical properties, and print fidelity of six commercial resins (Table 3) were printed using a low-cost Phrozen Sonic Mini 8K and evaluated for biohybrid actuator applications when manufactured using this accessible, low-cost equipment. Of the six resins, three were considered rigid, and three were considered elastomeric to investigate a range of stiffnesses. Both mechanical testing and sample feature fidelity were evaluated for nonsterile and sterilized (autoclave or ethanol/UV) samples since material properties may change after sterilization. Cytotoxicity testing was only conducted with sterilized samples. Due to the inherent dependence of resin curing and, therefore, cytotoxicity on the printer and curing station parameters, these results should be interpreted in the context of using this low-cost, accessible equipment with the goal of broadening access to materials suitable for biohybrid robotics.

**Table 3:** Resins used for the analysis with their associated post-print treatments and related mechanical testing standards used to guide their assessment. All resins had sample groups with identical isopropanol (IPA) treatment. Formlabs Silicone 40A and 3Dresyns Bioflex A10 MB were also assessed with a second sample group that was treated based on manufacturer’s recommendations (80% IPA/20% *n*-butyl acetate (BuOAc) and UNW2, respectively).

|  | Resin | Post-Print Treatment | Tensile Test | Compression Test |
| --- | --- | --- | --- | --- |
| **Rigid** | Phrozen AquaGray 8K | IPA |  |  |
|  | Liqcreate Bio-Med Clear | IPA | ASTM D638<br>5 mm/min | ASTM D695<br>1.3 mm/min |
|  | Asiga DentaGUIDE | IPA |  |  |
| **Elastomeric** | Formlabs Silicone 40A | IPA<br>IPA/BuOAc |  |  |
|  | 3Dresyns Bioflex A10 MB | IPA<br>UNW2 | ASTM D412<br>500 mm/min | ASTM D575<br>12 mm/min |
|  | Asiga DentaGUM | IPA |  |  |

## 2. Material and methods

To identify suitable 3D-printable resins for biohybrid muscle-based devices, several commercial resins were characterized. These resins ranged in their mechanical properties (Table 1) and advertised biocompatibility (Table 2). However, the ISO 10993-1:2018 standard recommends that endpoint tests should be completed based on specific use cases (International Organization for Standardization, 2018). Furthermore, all of the resins, with the exception of Liqcreate Bio-Med Clear, do not report mechanical properties after a sterilization treatment. Therefore, for the use case in biohybrid devices where there may be direct contact with cultured muscle cells and tissues, the cytotoxicity, tensile, and compressive properties were measured and compared with and without common sterilization techniques used in biohybrid robotics.

### 2.1. Sample Fabrication

Six commercially available resins were selected for the analyses (Table 3). All resins were printed using a low-cost Phrozen Sonic Mini 8K resin printer (Phrozen Tech Co., LTD., Hsinchu, Taiwan). After printing, the samples were submerged in a container with 100% isopropanol (IPA) and placed in an ultrasonic bath for 5 minutes. This wash step was repeated once in a second container of 100% IPA. Each resin had its own dedicated first and second wash containers. After the two washes, the samples were dried and cured in a Phrozen Curing Station (405 nm UV lights; Phrozen Tech Co., LTD., Hsinchu, Taiwan) with a fan period of 30 minutes followed by a cure period of 15 minutes. Then, the samples were flipped and cured for another 15 minutes. It is important to note that this printer and curing system, while broadly accessible due to their low cost, has a lower light power than other, less accessible, commercially available systems. Therefore, the tests reported here should only be interpreted in the context of using such equipment as lower power systems have a higher risk of resulting in under-cured resins, which are known to present cytotoxicity risks.

For two of the resins, Formlabs Silicone 40A and 3Dresyns Bioflex A10 MB, the respective manufacturers recommended alternative post-processing. Therefore, in addition to the IPA-processed samples, additional samples were fabricated for these resins and post-processed using their manufacturers’ protocols. The additional Formlabs Silicone 40A samples were washed using the ultrasonic bath in an 80% IPA and 20% *n*-butyl acetate (BuOAc) solution. To wash the samples, the samples were placed in a container filled with IPA/BuOAc. This container was placed in an ultrasonic water bath for 10 minutes. This was repeated once in a second IPA/BuOAc container for a total of two wash cycles. After the washes, the samples were dried in the Phrozen Curing Station (405 nm UV light, Phrozen Tech Co., LTD., Hsinchu, Taiwan) with a 15-minute fan period. Afterward, they were transferred to a Petri plate filled with water and cured in the Phrozen Curing Station (Phrozen Tech Co., LTD., Hsinchu, Taiwan) for 45 minutes. Lastly, they were removed from the water and stored in dry Petri plates before sterilization or testing.

For the second set of 3Dresyns Bioflex A10 MB samples, samples were washed twice in containers filled with Cleaning Fluid UNW2 Bio (UNW2) (P20759; 3Dresyns, Barcelona, Catalonia, Spain) for 5 minutes each in an ultrasonic bath. After the washes, the samples were cured in the Phrozen Curing Station (Phrozen Tech Co., LTD., Hsinchu, Taiwan) for 6 minutes while submerged in UNW2. Then, the samples were briefly dipped in IPA and allowed to dry. The duration in IPA was not timed and ranged from a few seconds to a couple of minutes as individual samples were removed from the IPA in a random order to dry on a paper towel. Afterward, they were submerged in UNW2 for 1 hour at 65 °C. Finally, they were submerged in deionized water for 1 hour at 65 °C. After the final post-printing process, the samples were removed from the water and stored in dry Petri plates until they were sterilized or used for testing.

In addition to the resin samples, polydimethylsiloxane (SYLGARD 184; Dow Corning, Midland, MI, USA) with a 10:1 base-to-curing-agent (w/w) ratio was used as a negative control (no impact on cell viability) in the biocompatibility assays. PDMS was cast in a Petri plate and cured for 6 hours at 65 °C (master sample). After the PDMS was cured, cylindrical samples were cut from the master sample using a 2-mm biopsy punch.

For consistency, all of the resins post-processed with the same IPA washes and curing procedure were simultaneously assessed for cytotoxicity. Since the Formlabs Silicone 40A samples were also assessed with an alternative, manufacturer’s post-processing procedure (IPA/BuOAc), the cytotoxicity assessment of these samples was done in a separate culture period than the assessment for the other IPA-processed resins. Therefore, separate PDMS control samples were used for this group. The PDMS samples were cut from the same master sample using a 2-mm biopsy punch.

### 2.2. Sample Sterilization

To sterilize the resins, two common methods of sterilization used in biohybrid robotics were explored: (1) autoclaving and (2) submersion in 70% ethanol with UV exposure. For autoclaving select samples, a gravity cycle at 121 °C for 45 minutes with 15 minutes of dry time was used (ADV-PRO; Consolidated Sterilizer Systems, Billerica, MA, USA).

For the ethanol/UV sterilization method, samples were submerged in 70% ethanol for 1 hour at room temperature. Afterward, they were swirled three times in 1X phosphate-buffered saline (PBS, 10-010-049; Gibco ThermoFisher, Waltham, MA, USA) and, then, left to dry overnight in the biosafety cabinet (1300 Series A2, Model 1377; ThermoFisher, Waltham, MA, USA) with 1 hour of ultraviolet (UV) light exposure from the biosafety cabinet’s regular sterilization cycle.

### 2.3. Cytotoxicity Analysis

Following the manufacturing process described above, a biocompatibility analysis was conducted to identify 3D-printable resins that can be used in direct contact with cultured cells over several days or weeks when produced with the low-cost equipment tested in this work. The appropriate biocompatibility analysis for this specific use was determined based on guidance from the International Organization for Standardization (ISO) “Biological Evaluation of Medical Devices” standards 10993-1:2018 (International Organization for Standardization, 2018), 10993-5:2009 (International Organization for Standardization, 2009), and 10993-12:2021 (International Organization for Standardization, 2021). Based on 10993-1:2018 (International Organization for Standardization, 2018), the use case was considered for prolonged periods of exposure (24 hours to 30 days), cytotoxicity tests with cell culture would be appropriate evaluation in line with ISO 10993-5:2009 (International Organization for Standardization, 2009), and samples should be prepared based on ISO 10993-12:2021 (International Organization for Standardization, 2021).

Based on 10993-5:2009 (International Organization for Standardization, 2009), a direct contact test was determined to be the appropriate analysis for prolonged exposure to cultured cells. The established mouse myoblast cell line, C2C12 (CRL-1772; ATCC, Manassas, Virginia, USA), was selected as a commonly established cell line for biohybrid muscle-based devices. For contact with the cells, cylindrical samples were fabricated with a flat face that occupied approximately 10% of the surface area of a single well of a 96-well plate. The positive control used was 70% ethanol since it will cause a cellular response - in this case, cell death. The negative control, which should not cause any cellular responses, used was PDMS samples created in 10:1 (w/w) ratio of base:curing agent and a 2-mm biopsy punch (refer to Section 2.1).

#### 2.3.1. Cell Culture

Cryopreserved C2C12 cells were thawed and plated in 96-well plates (655087; Greiner Bio-One, Kremsmünster, Austria) at a density of 2,100 cells/well. The plating time corresponded with the beginning of the 9th passage for the cells. C2C12 growth media composed of high-glucose Dulbecco’s Minimum Essential Medium (DMEM, 11965126; Gibco ThermoFisher, Waltham, MA, USA) supplemented with 10% fetal bovine serum (A5256801; Gibco ThermoFisher, Waltham, MA, USA), 1% penicillin/streptomycin (10,000

U/mL, 15140122; Gibco ThermoFisher, Waltham, MA, USA), and 1% L-glutamine (200 mM, 25030081; Gibco ThermoFisher, Waltham, MA, USA). The cells were cultured for 2 days before a complete media change and the introduction of resin samples. Cytotoxicity measurements were taken 72 hours after the introduction of resin samples. There were 6 replicate wells for each resin type, the negative (PDMS) controls, the positive (70 % ethanol) controls, and the blank (cell only, no resin exposure) controls.

#### 2.3.2. Cytotoxicity (Plate Reader and Imaging)

Cytotoxic effects were measured using ThermoFisher LIVE/DEAD Viability/Cytotoxicity Kit (L3224; Invitrogen ThermoFisher, Waltham, MA, USA), which uses fluorescence dyes calcein-AM and ethidium homodimer-1 for identifying live and dead cells, respectively. For the wells with samples and the negative controls (PDMS), before washing the cells and applying the dyes, all of the resin samples were carefully removed to minimize the risk of cells detaching from the culture surface due to the movement of the samples during the wash steps. For the positive controls, the media was first aspirated and replaced with cold 70% ethanol and incubated at room temperature for at least 15 minutes prior to the wash steps.

After sample removal, the wells were washed three times with 1X PBS. After the last rinse, 100 *μL* of 1X PBS was added to each well. For the wells that contained the samples and the negative control wells, 100 *μL* of the dye solution containing 1 *μM* calcein-AM and 2 *μM* of ethidium homodimer-1 was added to each well. For the positive and blank controls, 100 *μL* of only 1 *μM* calcein-AM was added to half of the wells, and 100 *μL* of only 2 *μM* ethidium homodimer-1 was added to the remaining half, as recommended by the manufacturer’s protocol (Probes, 2005).

After 30 minutes of incubation, the fluorescence was measured using a plate reader (Synergy H1; Agilent BioTek, Santa Clara, CA, USA) using the fluorescence endpoint test of the entire well, read from the bottom of the well plate, for each sample. A monochromater with a xenon flash light source with high lamp energy, extended gain, and extended dynamic range was used with two filter sets: (1) default “Calcein” setting (485 nm excitation, 526 nm emission) and (2) default “Texas Red” setting (584 nm excitation, 625 nm emission). For the measurements, a normal read speed was used with a 100 ms delay, 10 data point measurements per well, and a 7 mm read height. During the measurements, the plate lids of the 96-well plates were still used. For the analysis, the ratio of calcein AM fluorescence and ethidium homodimer-1 (Texas Red) fluorescence (CalAM:EthD-1) was used.

Once the plate was read, the wells were imaged using a 5 MP sCMOS mono camera on an inverted Echo Revolution microscope under epifluorescence with the FITC and TXRED LED light cubes at 10X magnification (Revolution; Echo Bico, San Diego, CA, USA).

#### 2.3.3. Cytotoxicity Statistical Analysis

All statistical analyses were completed using Minitab 2024 (v 22.1). Distributions of CalAM:EthD-1 were first tested for normality using the Anderson-Darling test and for equal variances using the Bartlett test. Based on this initial assessment, a balanced analysis of variance (ANOVA) was selected to compare the groups with a post-hoc Bonferroni multiple comparison between groups. A significance level *α* of 0.05 was selected for these analyses, with a *p* ≤ *α* used as the criterion to reject the null hypothesis in which there is no association between CalAM:EthD-1 responses and the terms (resin type and sterilization method).

### 2.4. Materials Characterization

#### 2.4.1. Mechanical Behavior

To assess the effects of the sterilization method (refer to Section 2.2) on the mechanical properties of each resin, tensile and compression tests (Table 3) were conducted on nonsterile, ethanol/UV-sterilized, and autoclaved samples (Figure 1). Prior to the tests, the samples were conditioned in 1X PBS at 37°C for 16 hours. The tests were performed using an MTS Criterion Model 42 Universal Test Stand. Furthermore, during testing, the samples were maintained in 1X PBS at 37°C using an MTS Bionix EnviroBath. Just before inserting each sample into the grips for testing, the rest geometry of each sample was measured using Neiko digital calipers. For compression samples, the gauge length was measured as the full height of the puck. For tensile samples, the gauge length was measured as the grip separation after the sample was installed into the machine. Testing protocols for the tensile and compression tests were based on ASTM standards D695-23 (rigid compression), D575-91 (elastomeric compression), D638-14 (rigid tensile), and D412-06a (elastomeric tensile). The protocols in these standards are briefly summarized here:

**Figure 1:**
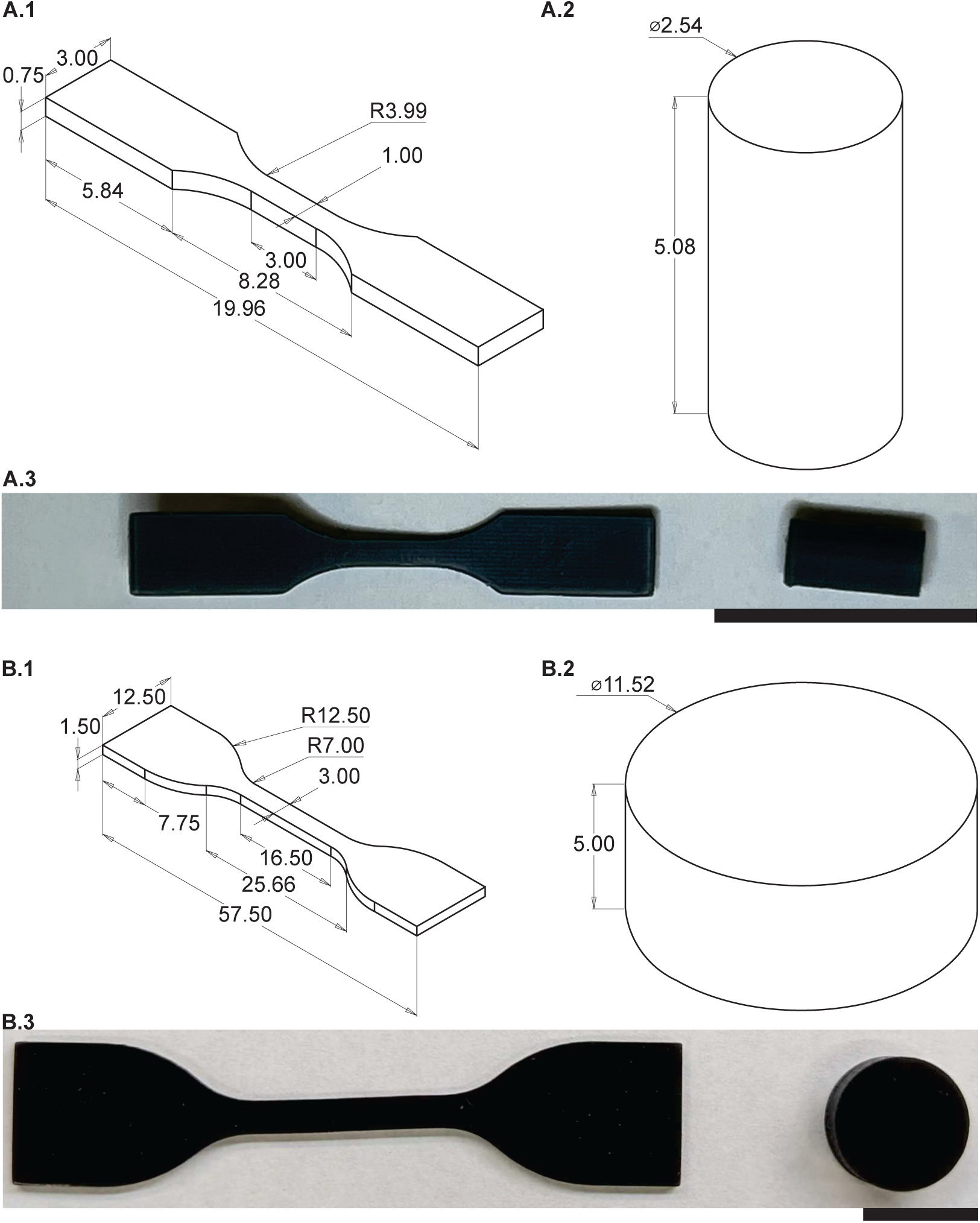
Specimen samples for the (A) rigid resin samples as per ASTM D638 and ASTM D695 recommendations and for the (B) elastomeric resin samples as per ASTM D412 and ASTM D575 recommendations. (1) Tensile sample specifications. (2) Compression sample specifications. The drawings are not to scale with respect to each panel. (3) Actual samples of (A.3) Phrozen AquaGray 8K and (B.3) Formlabs Silicone 40A with IPA-post-treatment. Scale bars represent 10 mm. Refer to supplementary materials for detailed files.

##### Rigid Compression Test

Compression tests were conducted on rigid samples (Figure 1A.2), with six samples tested from each group. For the testing, sandpaper was applied to both the lower and upper compression plates to prevent the samples from slipping. Subsequently, the samples were positioned at the center of the bottom plate, and their alignment with the upper plate was verified. Next, mild contact was made between the upper plate and the specimen by slowly adjusting the crosshead until a negligibly small load (0.05-0.1 N) was registered. During this stage, careful consideration was given to ensuring that no load was transferred to the specimen before conducting the tests. Subsequently, the test was commenced, and it was promptly halted when the force hit 48 N. Any force greater than 50 N has the potential to cause damage to the 50-N-load cell in the test machine. Consequently, the test was concluded at 48 N, and the test data was recorded.

##### Elastomeric Compression Test

Elastomeric compression tests were performed on the specimens shown in Figure 1B.2. The test was conducted utilizing the identical experimental configuration as the rigid compression tests. The test was conducted in three stages once the upper plate made slight contact, as indicated by the registration of a negligibly small load (0.05-0.1 N), with the test material. During the initial phase, a force of 1 N was applied to the sample at a testing rate of 1.3 mm/min. Once the force of 1 N was reached, in the subsequent phase, the upper plate of the test device remained stationary for a duration of 30 seconds. During the third step, a force was exerted at a constant test speed until it achieved a magnitude of 40 N. The test was subsequently halted. Only the data from the third stage was captured during this test, complying with the ASTM D575 test standard.

##### Rigid Tensile Test

Rigid tensile tests were performed on the samples shown in Figure 1A.1. Sandpaper was applied to the interior surfaces of the test grips to keep samples from slipping during testing, and samples were placed between the grips. This procedure ensured that the samples were properly aligned between the grips, which is critical for the tests’ uni-axial tensile nature. After inserting the specimen and ensuring alignment with the testing axis, the distance between the grips (gauge length) was measured using a caliper. The strain could not be measured using strain gauges or any other contact extensometer since the sample size was so small. Tests were then performed using displacement control at a crosshead speed of 5 mm/min (ASTM D638), with a 48 N force limit applied to avoid damaging the test machine’s 50-N-load cell.

##### Elastomeric Tensile Test

Elastomeric tensile tests were performed on the samples shown in Figure 1B.1. The most important aspects of these tests were correct sample alignment and maintaining the sample completely parallel to the vertical axis of the testbed (direction of force application) without loading it. After aligning the specimen and keeping it parallel to the vertical axis, the gauge length was measured using a caliper, and the test proceeded. The sample was stretched at a constant crosshead speed of 500 mm/min (ASTM D412). The test ended after either a complete failure or nearing the physical limits of the EnviroBath cabinet.

##### Data Analysis and Modeling

During all testing protocols, material displacement and reaction force were recorded by the MTS at 10 Hz. From these data, engineering stress (*σ_eng_*: force / rest cross sectional area) and strain (*E*) were calculated. For the dogbone samples, strain was calculated relative to the gauge length of the sample. For consistency between the different protocols with different durations (and thus different numbers of data points), the stress and strain data were resampled at 50 equally spaced strain values, and analysis was performed on this resampled data.

For the rigid samples, the Young’s modulus was calculated based on the ASTM D638 (Table 3). For consistency across samples with differently-shaped stress-strain curves, the ASTM standards for materials with no linear region were utilized. Briefly, from the stress-strain data, the slope between adjacent points was calculated and then smoothed with a 10-point moving average filter. The location of the maximum instantaneous slope was chosen as the linear region, and a linear function was fit to the surrounding 3 points. To account for the toe region in the data, the x-axis intercept of this line was then used as the zero point for the stain, and the strain data were adjusted accordingly. Finally, the slope of this line was recorded as the Young’s (or Compressive) Modulus of that sample. In addition to the modulus, for tensile tests, the Ultimate Tensile Strength (UTS) and Strain at Break were measured for each sample as the maximal stress and strain values, respectively. This procedure was followed for all rigid samples.

For the elastomeric material samples, a simple linear elastic model was insufficient to capture their elastic response. Instead, an incompressible Yeoh hyperelastic model was fit to the data (Yeoh and Fleming, 1997). The strain energy density function (Ψ) of the Yeoh model used here takes the form:

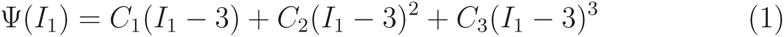

where *I*_1_ is the first invariant of the Cauchy deformation tensor (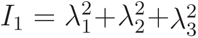, where *λ_i_* is the *i^th^* principle stretch), and *C_j_* are material parameters that were fit from the data. For the uniaxial experiments performed here, *λ*_1_ = *λ* = 1+*E* and, from incompressibility, *λ*_2_ = *λ*_3_ = *λ*^−0.5^. The engineering stress for the material can then be calculated (taking into account the hydrostatic pressure associated with the incompressibility constraint) as

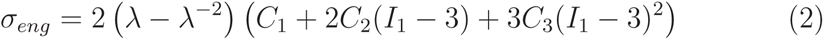

For uniaxial tests, *I*_1_ = *λ*^2^ + 2*/λ*. The equivalent Young’s Modulus can be calculated from the model parameters as *E* = 6*C*_1_ (Yeoh and Fleming, 1997). As with the rigid material, the toe region of the data was ignored. However, a similar standard procedure for doing so was not present for elastomeric materials. Instead, all stress below 5% of the max stress was considered potential toe region. Then, to adjust the strain levels, it was assumed that the beginning of the data was still in an approximately linear region. The first five data points were fit to a line (*σ* = *aλ* + *b*), and a displacement offset was calculated as *δL* = *L*_0_(*b/a* + 1), where *L*_0_ is the gauge length of the sample. This offset *δL* was then subtracted from the measured displacement, and the adjusted stretch was recalculated. This adjusted stretch was used for model fitting.

To ensure that the same model properly captured compression and tensile tests, the model parameters were calibrated using compression and tensile data simultaneously. However, tensile and compression data are not available for the same specimen. To overcome this limitation, a bootstrapping approach was utilized. For each step of the bootstrap, one compression and one tensile dataset were sampled to construct a dual-side stress-strain curve. Then, the three material parameters were fit by minimizing the squared error between the model and experimental data using MATLAB’s *fminsearch* optimization function. Penalty terms were added to ensure *C*_1_ and *C*_3_ were greater than 0 for stability, but *C*_2_ was allowed to be positive or negative (Yeoh and Fleming, 1997). These model parameters were then logged, and this process was repeated 50 times. The mean model response was then calculated as the model response given the mean model parameters. This is possible because the strain energy model is linear in its parameters. To determine the goodness of fit for the mean models, the coefficient of determination (*R*^2^) was calculated using all experimental data.

Scalar metrics were also calculated for the elastomer samples. UTS and Strain at Break could only be calculated for a subset of samples because not all samples failed during testing. Additionally, to align with metrics reported by select manufacturers, we calculate the stress at 50% and 100% (when possible) strain. This was found for each sample by linearly interpolating the experimental data and sampling at stretches of 1.5 and 2. For samples that did not reach 50% or 100% strain before the end of the test, no stress was calculated for that sample.

#### 2.4.2. Mechanical Properties Statistical Analysis

The mechanical properties extracted based on the raw data (Table 4) were statistically analyzed using Minitab (v. 22.1; Minitab, LLC., State College, PA, USA). The Anderson-Darling test and the Bartlett test were used to assess normality and equivalent variances, respectively. Based on these initial tests, a majority of the groups were normal and did not have significantly different variances. Therefore, a General Linear Model (ANOVA) was fit to the data with a post-hoc Bonferroni pairwise comparison. A level of significance of *α* = 0.05 was used for all analyses.

**Table 4:** Mechanical properties measured for each rigid and elastomeric (Elast.) resin. For rigid resins, stress at 50% and 100% strain was not considered as they break before achieving 50% strain. For 3Dresyns Bioflex A10 MB, the samples broke before achieving 100% strain. The ultimate tensile strength and elongation at break was not calculated for Formlabs Silicone 40A since a majority of those samples (32 of 36 samples) did not break due to machine limits. The compressive and tensile moduli were not calculated for elastomeric resins due to their hyperelasticity.

| Resin |  | Compressive Modulus | Young's (Tensile) Modulus | Ultimate Tensile Strength | Elongation at Break | Stress at 50% Strain | Stress at 100% Strain |
| --- | --- | --- | --- | --- | --- | --- | --- |
| Rigid | Asiga DentaGUIDE | ✓ | ✓ | ✓ | ✓ |  |  |
|  | Liqcreate Bio-Med Clear | ✓ | ✓ | ✓ | ✓ |  |  |
|  | Phrozen AquaGray 8K | ✓ | ✓ | ✓ | ✓ |  |  |
| Elast. | 3Dresyns Bioflex A10 MB |  |  | ✓ | ✓ | ✓ |  |
|  | Formlabs Silicone 40A |  |  |  |  | ✓ | ✓ |
|  | Asiga DentaGUM |  |  | ✓ | ✓ | ✓ | ✓ |

### 2.5. Print Fidelity

To assess the print fidelity for each resin using our low-cost 3D printer, samples with feature sizes ranging from 25 μm to 600 μm were printed (Figure 2). The samples were fabricated and sterilized as described in Sections 2.1-2.2. For the print fidelity assessment, the individual nonsterile, ethanol/UV-sterilized, and autoclaved samples were imaged using a 10MP CMOS camera on an AmScope SM-1T trinocular stereo microscope with a ring light and without a Barlow lens (SM-1TSZZ-144S-10M, AmScope). The samples were then analyzed by three co-authors for feature size and overall fidelity according to a set scoring rubric using Likert-type scales. The scorers were asked to select the location of the largest and smallest features by sectioned column and row. The scorers were also asked to rank overall print fidelity and print fidelity in quadrants on a 0-5 scale (0: no features visible; 5: all the features are identical to the target model from the CAD file). The quadrants were selected to distribute the number of features evenly, with quadrant 1 containing the largest features and quadrant 4 containing the smallest. (Figure 2) CAD files and additional experimental images can be found in the data repository associated with this work (See *Data Availability*). The ground truth score for each feature is collected in Table 5.

**Figure 2:**
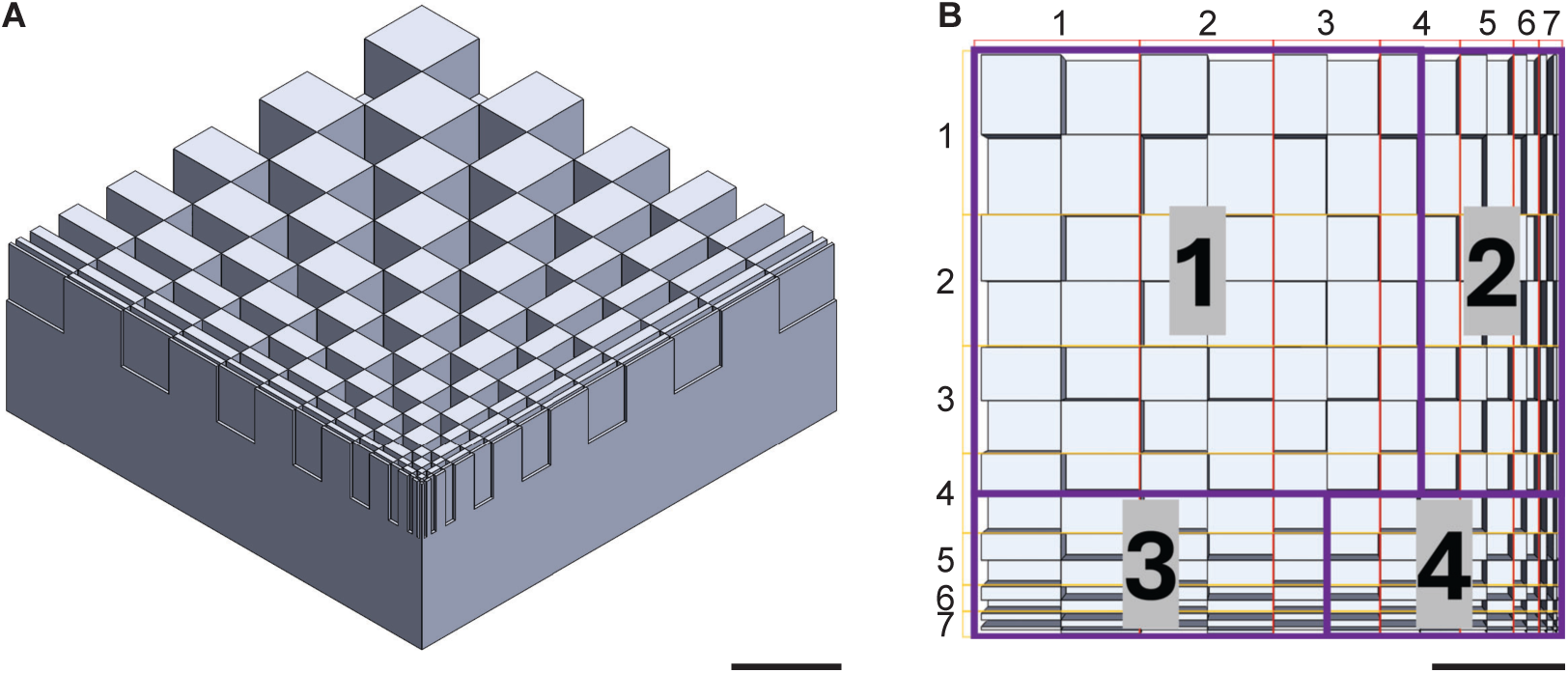
The printed samples had an alternating pattern of raised rectangles with decreasing length and width. The largest feature (Row 1, Column 1) had a length and width of 0.6mm and the smallest feature (Row 7, Column 7) had a length and width of 0.025mm. (A) Isometric view of the print fidelity sample CAD model (B) Row, column, and quadrant labels of the top face of the print fidelity sample. These labels tagged regions are used for the manual assessment of feature quality. All scale bars represent 1 mm. For detailed CAD models, see *Data Availability*

**Table 5:**
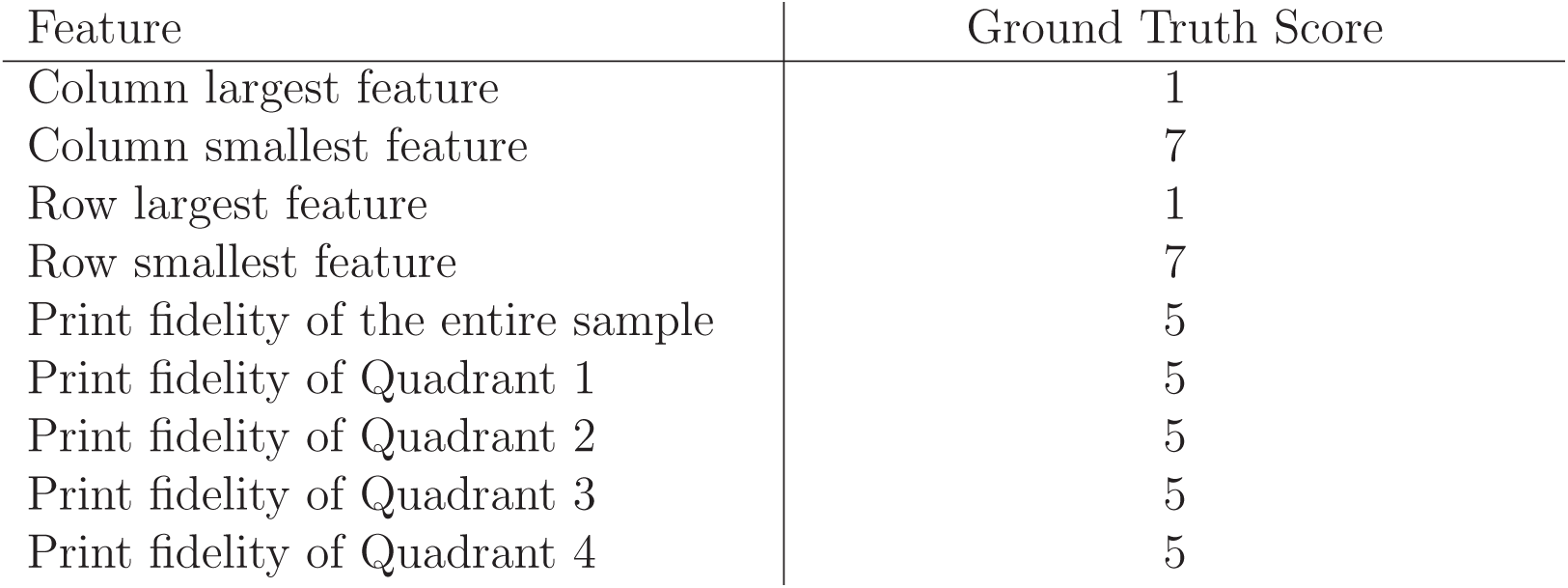
Table of each scored feature and the associated ground truth score.

| Feature | Ground Truth Score |
| --- | --- |
| Column largest feature | 1 |
| Column smallest feature | 7 |
| Row largest feature | 1 |
| Row smallest feature | 7 |
| Print fidelity of the entire sample | 5 |
| Print fidelity of Quadrant 1 | 5 |
| Print fidelity of Quadrant 2 | 5 |
| Print fidelity of Quadrant 3 | 5 |
| Print fidelity of Quadrant 4 | 5 |

#### 2.5.1. Print Fidelity Statistical Analysis

All statistical analyses were completed using Minitab 2024 (v 22.1). A mixed effects model (ANOVA) was selected to compare the groups with a post-hoc Bonferroni multiple comparison between groups. The categorical factors used in the mixed effects model were scorer ID, sterilization type, resin type, and interaction between sterilization and resin type, with the scorer ID being the random factor. These factors fit the data requirements for the model chosen. While the scoring responses were not continuous, Likert-type scale responses can be assessed using parametric tests (Sullivan and Artino, 2013). A significance level *α* of 0.05 was selected for these analyses, with a *p* ≤ *α* used as the criterion to reject the null hypothesis, where the null hypothesis is that there is no relationship between the factors and response.

## 3. Results and Discussion

### 3.1. Cytotoxicity

For the cytotoxicity assessment, the sterilization technique had minimal impact on the cellular response (Table 6 and Table 7), while the resin type and the interaction between resin and sterilization had significant impacts (Table 6). The quantitative assessment of CalAM:EthD-1 (Tables 6,8,7,9) was supported by the qualitative observations since minimal cellular responses were observed for four of the eight combinations of resin and post-treatment conditions, regardless of sterilization technique: Asiga DentaGUIDE (Figure 4D and Figure 5D), Liqcreate Bio-Med Clear (Figure 4C and Figure 5C), and Formlabs Silicone 40A, both IPA and IPA/BuOAc post-treatments (Figure 4G, Figure 5G, Figure 7B,D). In addition, ethanol/UV-sterilized Phrozen AquaGray 8K samples were not qualitatively observed to cause major cellular responses (Figure 4B). Surprisingly, the procedure of washing the cells with PBS and loading them with calcein AM and ethidium homodimer-1 (refer to Section 2.3.2) appeared to qualitatively impact C2C12 morphology. However, since all of the quantitative analysis was conducted using fluorescence intensity ratios, the change in morphology should not impact the cytotoxicity analysis since healthy morphology was observed in the cell-only (blank) controls before the dye was loaded (Figures 3, 6).

**Figure 3:**
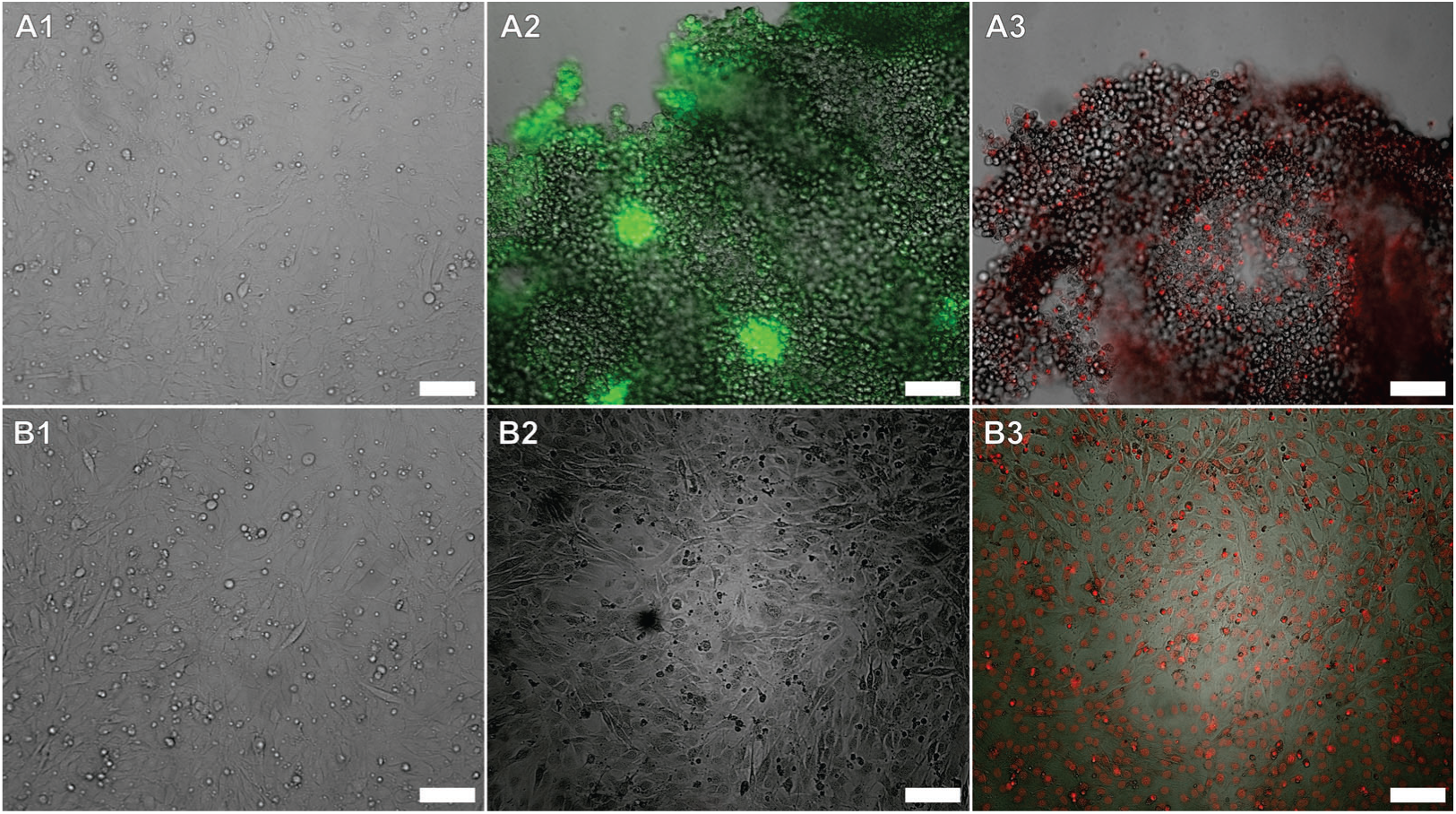
Brightfield and fluorescence microscopy images of cells that were not exposed to any resin samples. These controls were used for the biocompatibility assessment of all resins except for the Formlabs Silicone 40A post-treated with IPA/BuOAc. (A) represent the blank (live cell) controls. (B) represent the positive (dead cell) controls, with (B1) illustrating the cells prior to incubation with 70% ethanol. (1) are brightfield-only images of the cells prior to loading of any dye. (2) are brightfield images overlaid with fluorescence images of cells loaded only with calcein AM (green). (3) are brightfield images overlaid with fluorescence images of cells loaded only with ethidium homodimer-1 (red). All scale bars are 100 μm.

**Figure 4:**
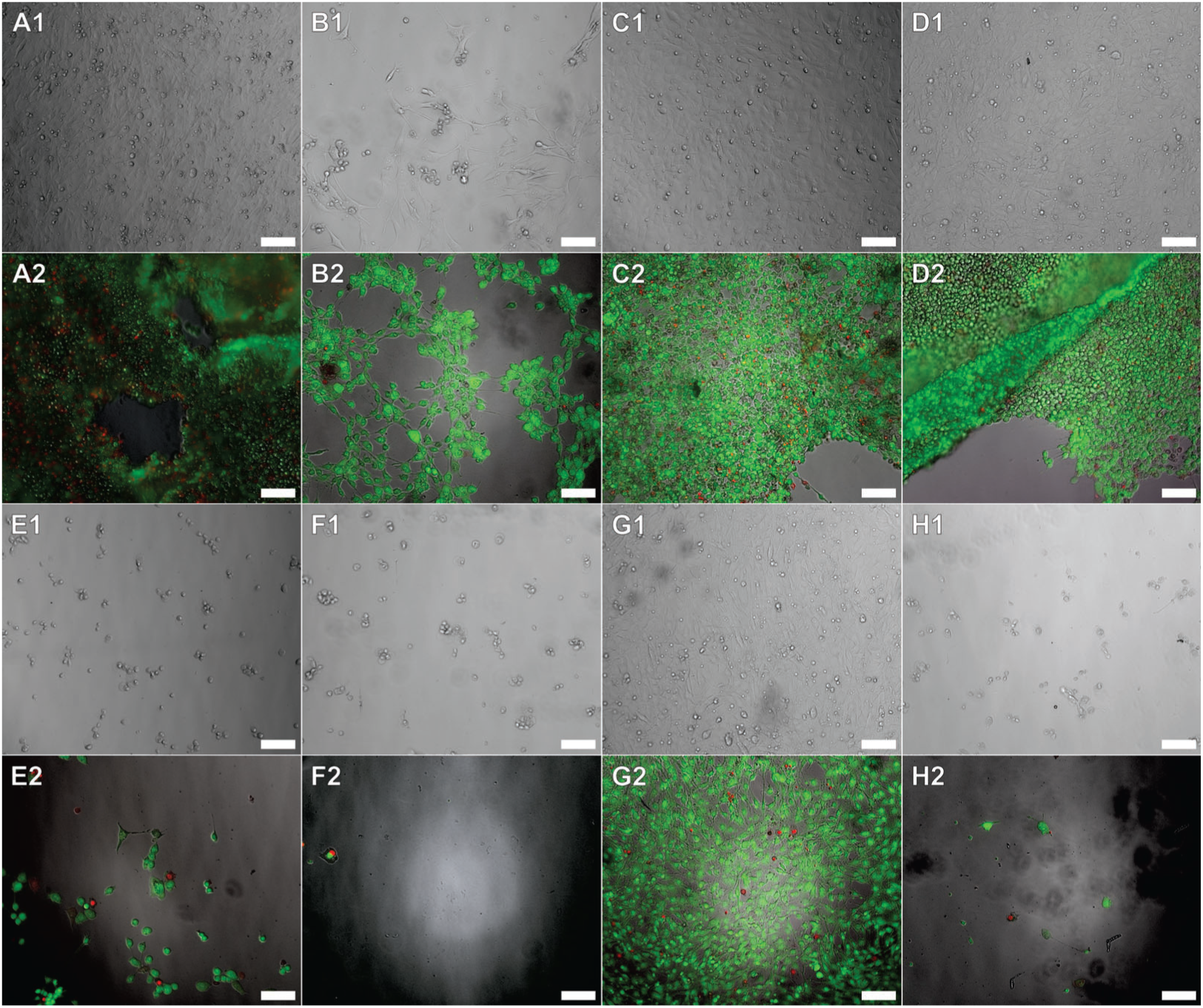
Brightfield (1) and fluorescence (2) microscopy images of cells that were exposed to ethanol/UV-sterilized samples. The brightfield-only images (1) were captured prior to PBS washing and dye loading. (2) are brightfield images overlaid with calcein AM (green) and ethidium homodimer-1 (red) fluorescence images of cells. The panels represent cells exposed to the following ethanol/UV-sterilized samples: (A) PDMS; (B) Phrozen AquaGray 8K; (C) Liqcreate Bio-Med Clear; (D) Asiga DentaGUIDE; (E) 3Dresyns Bioflex A10 MB IPA-post-treatment; (F) 3Dresyns Bioflex A10 MB UNW2-post-treatment; (G) Formlabs Silicone 40A IPA-post-treatment; (H) Asiga DentaGUM. All scale bars are 100 μm.

**Figure 5:**
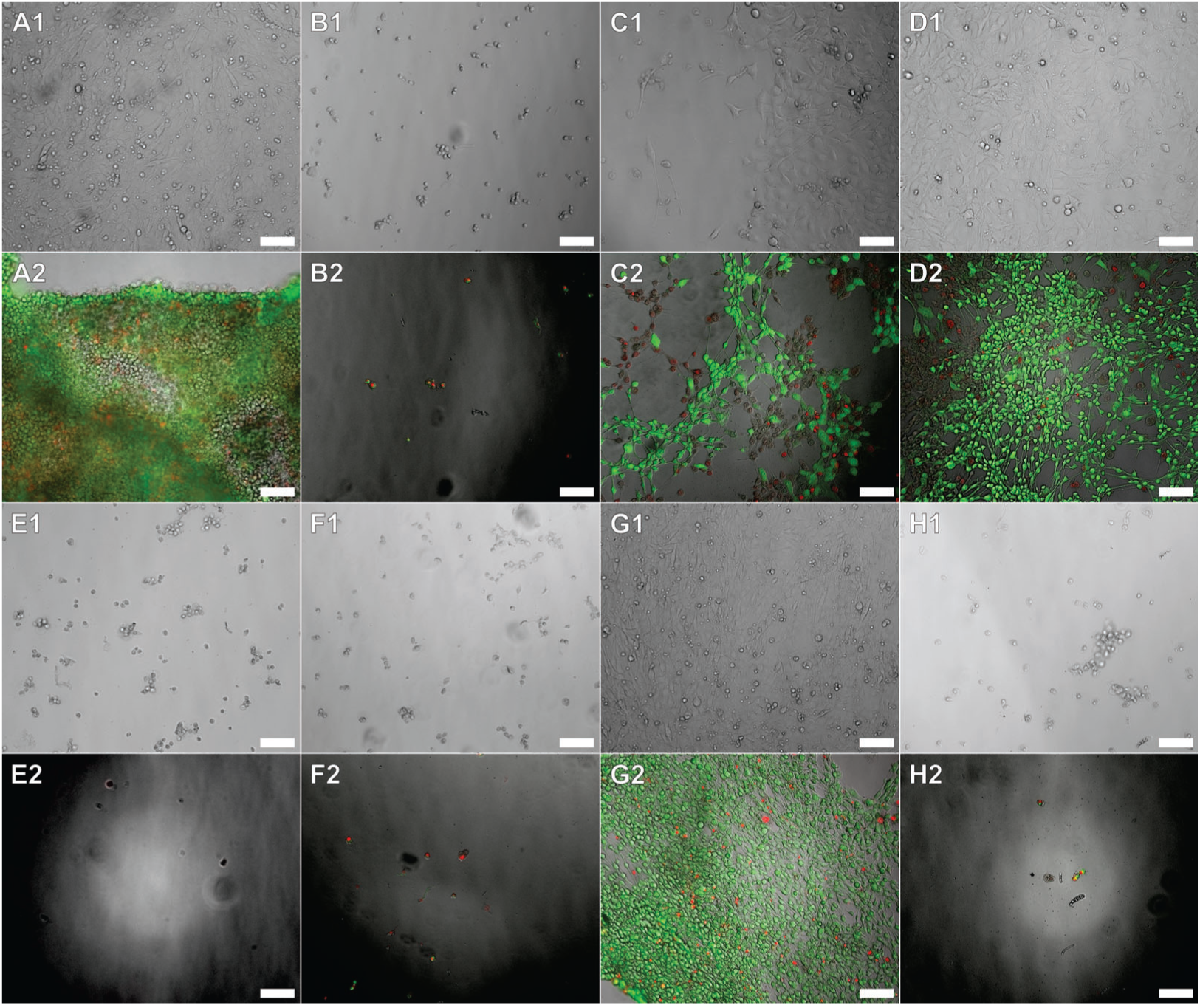
Brightfield (1) and fluorescence (2) microscopy images of cells that were exposed to autoclaved samples. The brightfield-only images (1) were captured prior to PBS washing and dye loading. (2) are brightfield images overlaid with calcein AM (green) and ethidium homodimer-1 (red) fluorescence images of cells. The panels represent cells exposed to the following autoclaved samples: (A) PDMS; (B) Phrozen AquaGray 8K; (C) Liqcreate Bio-Med Clear; (D) Asiga DentaGUIDE; (E) 3Dresyns Bioflex A10 MB IPA-post-treatment; (F) 3Dresyns Bioflex A10 MB UNW2-post-treatment; (G) Formlabs Silicone 40A IPA-post-treatment; (H) Asiga DentaGUM. All scale bars are 100 μm.

**Figure 6:**
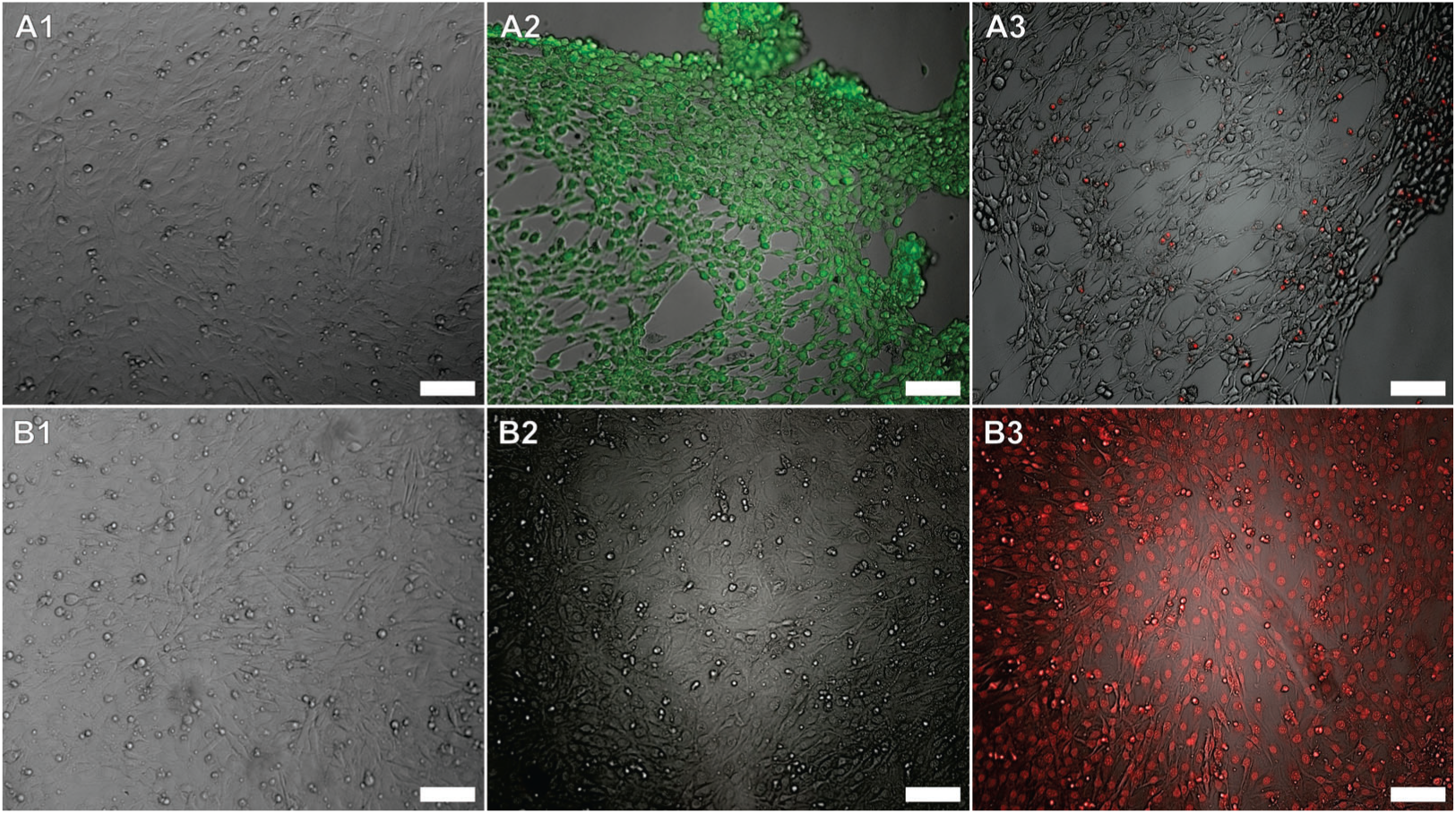
Brightfield and fluorescence microscopy images of cells that were not exposed to any resin samples. These controls were used for the biocompatibility assessment of the Formlabs Silicone 40A post-treated with IPA/BuOAc. (A) represent the blank (live cell) controls. (B) represent the positive (dead cell) controls, with (B1) illustrating the cells prior to incubation with 70% ethanol. (1) are brightfield-only images of the cells prior to loading of any dye. (2) are brightfield images overlaid with fluorescence images of cells loaded only with calcein AM (green). (3) are brightfield images overlaid with fluorescence images of cells loaded only with ethidium homodimer-1 (red). All scale bars are 100 μm.

**Figure 7:**
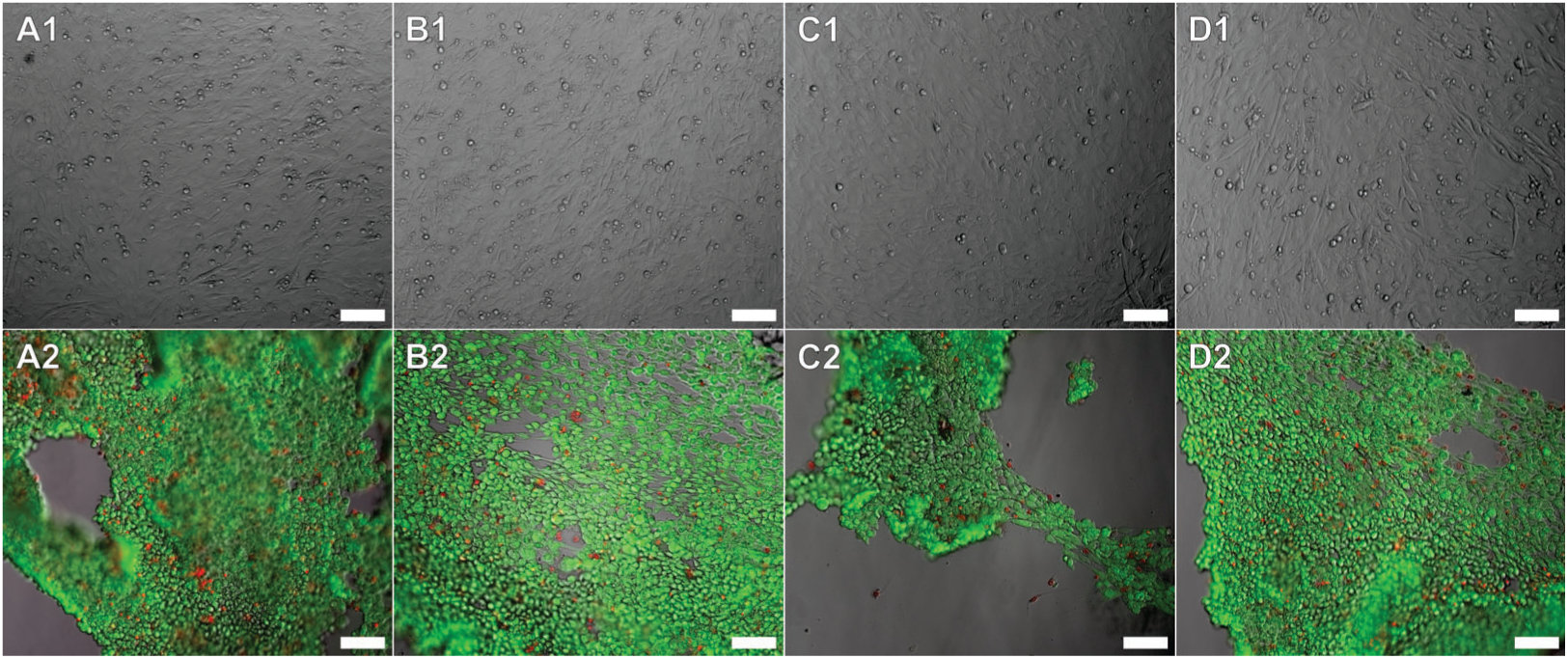
Brightfield (1) and fluorescence (2) microscopy images of cells exposed to the Formlabs Silicone 40A samples post-treated with IPA/BuOAc or PDMS. (1) are brightfield-only images of cells prior to dye loading. (2) are brightfield images overlaid with calcein AM (green) and ethidium homodimer-1 (red) fluorescence images of cells. The panels represent cells exposed to the following samples: (A) Ethanol/UV-Sterilized PDMS; (B) Ethanol/UV-Sterilized Formlabs Silicone 40A IPA/BuOAc; (C) Autoclaved PDMS; (D) Autoclaved Formlabs Silicone 40A IPA/BuOAc. All scale bars are 100 μm.

**Table 6:**
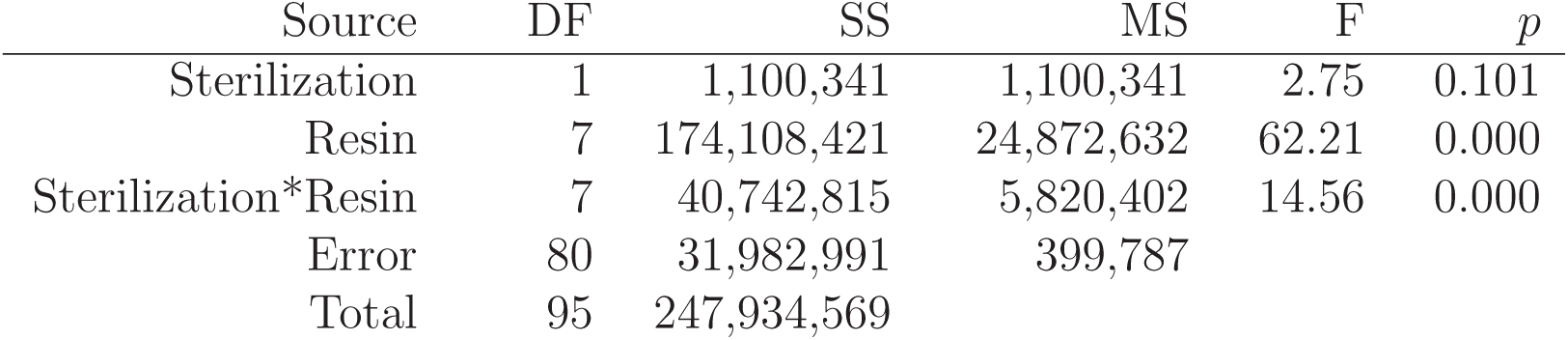
Analysis of variance (ANOVA) for assessing the effects on CalAM:EthD-1 responses for all resins except for the Formlabs Silicone 40A with the IPA/BuOAc posttreatment. A PDMS negative control was also included in this group. The “Sterilization*Resin” factor represents the interaction between the two factors. DF: degrees of freedom; SS: adjusted sum of squares; MS: adjusted mean squares; F: ANOVA test statistic; *p*: probability value.

**Table 7:** Analysis of variance (ANOVA) for assessing the effects on CalAM:EthD-1 responses for only Formlabs Silicone 40A with the IPA/BuOAc post-treatment and PDMS (negative control). The “Sterilization*Resin” factor represents the interaction between the two factors. DF: degrees of freedom; SS: adjusted sum of squares; MS: adjusted mean squares; F: ANOVA test statistic; *p*: probability value.

| Source | DF | SS | MS | F | $p$ |
| --- | --- | --- | --- | --- | --- |
| Sterilization | 1 | 211,293 | 211,293 | 0.26 | 0.615 |
| Resin | 1 | 1,685,275 | 1,685,275 | 2.09 | 0.164 |
| Sterilization*Resin | 1 | 4,134 | 4,134 | 0.01 | 0.944 |
| Error | 20 | 16,159,580 | 807,979 |  |  |
| Total | 23 | 18,060,283 |  |  |  |

#### 3.1.1. Elastomeric Resin Cytotoxicity

*3Dresyns Bioflex A10 MB* Regardless of the post-print process with IPA or UNW2 and sterilization method, the 3Dresyns Bioflex A10 MB samples produced the lowest average CalAM:EthD-1 response, which was significantly different from the response to PDMS (Table 8, Figure 8, and Figure 9). This is complemented with a qualitative assessment of the cultures exposed to 3Dresyns Bioflex A10 MB, in which the cells were rounded and detached from the surface prior to the PBS washes and loading of the dyes (Figure 4E1-F1 and Figure 5E1-F1). In the quantitative, post-hoc Bonferroni pairwise analysis, the only exception were comparisons to ethanol/UV-sterilized PDMS where the CalAM:EthD-1 responses were not significantly different (Table 8), which was unexpected due to the healthy morphology exhibited in the cultures exposed to PDMS prior to the PBS washes (see Section 3.1.3 for an expanded discussion). Taking into account the qualitative morphology prior to the washes and dyes and the lower, significantly different response from the response to autoclaved PDMS, 3Dresyns Bioflex A10 MB was assumed to have a negative impact on the viability of the C2C12 cultures. However, the manufacturer does note that this monomer-based formulation has a lower biocompatibility than their monomer-free formulation. Monomers in resins have been found to be cytotoxic (Wiertelak-Makal-a et al., 2023), especially if the print structures were not fully cured to polymerize any free monomers. Furthermore, the manufacturer suggests that the use of a low-power LCD resin printer, such as the Phrozen Sonic Mini 8K used in this study, may result in high concentrations of uncured monomers (3Dresyns, 2024), which may contribute to the low CalAM:EthD-1 responses found in this study.

**Figure 8:**
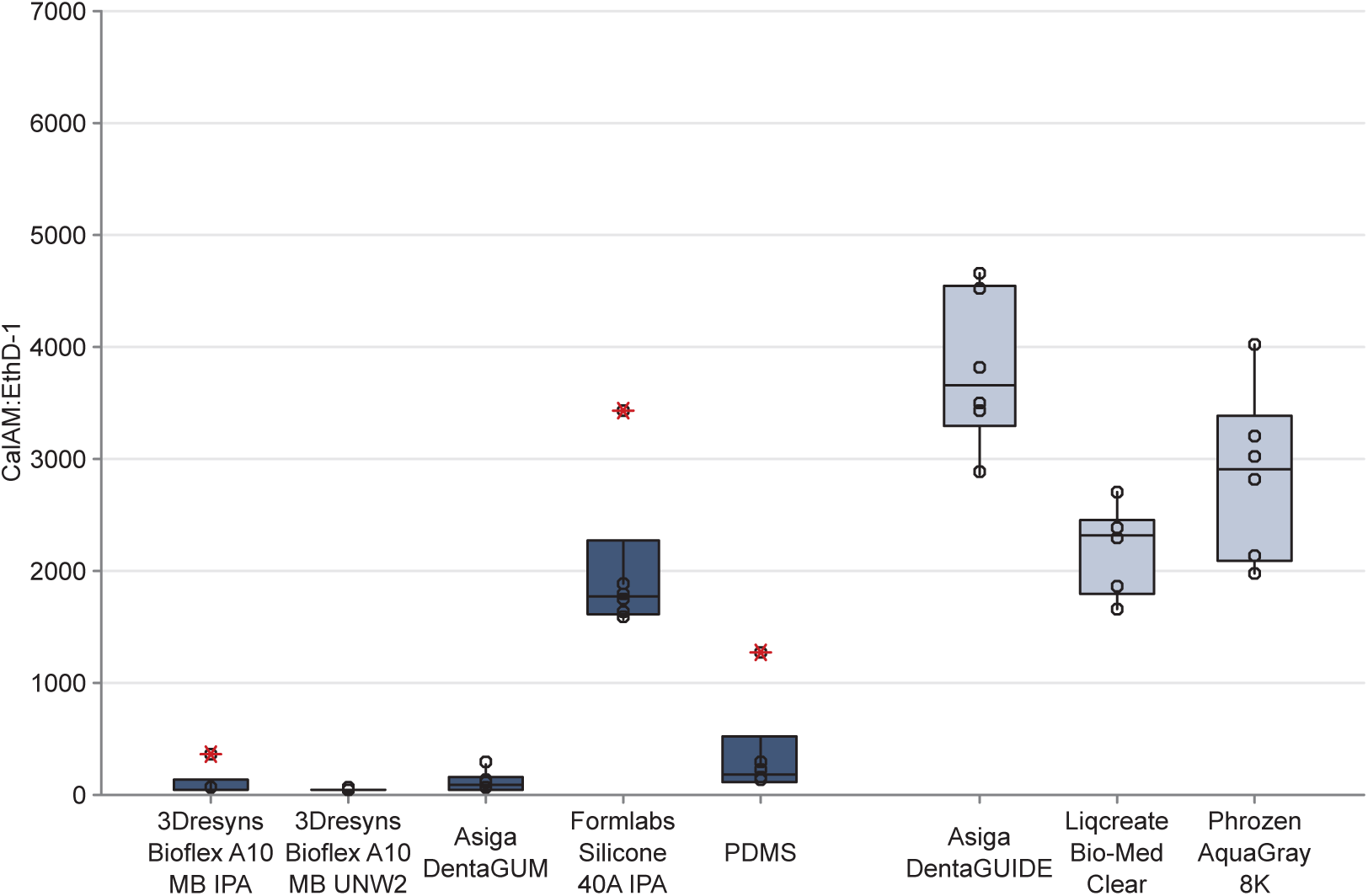
The CalAM:EthD-1 responses represented as boxplots of the ethanol/UV-sterilized rigid (dark blue-gray) and elastomeric (light blue-gray) resin samples, excluding the Formlabs Silicone 40A post-treated with IPA/BuOAc. Open circles (◦) represent individual data points. The red asterisks (*) denote outliers. The upper and lower boundaries of the box represent the first and third quartiles. Whiskers extend from the interquartile boxes to the minimum and maximum values, excluding outliers.

**Figure 9:**
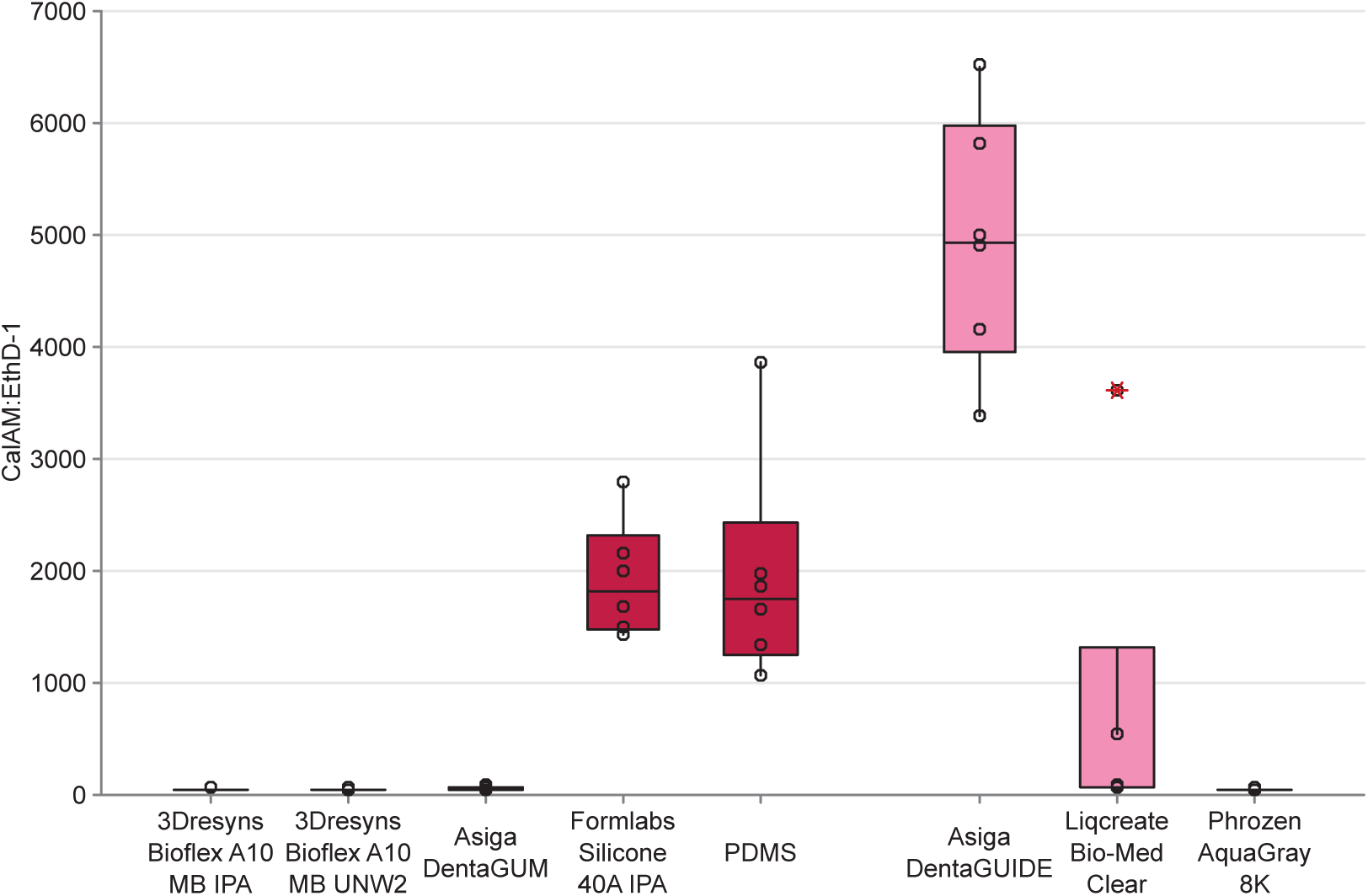
The CalAM:EthD-1 responses represented as boxplots of the autoclaved rigid (dark pink) and elastomeric (light pink) resin samples, excluding the Formlabs Silicone 40A post-treated with IPA/BuOAc. Open circles (◦) represent individual data points. The red asterisks (*) denote outliers. The upper and lower boundaries of the box represent the first and third quartiles. Whiskers extend from the interquartile boxes to the minimum and maximum values, excluding outliers.

**Table 8:** Post-hoc Bonferroni pairwise comparisons for the CalAM:EthD-1 responses between each type of factor per source, as listed in the ANOVA for all of the resins except for the IPA/BuOAc-post-treated Formlabs Silicone 40A (Table 6). N is the number of samples per factor. StD is the standard deviation. Factors that share a Group letter per type of source are not significantly different.

| **Source (bold)**<br>Factor | | N | Mean ± StD | Group |
| --- | --- | --- | --- | --- |
| **Sterilization** |  |  |  |  |
| Ethanol/UV | | 48 | 1,421 ± 1,460 | A |
| Autoclave | | 48 | 1,207 ± 1,766 | A |
| **Resin** |  |  |  |  |
| Asiga DentaGUIDE | | 12 | 4,362 ± 1,072 | A |
| Formlabs Silicone 40A IPA | | 12 | 1,948 ± 585 | B |
| Liqcreate Bio-Med Clear | | 12 | 1,459 ± 1,254 | B |
| Phrozen AquaGray 8K | | 12 | 1,435 ± 1,551 | B |
| PDMS | | 12 | 1,143 ± 1,108 | B |
| Asiga DentaGUM | | 12 | 72.7 ± 65.8 | C |
| 3Dresyns Bioflex A10 MB IPA | | 12 | 64.4 ± 83.9 | C |
| 3Dresyns Bioflex A10 MB UNW2 | | 12 | 29.6 ± 5.86 | C |
| **Steril.** | **Resin** |  |  |  |
| Autoclave | DentaGUIDE | 6 | 4,941 ± 1,122 | A |
| Ethanol/UV | DentaGUIDE | 6 | 3,783 ± 683 | A B |
| Ethanol/UV | AquaGray 8K | 6 | 2,839 ± 749 | B C |
| Ethanol/UV | Bio-Med Clear | 6 | 2,194 ± 384 | C |
| Ethanol/UV | Silicone 40A IPA | 6 | 1,991 ± 703 | C D |
| Autoclave | PDMS | 6 | 1,940 ± 989 | C D |
| Autoclave | Silicone 40A IPA | 6 | 1,906 ± 505 | C D |
| Autoclave | Bio-Med Clear | 6 | 724 ± 1420 | D E |
| Ethanol/UV | PDMS | 6 | 345 ± 446 | E |
| Ethanol/UV | DentaGUM | 6 | 102 ± 86.0 | E |
| Ethanol/UV | Bioflex A10 MB IPA | 6 | 87.6 ± 119 | E |
| Autoclave | DentaGUM | 6 | 43.77 ± 9.38 | E |
| Autoclave | Bioflex A10 MB IPA | 6 | 41.2 ± 4.74 | E |
| Autoclave | AquaGray 8K | 6 | 30.6 ± 3.62 | E |
| Autoclave | Bioflex A10 MB UNW2 | 6 | 30.4 ± 4.00 | E |
| Ethanol/UV | Bioflex A10 MB UNW2 | 6 | 28.8 ± 7.63 | E |

##### Formlabs Silicone 40A

When compared to the PDMS negative controls, Formlabs Silicone 40A minimally impacted the CalAM:EthD-1 response. Prior to the loading of the dyes, the cultures exposed to Formlabs Silicone 40A, under any sterilization and post-treatment conditions (Figure 4G1, Figure 5G1, Figure 7B1,D1), appeared qualitatively similar to the cultures exposed to PDMS, under any sterilization condition (Figure 4A1, Figure 5A1, Figure 7A1,C1) and to cultures not exposed to any resin samples (Figures 3A1, 6A1). This qualitative observation aligned with the quantitative analysis of CalAM:EthD-1 responses where the post-hoc Bonferroni pairwise comparisons indicated that responses to Formlabs Silicone 40A were not significantly different from the responses to PDMS (Table 8, Table 9, Figure 9, Figure 8, Figure 10). Currently, the manufacturer lists this resin as pending for their internal cytotoxicity assessment (Formlabs, 2024d), but these results demonstrate potential biocompatibility with C2C12 myoblasts.

**Figure 10:**
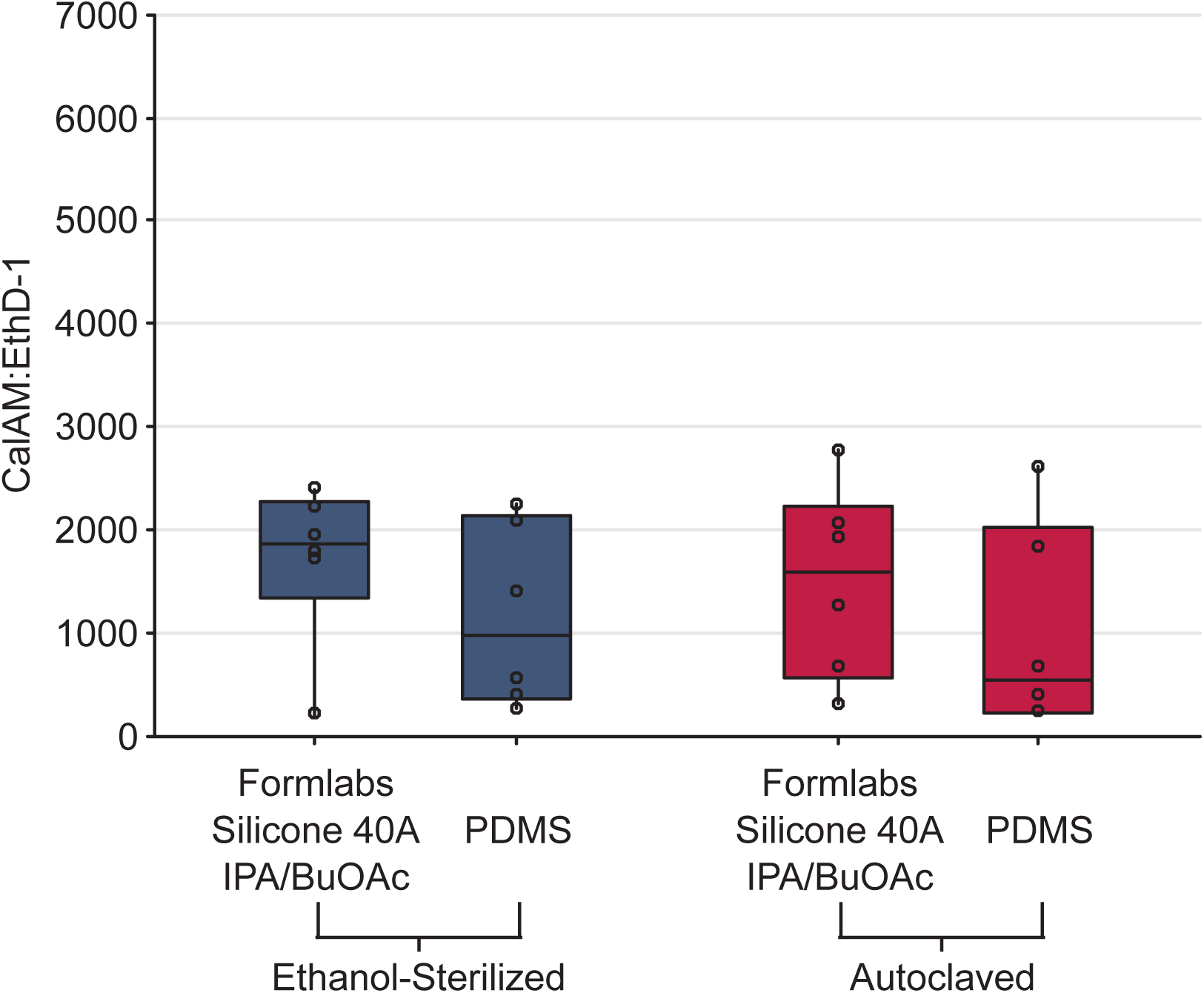
The CalAM:EthD-1 responses represented as boxplots of the ethanol/UV-sterilized (dark blue-gray) and autoclaved (dark pink) Formlabs Silicone 40A post-treated with IPA/BuOAc samples and its associated PDMS negative control samples. Open circles (◦) represent individual data points. None of the individual data points were outliers. The upper and lower boundaries of the box represent the first and third quartiles. Whiskers extend from the interquartile boxes to the minimum and maximum values, excluding outliers.

**Table 9:** Post-hoc Bonferroni pairwise comparisons for the CalAM:EthD-1 responses between each type of factor per source, as listed in the ANOVA for the analysis of IPA/BuOAc-post-treated Formlabs Silicone 40A (Table 7). N is the number of samples per factor. StD is the standard deviation. Factors that share a Group letter per type of source are not significantly different.

| **Source (bold)**<br>Factor | | N | Mean ± StD | Group |
| --- | --- | --- | --- | --- |
| **Sterilization** |  |  |  |  |
| Ethanol/UV | | 12 | 1,429 ± 844 | A |
| Autoclave | | 12 | 1,242 ± 954 | A |
| **Resin** |  |  |  |  |
| Formlabs Silicone 40A IPA/BuOAc | | 12 | 1,600 ± 825 | A |
| PDMS | | 12 | 1,071 ± 899 | A |
| **Sterilization** | **Resin** |  |  |  |
| Ethanol/UV | Silicone 40A IPA/BuOAc | 6 | 1,707 ± 780 | A |
| Autoclave | Silicone 40A IPA/BuOAc | 6 | 1,494 ± 928 | A |
| Ethanol/UV | PDMS | 6 | 1,151 ± 879 | A |
| Autoclave | PDMS | 6 | 990 ± 995 | A |

##### Asiga DentaGUM

When compared to the PDMS negative controls, Asiga DentaGUM caused a significant change in CalAM:EthD-1 response. For both the ethanol/UV-sterilized and autoclaved techniques, there were very few cells observed in the DentaGUM cultures, even before the PBS washes and dye loading (Figure 4H1, Figure 5H1), whereas the PDMS cultures were observed to have *>*90% confluency (Figure 4A1, Figure 5A). Overall, this was in alignment with the quantitative, post-hoc Bonferroni pairwise analysis in which DentaGUM was significantly different from PDMS (Table 8). The only exception was if the sterilization technique was in consideration with the resin type in which the CalAM:EthD-1 responses to DentaGUM under either sterilization method were not significantly different from ethanol/UV-sterilized PDMS (Table 8, Figure 8), which was unexpected (see Section 3.1.3 for an expanded discussion). However, since DentaGUM under either sterilization technique was still significantly different from autoclaved PDMS and their cultures qualitatively appeared different from the PDMS cultures, DentaGUM was inferred to negatively impact C2C12 viability. The cytotoxicity of Asiga DentaGUM is not surprising as the manufacturer’s intended use for this resin is as a model of the gingiva (Asiga, 2024a) as opposed to a medical device in direct contact with patients.

#### 3.1.2. Rigid Resin Cytotoxicity

##### Asiga DentaGUIDE

Asiga DentaGUIDE did not negatively impact cell viability. Prior to the PBS washes and loading of the dye, the cultures exposed to DentaGUIDE (Figure 4D1 and Figure 5D1) were qualitatively indistinguishable from cultures exposed to PDMS (Figure 4A1 and Figure 5A1) and cultures not exposed to any resins (Figure 3A1). Unexpectedly, the quantitative assessment identified a significant difference between DentaGUIDE and PDMS (Table 8, Figure 8, Figure 9). However, in the quantitative assessment, the average CalAM:EthD-1 response to DentaGUIDE was consistently larger than the average response to PDMS. The larger ratio suggests a larger population of living cells or much fewer dead cells present in the cultures. Therefore, the quantitative analysis suggests that DentaGUIDE did not negatively impact the viability of C2C12, which aligns with the manufacturer’s report of DentaGUIDE’s Class I biocompatibility certification (Asiga, 2024a).

##### Liqcreate Bio-Med Clear

Overall, Liqcreate Bio-Med Clear samples did not significantly affect the CalAM:EthD-1 response. Prior to the PBS washes and loading of the dye, the cultures exposed to the Bio-Med Clear samples (Figure 4C1 and Figure 5C1) were qualitatively similar to the PDMS negative control cultures (Figure 4A1 and Figure 5A1). This was aligned with the quantitative analyses where there was not a significant difference between Liqcreate Bio-Med Clear and PDMS (Table 8, Figure 8, Figure 9). The minimal C2C12 cytotoxicity observed for Liqcreate Bio-Med Clear complements the manufacturer’s assessment of biocompatibility using ISO 10993-5:2009 (Liqcreate, 2024). In addition, the manufacturer noted that Bio-Med Clear can be autoclave sterilized and generally cleaned with “commonly used disinfectants” (Liqcreate, 2024). Furthermore, the ethanol/UV-sterilized Bio-Med Clear samples produced a larger CalAM:EthD-1 response, which was significantly different from the autoclaved Bio-Med Clear samples. Thus, this study’s results indicate that the use of 70% ethanol submersion with UV exposure is also an acceptable sterilization technique.

##### Phrozen AquaGray 8K

After sterilizing with ethanol/UV, Phrozen AquaGray 8K did not have significant impacts on C2C12 viability (Figure 8B, Table 8). Autoclaved AquaGray 8K samples caused a significantly different CalAM:EthD-1 response (Figure 9B, Table 8). Qualitatively, cultures exposed to ethanol/UV-sterilized AquaGray 8K samples exhibited several C2C12 cells with a typical morphology (Figure 4B1), while the cultures exposed to the autoclaved samples had a more rounded morphology (Figure 5B1), which is indicative of poor cell health. These qualitative observations complemented the quantitative assessment using the post-hoc Bonferroni pairwise comparison where the ethanol/UV-sterilized AquaGray 8K were not significantly different from the autoclaved PDMS controls (Table 8). Ethanol/UV-sterilized AquaGray 8K was significantly different from ethanol/UV-sterilized PDMS, but similar to the discussion of the DentaGUIDE results, the average CalAM:EthD-1 response was larger than the response due to ethanol/UV-sterilized PDMS, which would indicate no negative impacts. In contrast, the autoclaved AquaGray 8K was significantly different from the autoclaved PDMS (smaller CalAM:EthD-1 ratio) and not significantly different from the ethanol/UV-sterilized PDMS. Similar to the interpretations of the DentaGUM results, since the autoclaved Aquagray 8K exhibited qualitatively poor morphologies in comparison to both PDMS cultures and was significantly different from the autoclaved PDMS, autoclaved AquaGray 8K was assumed to negatively impact the C2C12 viability. The cytotoxicity of autoclaved AquaGray was not surprising since, unlike the other resins in this study, Phrozen AquaGray 8K is not intended for medical or biological applications. Under autoclave conditions, the temperature reaches 121 °C, which may impact the AquaGray 8K samples which the manufacturer reported a heat deflection temperature of 60.3 - 62.7 °C under a 0.45 MPa load. Although the AquaGray 8K cytotoxicity samples were not noticeably distorted after autoclaving (see Section 3.3 for detailed analyses), the high heat may have impacted the samples to release leachants into the culture media. Unlike the high heat of the autoclave procedure, the ethanol/UV-sterilization procedure is similar to the normal post-printing process involving alcohol and UV post-cure. Therefore, ethanol/UV-sterilized AquaGray 8K could potentially be suitable for use with C2C12 cultures.

#### 3.1.3. Cytotoxicity Limitations

While the use of calcein AM and ethidium homodimer-1 fluorescent dyes in a direct contact assessment provides insight into the potential cytotoxicity of these resin samples, there are several limitations to the experimental procedures. Notably, the PBS washes and application of the dyes impacted the culture morphology, which was even observed in the cell-only (blank) cultures. Before the washes and dye, the C2C12 exhibited a relatively flat cell sheet with an elongated morphology typical of healthy C2C12 (Figure 3A1 and Figure 6A1). However, after these procedures, the C2C12 morphology became more rounded and began to detach from the culture surface (Figure 3A2-3 and Figure 6A2-3). This was not observed in the positive control in which the C2C12 were intentionally killed using ethanol, which fixed the cells to the culture surface and preserved their morphology (Figure 3B and Figure 6B). However, despite the impacts of the washes and the dye, the vivid fluorescence of calcein AM with the minimal fluorescence of ethidium homodimer-1 in the blank controls (Figure 3A2-3 and Figure 6A2-3) indicate that the dyes are still functional and indicate the cell viability and toxicity as intended. Furthermore, since the calcein AM and ethidium homodimer-1 assessment was completed as an endpoint analysis, any long-term impacts on the cellular health from these dyes will not affect the results of this study. In addition to the dyes, the PBS wash steps may also contribute to the removal of dead cells, especially in the cultures that already exhibited cells with a more rounded morphology (i.e. ethanol/UV-sterilized DentaGUM (Figure 4H1), autoclaved DentaGUM (Figure 5H1), autoclaved AquaGray 8K (Figure 5B1)). For C2C12 cells, the observation of a rounded morphology is indicative of dead and dying cells that were likely already detached from the culture surface. If the cells were already detached from the culture surface prior to the PBS washes, they likely were removed from the culture before the dye application, which could account for the post-dye reduction of cells in these cultures (Figure 4H2) and Figure 5H2,B2). Cultures exhibiting a healthy morphology are more likely to remain in the well due to more robust cell adhesion, which is supported by the live cell (blank) control cultures, which did not demonstrate the rounded morphology before dye application and largely remained in the wells after the PBS washes and dye application (Figure 3A, Figure 6A). Due to the much larger number of cells in these healthy cultures, any dead cell losses are more difficult to ascertain. However, assessing cytotoxicity via the ratio of the fluorescence intensities of calcein AM and ethidium homodimer-1 (CalAM:EthD-1), rather than directly comparing the individual fluorescence, provided a more holistic assessment that accounted for the expected reduction of dead cell (ethidium homodimer-1) fluorescence due to cell detachment before dye exposure. Furthermore, when using calcein AM and ethidium homodimer-1 as the method for determining cell viability, analyzing the ratio of calcein AM to ethidium homodimer is more appropriate for analysis than using individual fluorescence intensities to quantify living and dead cells (Gantenbein-Ritter et al., 2008).

To further investigate the impacts of these resins on cellular health, additional cell viability assessments can be explored. Calcein AM is a live cell marker by tagging intracellular esterases typically only present in living cells whereas ethidium homodimer-1 binds to nucleic acids, but is only permeable across damaged plasma membranes that are typical of dead cells (Probes, 2005). However, cell health may be impacted without causing cell death. Other assays that investigate metabolism or proliferation, such as tetrazolium-based assays (*e.g.* MTT) (Riss et al., 2004) or 5-ethynyl-2’-deoxyuridine (EdU) (Salic and Mitchison, 2008) respectively, could provide valuable cellular health information as C2C12 are highly active and proliferative as an immortal cell line. Furthermore, although this study provides insight into potential resins, the current work is limited to effects in a 2D culture environment. To continue assessing the applicability of these resins to biohybrid actuators, future work should include cell viability and cytotoxicity assays in 3D cultures, in which calcein AM and ethidium homodimer-1 can still be used.

It is also important to note that while the viability assessment reported here provides insight into the effect of the as-manufactured samples, it does not reveal the cause of decreased viability. The low-power nature of the equipment used from manufacturing and curing, may result in undercured samples, and residual uncured monomer is likely to contribute to cytotoxicity. Future work is needed to assess the presence of leachants in the media and the percentage of uncured polymer remaining. As this work is beyond the scope of the studies completed here, our results should only be interpreted for use of these materials given the low-cost manufacturing equipment described in the methods.

In addition to the viability assessment, the use of a direct contact assessment also has limitations. In particular, to prevent any discrepancies in the fluorescence readings due to the potential autofluorescence of resins, all of the resin samples had to be removed prior to the PBS washes and loading of the dyes. The resins were carefully removed using forceps, but any movement of the resins could cause the samples to act as a cell scraper and detach a portion of the cells from the cell surface. In particular, this may explain the unexpectedly low CalAM:EthD-1 response to ethanol/UV-sterilized PDMS. The PDMS samples tended to float in the culture medium, whereas most of the other resin samples settled at the bottom. The floating phenomenon causes the sample to move in the culture medium during plate handling and sample removal, whereas samples settled at the bottom of the well were less likely to move. Therefore, floating samples could lead to multiple removal attempts and heighten the chances of the forceps or the moving sample scraping cells off of the culture surface prior to the wash steps, resulting in a potentially lower CalAM:EthD-1 response. In the future, this hypothesis could be verified by using an elution assay, which uses culture media conditioned through long-term exposure to samples to indirectly test for biocompatibility (ISO 10993-5:2009 (International Organization for Standardization, 2009) and ISO 10993-12:2021 (International Organization for Standardization, 2021)). However, elution studies would only assess the effects of potential leachants, which might not adequately reproduce biohybrid environments where cells are in direct contact with the material. Therefore, indirect tests should still be done in conjunction with direct contact assessments to more closely replicate the environmental conditions cells would experience in biohybrid systems.

### 3.2. Mechanical Behavior

#### 3.2.1. Elastomeric Resin Mechanical Properties

3D-printed samples of each elastomeric resin treated with each sterilization technique were subjected to tensile and compression mechanical tests. Stress-strain data were calculated relative to each specimen’s rest geometry, and all curves can be found in Appendix B. From these stress-strain curves, scalar quantities were calculated to allow for more direct comparisons between the groups. Specifically, we chose to examine the Young’s Modulus, stress value at 50% strain, and stress value at 100% strain to compare with metrics commonly reported by manufacturers of these resins (Table 1). These metrics were measurable for all samples except for the 3Dresyns Bioflex A10 MB groups. These groups saw some samples fail before 50% strain and many samples fail before 100% strain. For these samples, no values are reported for these metrics. In addition to these metrics, all Asiga Dentagum samples failed during their tensile tests, and from these curves, the Ultimate Tensile Strength and elongation at break were measured. Only four Formlabs Silicone 40A samples across both post-treatment groups and all sterilization groups failed. Because of this, these failure characteristics are not reported for Formlabs Silicone 40A groups.

##### 3Dresyns Bioflex A10 MB, IPA and UNW2 post treatments

Neither 3Dresyns Bioflex A10 MB post-treatment groups were heavily affected by the different sterilization techniques, with no metrics seeing a statistically significant difference to the nonsterile control group (Fig. 11, Column 1, 2, Fig. 12, Column 1, 2). For the IPA post-treatment group, both sterilization protocols only had a mild impact on the Young’s Modulus (autoclave: +3.3 ± 0.3%, ethanol/UV: −3.00 ± 0.19% relative to the nonsterile group). The UNW2 group saw a similarly mild change in the Young’s modulus for ethanol/UV sterilization (+4.5 ± 0.2%), but a larger change was seen in the autoclave group, with a decrease of 8.0 ± 0.4%. Small changes were also seen in the stress at 50% strain metric for both post-treatments (IPA, autoclave: −0.49 ± 0.03%, IPA, ethanol/UV: −3.0 ± 0.2%, UNW2, ethanol/UV: −4.2 ± 0.3%). However, for the UNW2 autoclaved group, not all samples survived to 50% strain, so this metric could not be calculated for all samples. We also could not examine the Stress at 100% because only the IPA post-treated, ethanol/UV sterilized group had all samples reach 100% strain (across all other groups, only 3 other samples failed after 100% strain).

**Figure 11:**
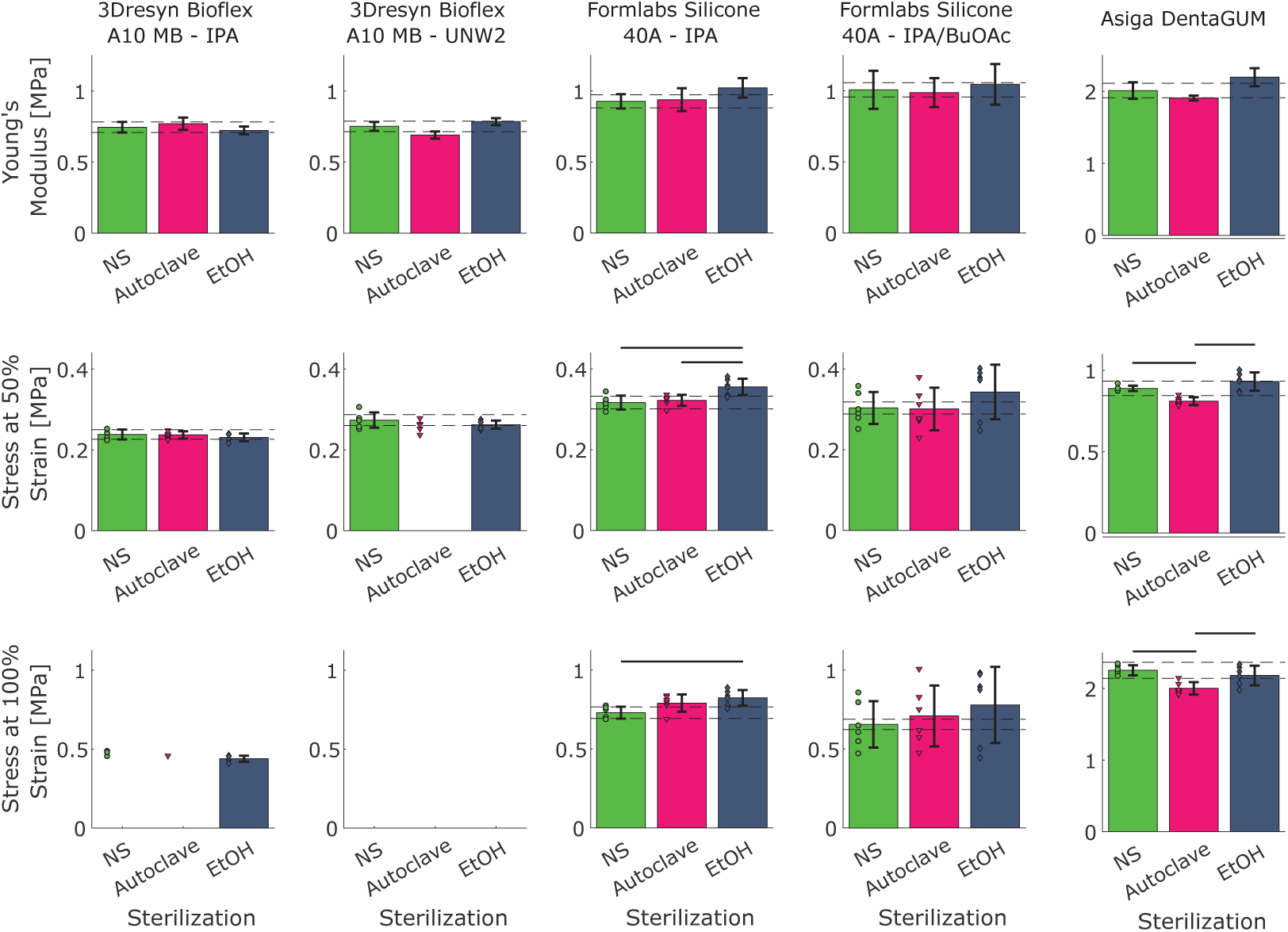
Summary metrics for elastomeric resins. The Young’s Modulus [MPa] (top row, calculated from model parameters as *E* = 6*C*_1_, see Table 10), Stress at 50% strain (middle row), and Stress at 100% strain (bottom row) are reported for the five elastomeric resin groups and for each sterilization technique. Bar height indicates mean value and error bars show ± standard deviation. For Stress at 50% strain and Stress at 100% strain, marks show individual data points, with one data point per specimen tested. Missing bar indicate that not all specimen in that group achieve the level of strain required to take that measurement (this only occurred in the two 3Dresyns Bioflex groups). Note that the scales for Asiga Dentagum is higher than the rest of the charts. Otherwise, scales are consistent across each row. Horizontal dashed lines indicate ±5% of the corresponding nonsterile group mean. Solid lines above bars indicate statistical difference in means at the 5% level.

**Figure 12:**
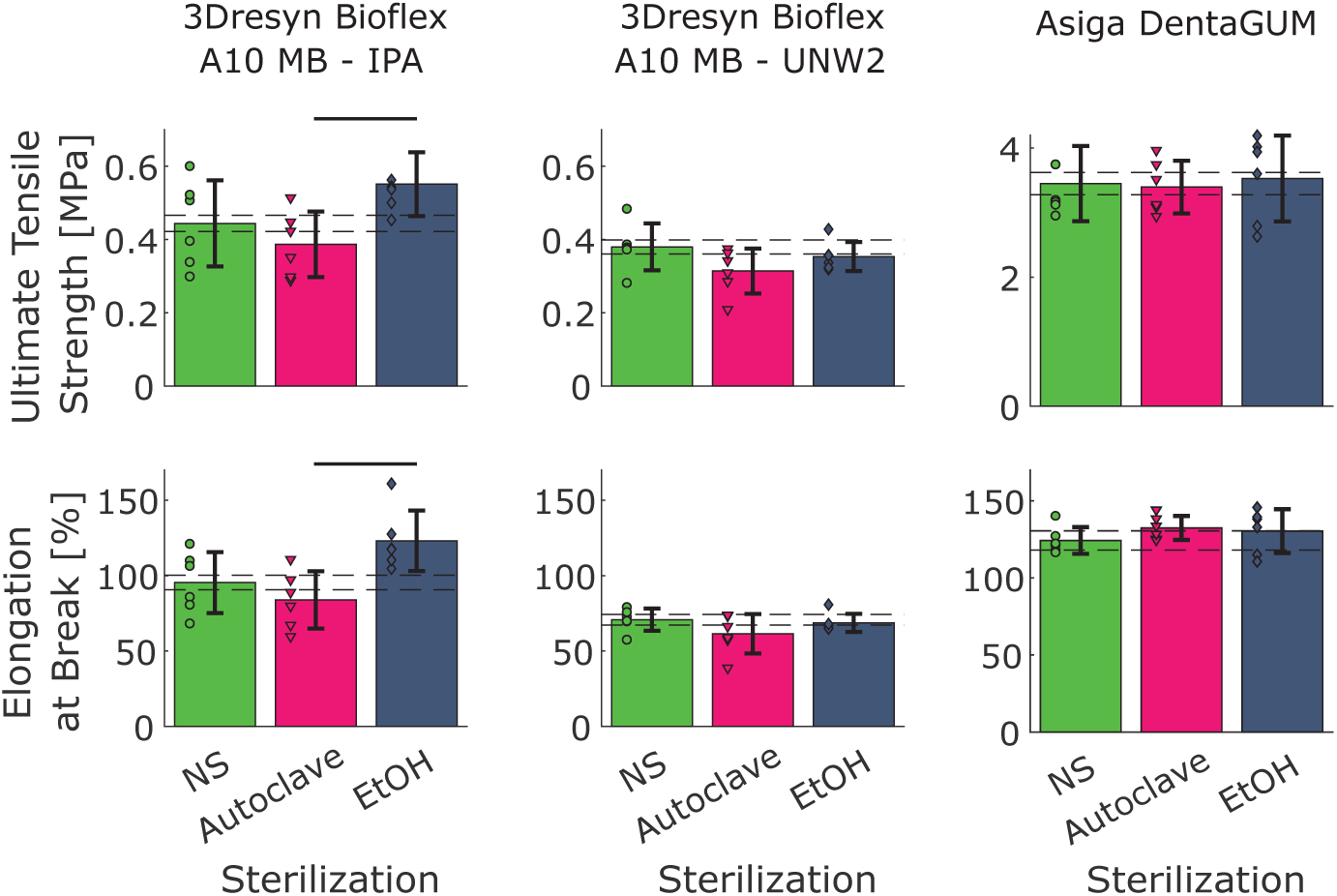
Failure characteristics for elastomeric resins. Ultimate Tensile Strength [MPa] (top row) and elongation at break [%] (bottom row) are reported for the three elastomeric resins that failed within the test regime – 3Dresyns Bioflex IPA, 3Dresyns Bioflex UNW2, and Asiga Dentagum. Neither FormLabs Silicone groups reliably failed within the test regime. Bar height indicates mean value and error bars show ± standard deviation. Circular marks show individual data points, with one data point per specimen tested. Note that the scales for Asiga Dentagum are higher than the rest of the charts. Otherwise, scales are consistant across each row. Horizontal dashed lines indicate ±5% of the corresponding nonsterile group. Solid lines above bars indicate statistical difference in means at the 5% level.

**Figure 13:**
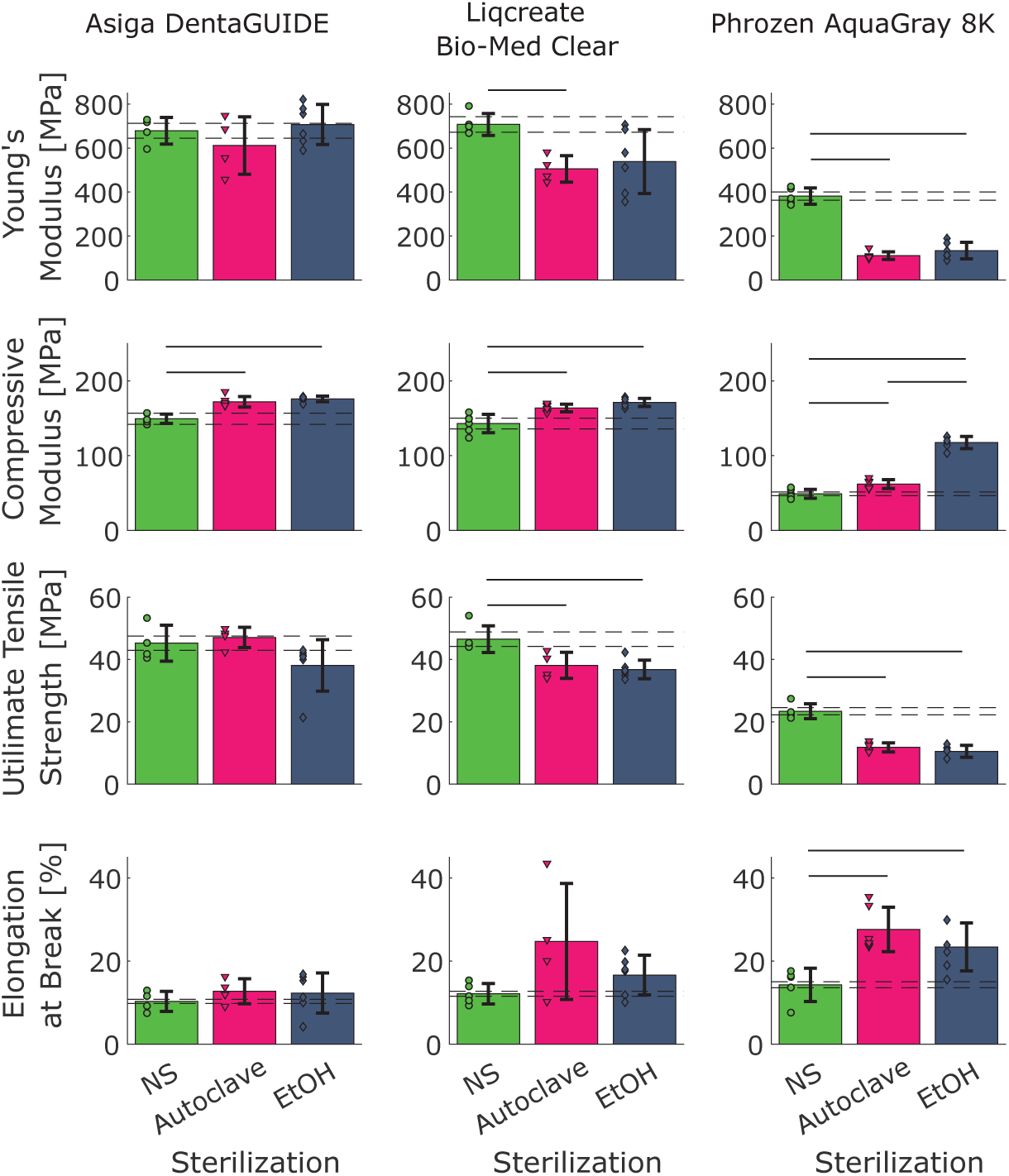
Summary Metrics for Rigid Resins. The Young’s Modulus [MPa] (top row), Compressive Modulus [MPa] (second row), Ultimate Tensile Strength [MPa] (third row), and Elongation at Break [%] (bottom row) are reported for the three rigid resin groups and for each sterilization technique. Bar height indicates mean value and error bars show ± standard deviation. Circular marks show individual data points, with one data point per specimen tested. Horizontal dashed lines indicate ±5% of the corresponding nonsterile group. Solid lines above bars indicate statistical difference in means at the 5% level.

**Table 10:** Elastomeric resin Yeoh Model parameters resulting from bootstrapped model fitting. Values are reported as mean ± standard deviation.

| Resin | Sterilization | $C_1$ [MPa] | $C_2$ [MPa] | $C_3$ [MPa] | $R^2$ |
| --- | --- | --- | --- | --- | --- |
| 3Dresyns<br>Bioflex MB IPA | Nonsterile | $0.124 \pm 0.006$ | $-1 \pm 8 \times 10^{-3}$ | $0.3 \pm 1.5 \times 10^{-3}$ | 0.990 |
| | Autoclave | $0.128 \pm 0.007$ | $-1 \pm 6 \times 10^{-3}$ | $3 \pm 5 \times 10^{-4}$ | 0.981 |
| | Ethanol/UV | $0.120 \pm 0.005$ | $3 \pm 3 \times 10^{-3}$ | $1 \pm 6 \times 10^{-4}$ | 0.986 |
| 3Dresyns<br>Bioflex MB<br>UNW2 | Nonsterile | $0.126 \pm 0.006$ | $1.5 \pm 1.4 \times 10^{-3}$ | $1.8 \pm 0.6 \times 10^{-3}$ | 0.996 |
| | Autoclave | $0.115 \pm 0.004$ | $3 \pm 4 \times 10^{-3}$ | $1.9 \pm 1.2 \times 10^{-3}$ | 0.997 |
| | Ethanol/UV | $0.131 \pm 0.004$ | $-1 \pm 3 \times 10^{-3}$ | $1 \pm 0.9 \times 10^{-3}$ | 0.995 |
| Asiga<br>Dentagum | Nonsterile | $0.335 \pm 0.019$ | $0.062 \pm 0.017$ | $5 \pm 4 \times 10^{-3}$ | 0.999 |
| | Autoclave | $0.318 \pm 0.006$ | $0.051 \pm 0.006$ | $4.3 \pm 0.5 \times 10^{-3}$ | 0.998 |
| | Ethanol/UV | $0.37 \pm 0.02$ | $0.051 \pm 0.016$ | $4.6 \pm 1.8 \times 10^{-3}$ | 0.995 |
| FormLabs<br>Silicone IPA | Nonsterile | $0.154 \pm 0.008$ | $9 \pm 6 \times 10^{-3}$ | $1.6 \pm 0.6 \times 10^{-3}$ | 0.995 |
| | Autoclave | $0.156 \pm 0.013$ | $0.010 \pm 0.009$ | $2.3 \pm 1.1 \times 10^{-3}$ | 0.989 |
| | Ethanol/UV | $0.170 \pm 0.011$ | $6 \pm 6 \times 10^{-3}$ | $3.3 \pm 1.7 \times 10^{-3}$ | 0.984 |
| FormLabs<br>Silicone Mix | Nonsterile | $0.17 \pm 0.02$ | $3 \pm 8 \times 10^{-3}$ | $1.6 \pm 0.7 \times 10^{-3}$ | 0.912 |
| | Autoclave | $0.165 \pm 0.017$ | $3 \pm 1 \times 10^{-3}$ | $1.8 \pm 0.8 \times 10^{-3}$ | 0.918 |
| | Ethanol/UV | $0.17 \pm 0.02$ | $0.011 \pm 0.013$ | $1.4 \pm 1.0 \times 10^{-3}$ | 0.860 |

These materials begin to show larger differences in their failure characteristics (Fig. 12, Column 1, 2), though only the IPA+autoclave and IPA+ethanol/UV groups were statistically different from each other. For both post-treatments, the autoclave sterilization led to a decrease in the mean value of both the Ultimate Tensile Strength (IPA: 13 ± 5%, UNW2: 17 ± 4%) and the elongation at break (IPA: 12 ± 4%, UNW2: 13 ± 3%). However, these differences were not statistically significant because of the variance observed. Conversely, ethanol/UV sterilization led to an increase in the UTS and elongation at break for the IPA post-treated group (UTS: 24 ± 7%, elongation at break: 29±8%), while for UNW2 group, decreases were observed in both metrics (UTS: 7±1%, elongation at break: 3.0±0.4%). Overall, neither post-treatment group was statistically significantly affected by the sterilization protocol. However, for the changes that were observed, the IPA posttreatment had mild impacts on elastic properties and improvements in failure properties (when sterilized with ethanol/UV), whereas the manufacturer-recommended post-treatment (UNW2) saw mild effects on elastic properties and worsening of failure properties for both sterilization techniques.

##### Formlabs Silicone 40A, IPA and IPA/BuOAc post treatments

For Formlabs Silicone 40A post-treated with IPA/BuOAc, no statistically significant differences were observed in any of the metrics measured as a function of sterilization technique (Fig. 11(a), Column 2). Conversely, when the parts were posttreated with only with IPA (Fig. 11(a), Columns 1), ethanol/UV sterilization led to a statistically higher Stress at 50% and 100% compared to the nonsterile group (no difference between autoclave and nonsterile groups). This mean difference was also greater than 5% of the nonsterile mean (12.2 ± 1.0% for Stress at 50% and 13.0 ± 1.0% for Stress at 100%). While it appears as though the manufacturer-recommended post-treatment performed better than the IPA-only treatment (no statistically significant differences in mean values were observed), there was a much larger observed variance in these metrics across all sterilization methods for the recommended post-treatment groups. This variance is detrimental both to the power of the statistical tests for the given sample size of the study and to the reproducibility of the components produced using this technique. In addition, while statistically significant differences exist in the ethanol/UV-sterilized group for the IPA-only post-treated group, no difference was seen in the autoclaved group. Autoclaving also led to only a 1.25 ± 0.13% increase in the Young’s modulus and a 1.75 ± 0.12% increase in the stress at 50%. Stress at 100% saw an increase of 8.4 ± 0.7%, but this was not statistically significant. Thus, for this resin, when it comes to maintaining mechanical properties relative to pre-sterilized conditions and creating reproducible samples, IPA-only posttreatment sterilized in the autoclave performed the best.

This conclusion can only be drawn for metrics within the elastic regime of this material. Because of the force and displacement limitations enforced by the experimental setup, we were unable to test these samples to failure. The max strains (150%-200%) and max stresses (1-3 MPa) are less than the manufacturer’s provided strain at break of 230% and Ultimate Tensile Strength of 5 MPa, providing a lower bound confirmation of these metrics (Fig. B.22, Fig. B.23, Table 3). However, the true failure characteristics could not be tested. That said, given the strains that this material was able to sustain (*>*150% for all samples in all groups), it is likely that this material will not fail due to mechanical strain within the operative range of typical biohybrid devices (Morimoto et al., 2018; Ricotti et al., 2017).

##### Asiga DentaGUM

Autoclave sterilization led to a decrease in the elastic domain properties of DentaGUM resins, with lower Young’s Modulus (5.1 ± 0.3%), Stress at 50% (8.9 ± 0.3%, *p<* 0.05 comparing NS to Autoclave), and Stress at 100% (11.2 ± 0.6%, *p<* 0.05 comparing NS to autoclave) (Fig. 11 (a), Column 3). Conversely, ethanol/UV sterilization led to no statistically significant difference in Stress at 50% or 100% compared to the nonsterile group, and means were within 5% (stress at 50%: 4.7 ± 0.3%, stress at 100%: −3.2 ± 0.2%). The Young’s modulus was slightly higher than that of the nonsterile group (9.2 ± 0.7%). Despite these changes in the elastic regime of these materials, there were no statistically significant differences in the failure properties of the different sterilization groups, with both the autoclave- and ethanol/UV-sterilized group means within 5% of the nonsterile mean for Ultimate Tensile Strength (Fig. 11(b)). When comparing the elongation at break, the mean for the autoclaved group was slightly greater than 5% of the nonsterile mean (6.5 ± 0.6%), but the variability in this metric for both the autoclaved and nonsterile groups prevents this from being a significant difference. Given these results, ethanol/UV sterilization may be the more appropriate sterilization technique for this resin to preserve pre-sterilization properties.

##### Constitutive Modeling

In addition to the scalar metrics discussed above, the collected stress and strain data were used to fit Yeoh hyperelastic constitutive models using a bootstrapping approach for each test group (Table 10, Figs. B.21-B.23). For all elastomeric resins, the model captured the mechanical response of the material well, with all groups except FormLabs Silicone IPA/BuOAc achieving an *R*^2^ greater than 0.98. Even FormLabs Silicone Mix reached an *R*^2^ of greater than 0.85, and the lower *R*^2^ values can be attributed to the variability in measured stress-strain curves for this material. For most materials, the upward-bending stress-strain curve requires the higher order terms in the Yeoh model. However, for 3Dresyns Bioflex IPA across all three sterilization conditions, the *C*_2_ and *C*_3_ model parameters were centered on 0, meaning they could likely be ignored without loss of accuracy. This would reduce the model simply to a Neohookean model with the same value of *C*_1_ (Yeoh and Fleming, 1997).

#### 3.2.2. Rigid Resin Mechanical Properties

Tensile and compression tests were also conducted on the rigid resin groups. The stress-strain curves for these materials are also reported in Appendix B. For these materials, the strain experienced in the elastic region was much smaller, as expected, so the stress at 50% and 100% were not appropriate measures. Therefore, for these materials, we calculated the Young’s modulus, Ultimate Tensile Strength (UTS), and elongation at break from the tensile tests, and the compressive modulus from the compression tests. These metrics were measurable for all samples and all test groups.

##### Asiga DentaGUIDE

Compared to the other rigid resins tested (see below), Asiga DentaGUIDE performed the best under the different sterilization techniques, only seeing a significant increase in the compressive modulus for both groups (autoclave: 15.1 ± 0.9%, ethanol/UV: 17.7 ± 0.8%). Both groups also saw similar increases in their elongation at break (autoclave: 24 ± 8, ethanol/UV: 19 ± 9%), though for neither group was this increase significant. The two sterilization groups differ in the change that was seen in the Young’s moduli and in the Ultimate Tensile Strength. The autoclaved samples saw a 10 ± 2% decrease in Young’s modulus and a 4.1 ± 0.6% increase in UTS, while the ethanol/UV group saw a 4.1 ± 0.7% increase in Young’s modulus and 16 ± 4% decrease in UTS. However, these differences were not significant compared to the nonsterile group. Overall, both groups performed well under sterilization, with the autoclave group becoming slightly softer but with better failure characteristics.

##### Liqcreate Bio-Med Clear

Sterilization had a large impact on the mechanical properties of Liqcreate Bio-Med Clear. Both sterilization techniques saw statistically significant increases in sample compressive moduli (autoclave: 14.0 ± 1.0%, ethanol/UV: 20 ± 2%) and decreases in Ultimate Tensile Strengths (autoclave: 18 ± 3%, ethanol/UV: 21 ± 3%). In addition, the autoclaved group saw a statistically significant decrease in Young’s modulus (29 ± 4%). The ethanol/UV group also saw a large decrease in Young’s modulus (24 ± 7%), but the variance in this group means this difference did not reach significance. Similarly, both groups saw nonsignificant increases in their elongation at break (autoclave: 100 ± 60%, ethanol/UV: 37 ± 13%). Across all metrics, the two sterilization groups performed similarly, with no significant differences found between the two groups.

##### Phrozen AquaGray 8K

Phrozen AquaGray 8K was also heavily affected by the sterilization process, with statistically significant differences from the nonsterile group for all metrics measured. In tension, the resin became significantly softer and more ductile. The autoclaved groups saw a decrease in the Young’s modulus of 71 ± 13% (*p<* 0.05, compared to nonsterile), and the ethanol/UV group saw a 65±19% decrease (*p<* 0.05 compared to nonsterile, *p >* 0.05 compared to autoclaved group). Simultaneously, both groups saw an increase in the elongation at break, 90 ± 30% for the autoclave group and 60 ± 20% for the ethanol/UV group (*p<* 0.05 for both compared to nonsterile, but *p>* 0.05 when compared to each other). This failure happened at a much greater strain but at a lower Ultimate Tensile Strength, with the autoclave group seeing a decrease of 50 ± 8% and the ethanol/UV group seeing a decrease of 55 ± 12% (*p<* 0.05 compared to nonsterile, *p>* 0.05 compared to each other). Contrary to the decrease in Young’s Modulus, both groups saw an increase in Compressive modulus, with a 25 ± 4% increase for the autoclave group and a 140 ± 20% increase for the ethanol/UV group (*p<* 0.05 for both groups compared to nonsterile group). Additionally, unlike the other metrics where no difference existed between the ethanol/UV and autoclave groups, here, the ethanol/UV had a statistically higher compressive modulus than the autoclaved group.

Across all three rigid resin groups, the compressive modulus increased with sterilization, while the tensile modulus stayed the same or decreased. It is uncommon for polymers to experience a loss in tensile modulus while also exhibiting a rise in compressive modulus. However, this phenomenon can occur under specific circumstances.

For example, this phenomenon could be due to microstructural changes in the polymer network (Mark, 2007; Donald and Kramer, 1982). Polymers are made up of long chains of molecules that can be arranged in different ways. In their natural state, these chains might be tangled and disorganized. When exposed to tensile forces, these entangled chains have a tendency to resist separation, resulting in a greater initial modulus. However, when the stress reaches a certain magnitude, the chains can initiate separation, resulting in a reduction in overall stiffness as the material elongates. In contrast, when subjected to compressive stress, the entangled chains can pack together in a more efficient manner, resulting in a more compact or denser network. The increased density of the packing can result in more resistance to deformation, hence causing a rise in the compressive modulus.

Alternatively, this behavior could be due to particular forms of damage to the samples caused by the sterilization. Specific forms of damage can selectively impact the tensile characteristics of a polymer (Hawkins, 1984). For example, when the polymer chains are exposed to ultraviolet radiation or harsh chemicals, they might degrade, which increases their vulnerability to breaking when subjected to tension. This would result in a reduction in the tensile modulus. Conversely, compressive loading may not be as susceptible to this kind of damage. Under some circumstances, the presence of microcracks or cavities/voids within the material can be compressed to the point of closure, resulting in a temporary increase in its compressive strength. This mechanism may be at play more in the ethanol/UV sterilization group, as these resins are partially methacrylate-based, and it has been shown that even mild exposure to ethanol can cause leaching of methacrylate (André et al., 2018). Additionally, part of the sterilization method for the ethanol/UV group included a one-hour exposure to UV light in the biosafety hood. This exposure could have either caused or exacerbated damage to the samples.

#### 3.2.3. Effect of PBS Soak on Mechanical Properties

For both Liqcreate Bio-Med Clear and Phrozen AquaGray 8K, the Young’s moduli that were measured from our nonsterile tensile tests were consistently lower than those that have been previously reported (Liqcreate Bio-Med Clear: measured = 710±50 MPa, reported 2000 MPa (no error provided) (Liqcreate, 2024), Phrozen AquaGray 8K: measured = 380±37 MPa, reported = 1195-1915 MPa (Phrozen, a)). Asiga did not report a modulus for their DentaGUIDE resin, so this same observation could not be made. We suspected that this decrease in nonsterilized Young’s modulus was due to the 16-hour soak in 37°C PBS that all samples were subjected to prior to mechanical tests. Subsequent experiments that measured the Young’s modulus of these materials using cantilevered beams agree with this suspicion (data not shown). In these subsequent tests, we mechanically tested both samples that had and had not been soaked in PBS, and the measured Young’s modulus for nonsoaked beams was in much better agreement with previous literature, while the soaked beams generally agreed with the above-reported Young’s moduli from our primary experiments. Thus the metrics measured from these experiments should be interpreted in the context of PBS-soaked parts. The mechanical properties and the effects of different sterilization techniques on these resins when not soaked are beyond the scope of this current work. However, as the goal of this work is to investigate these properties for use in biohybrid robotics contexts, where these materials will be in consistent contact with similar saline solutions to protect the biological components, the mechanics under the soaked conditions as presented here, rather than the unsoaked data from the manufacturers, should be used in the biohybrid design process.

#### 3.2.4. Mechanical Analysis Limitations

While the experiments conducted here present important information toward understanding the impact of sterilization on the mechanical properties of 3D-printable resins, there exist several limitations that could be addressed in future studies. First, the resins were only tested in single-stroke experiments. While this follows the current ASTM standard protocols, these experiments only provided limited information about the performance characteristics of these materials, particularly for the soft resins. For a more thorough characterization of these materials, cyclic testing should be performed for tensile and compression loading conditions. Many elastomeric materials exhibit the Mullin’s effect (Diani et al., 2009), and these single-stroke experiments only allow for the investigation of the virgin stress-strain curve. Using cyclic tests that span the range of strains the material is expected to experience *in situ*, a more performance-relevant analysis could be conducted.

These cyclic tests would also allow for the investigation of another important factor that was not investigated here – viscoelasticity. Many of these elastomeric resins also exhibit some degree of viscoelasticity, and this could be characterized using cyclic tests at varying strain rates. This strain rate dependence was not confined only to the soft resins but was also demonstrated here by the rigid resins. The tensile samples were tested at different strain rates than the compression samples in accordance with the ASTM standards (∼70%/min for tensile tests and ∼25%/min for compression tests. Exact strain rates are dependent on the rest geometry, as a fixed displacement rate was used for the two protocols). When we calculated moduli for these different loading conditions we saw much lower compressive moduli than tensile moduli for all resins and sterilization methods. This would be expected for viscoelastic materials tested at different strain rates as the Young’s modulus tends to increase with strain rate. In future experiments, these values should either be measured at various strain rates to capture their strain rate dependence or a single strain rate should be chosen that aligns with the expected operational strain rates for the material.

There were also limitations in the available equipment and in the experimental design. Given the load capacity of the load cells available and the size constraints of the EnviroBath that was used to ensure samples were tested in physiologically relevant solutions, not all elastomeric samples were brought to their failure point. However, given the range of strains that are expected in many biohybrid systems, these materials are not likely to fail from overstraining, so this may not be relevant to these systems. Additionally, given a lack of existing data for these materials, an *a priori* power analysis could not be conducted, and thus the sample size was chosen in agreement with the ASTM standards. However, with this sample size, and the observed variances, we were unable to achieve sufficient power for equivalence tests, which would be beneficial for showing statistically that two groups are the same within some allowable tolerance. We conducted a non-statistical analysis using a 5% tolerance window, but future studies should increase the sample size to test these equivalences statistically.

Finally, for 3D-printed photoresins, there are a variety of manufacturing parameters that could lead to mechanical property differences. The layer height, printing orientation, and exposure time all affect the quality and properties of the print. As the goal of this study was primarily to investigate the effect of sterilization techniques on mechanical properties, these factors were excluded. However, for a more complete analysis of the mechanical properties of these resins, these variables should be investigated in future studies.

### 3.3. Print Fidelity

Manual scoring of print feature quality and degradation was assessed on print fidelity samples that were nonsterile (Figure 14), autoclaved (Figure 15), or ethanol/UV-sterilized (Figure 16) for each resin type. Individual scorer identity did not significantly affect the response (Appendix C.13). Among the elastomeric resins, both sterile and nonsterile Asiga DentaGUM samples scored closest to the ground truth. For the rigid resins, Asiga DentaGUIDE and Liqcreate Bio-Med Clear samples exhibited minimum feature size limits and quality aligned with manufacturer specifications. Sterile samples of Asiga DentaGUIDE and Liqcreate Bio-Med Clear were not significantly different from their nonsterile counterparts.

**Figure 14:**
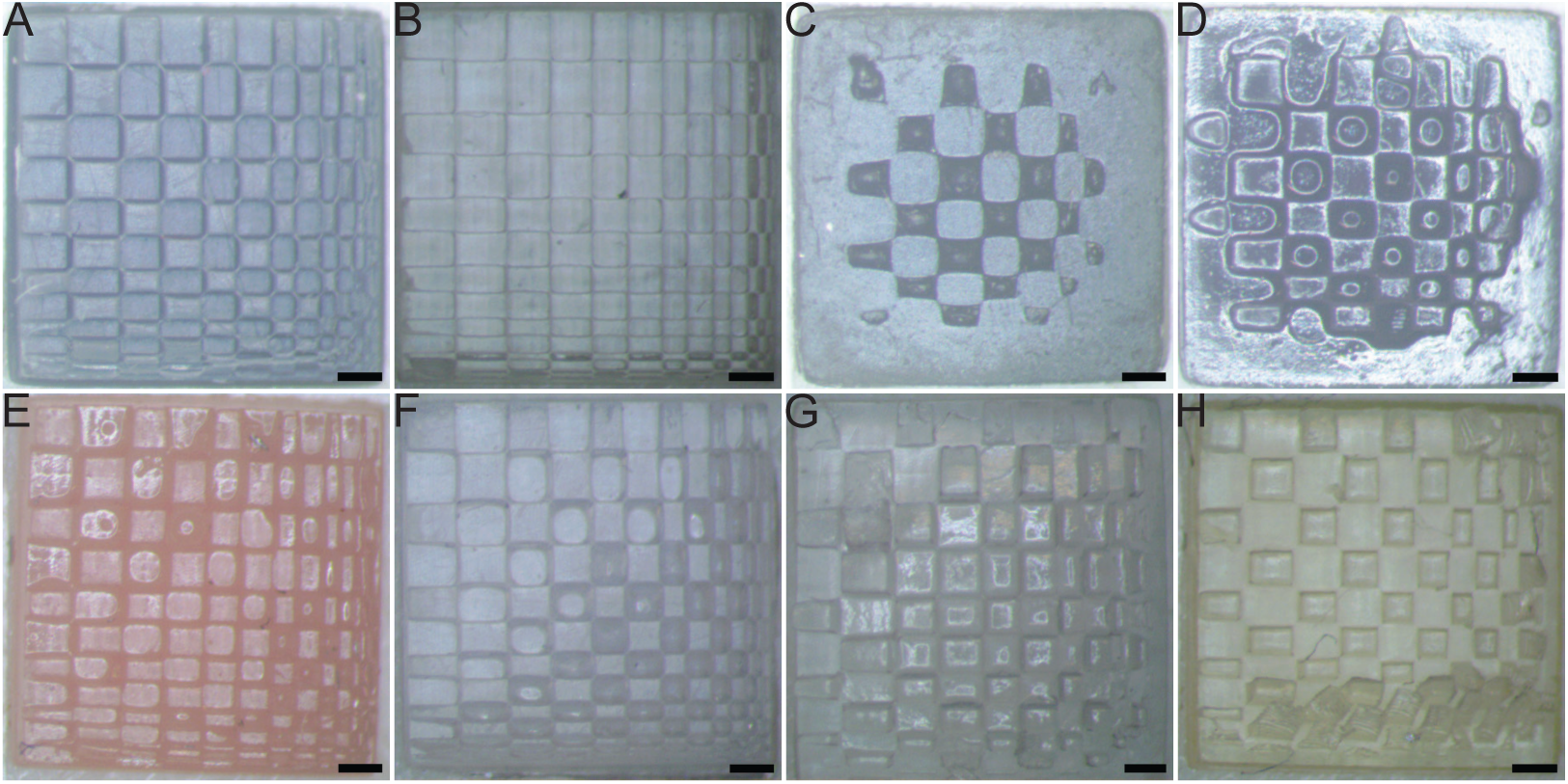
Representative samples for assessing the print fidelity of each nonsterile resin: (A) Phrozen AquaGray 8K; (B) Liqcreate Bio-Med Clear; (C) Formlabs Silicone 40A IPA/BuOAc post-treatment; (D) Formlabs Silicone 40A IPA-post-treatment; (E) Asiga DentaGUM; (F) Asiga DentaGUIDE; (G) 3Dresyns Bioflex A10 MB UNW2-posttreatment; (H) 3Dresyns Bioflex A10 MB IPA-post-treatment. All scale bars are 500 μm

**Figure 15:**
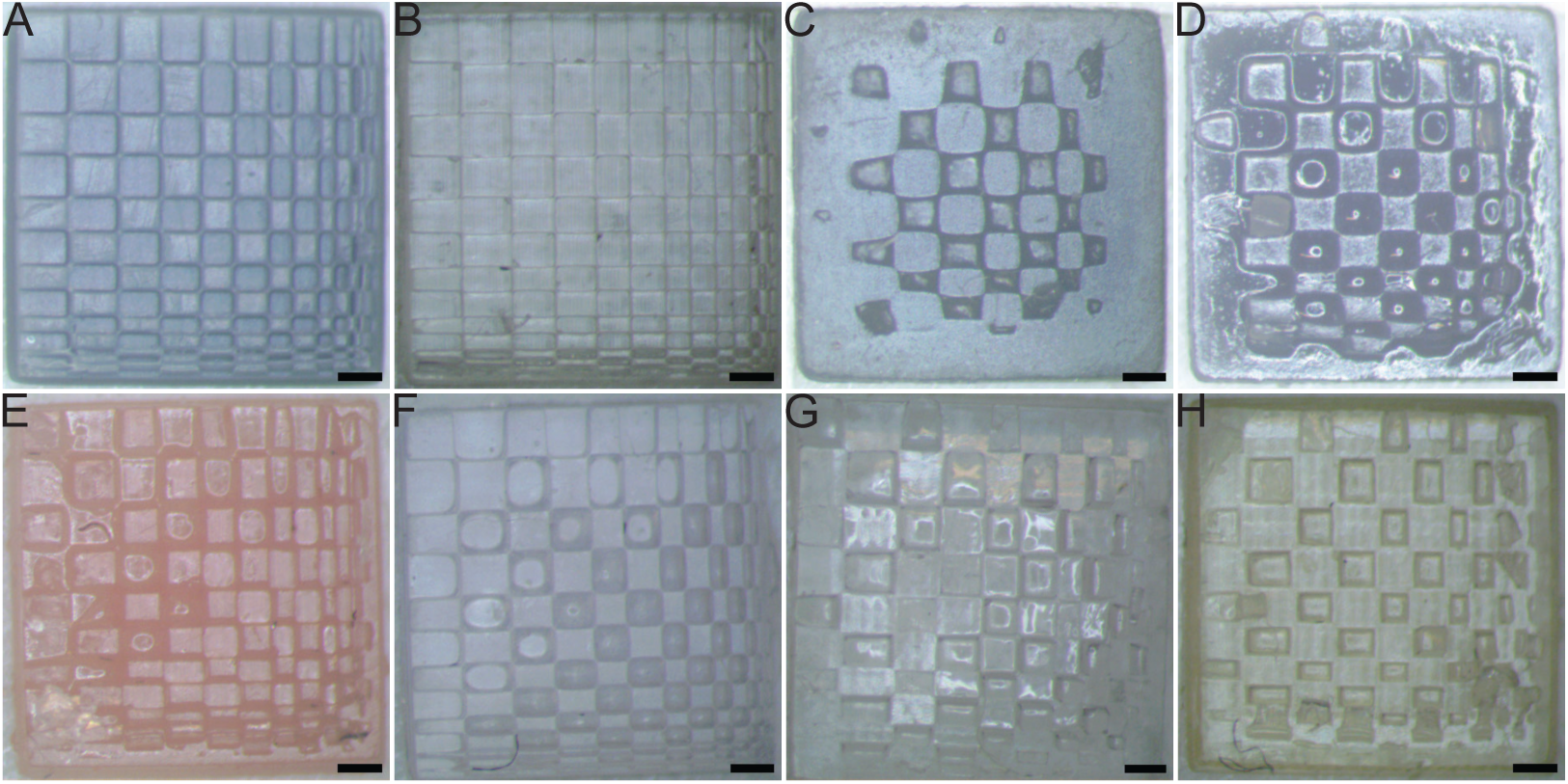
Representative samples for assessing the print fidelity of each autoclaved resin: (A) Phrozen AquaGray 8K; (B) Liqcreate Bio-Med Clear; (C) Formlabs Silicone 40A IPA/BuOAc post-treatment; (D) Formlabs Silicone 40A IPA-post-treatment; (E) Asiga DentaGUM; (F) Asiga DentaGUIDE; (G) 3Dresyns Bioflex A10 MB UNW2-posttreatment; (H) 3Dresyns Bioflex A10 MB IPA-post-treatment. All scale bars are 500 μm

**Figure 16:**
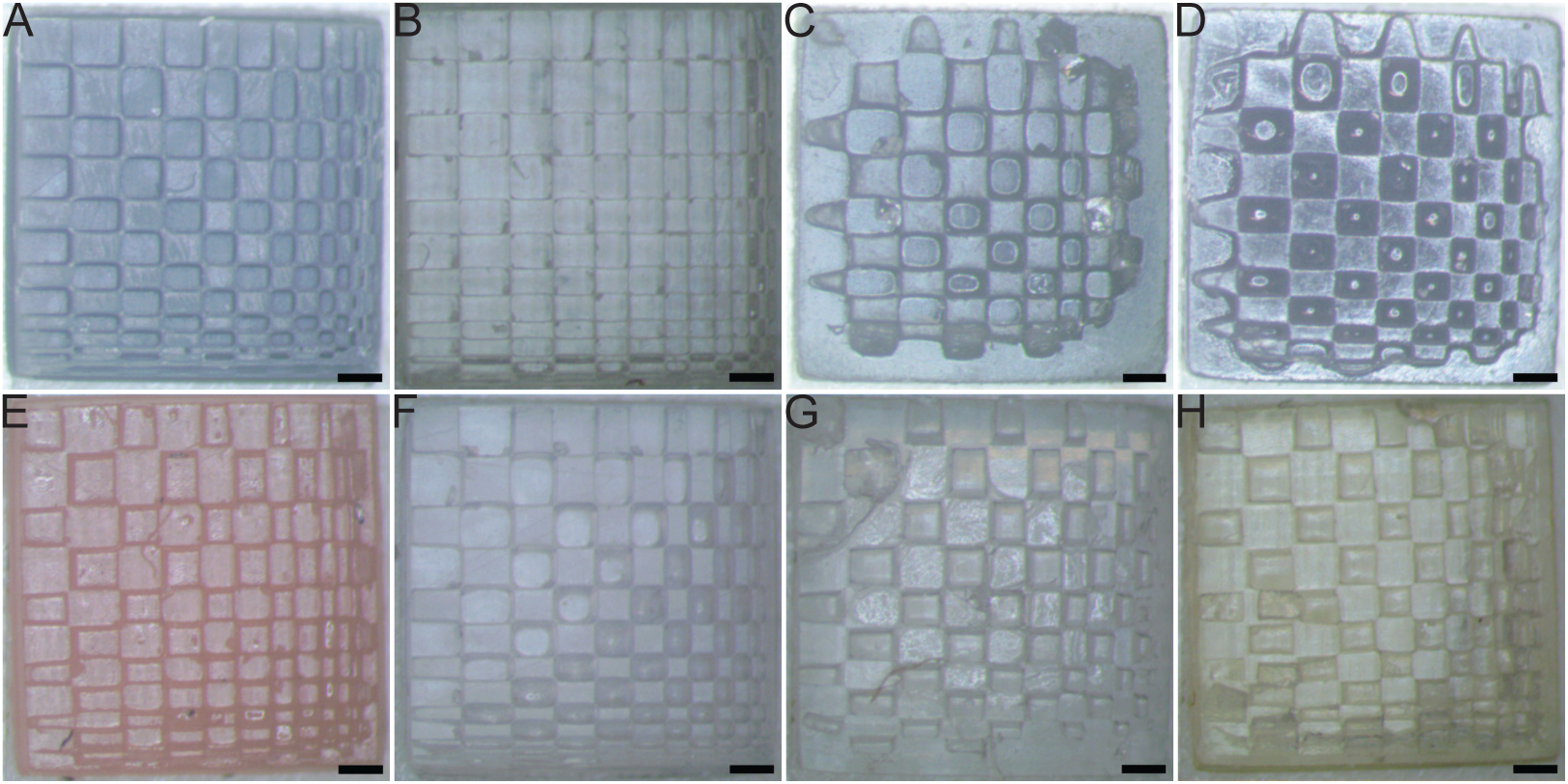
Representative samples for assessing the print fidelity of each ethanol/UV-sterilized resin: (A) Phrozen AquaGray 8K; (B) Liqcreate Bio-Med Clear; (C) Formlabs Silicone 40A IPA/BuOAc post-treatment; (D) Formlabs Silicone 40A IPA-post-treatment; (E) Asiga DentaGUM; (F) Asiga DentaGUIDE; (G) 3Dresyns Bioflex A10 MB UNW2-post-treatment; (H) 3Dresyns Bioflex A10 MB IPA-post-treatment. All scale bars are 500 μm

**Figure 17:**
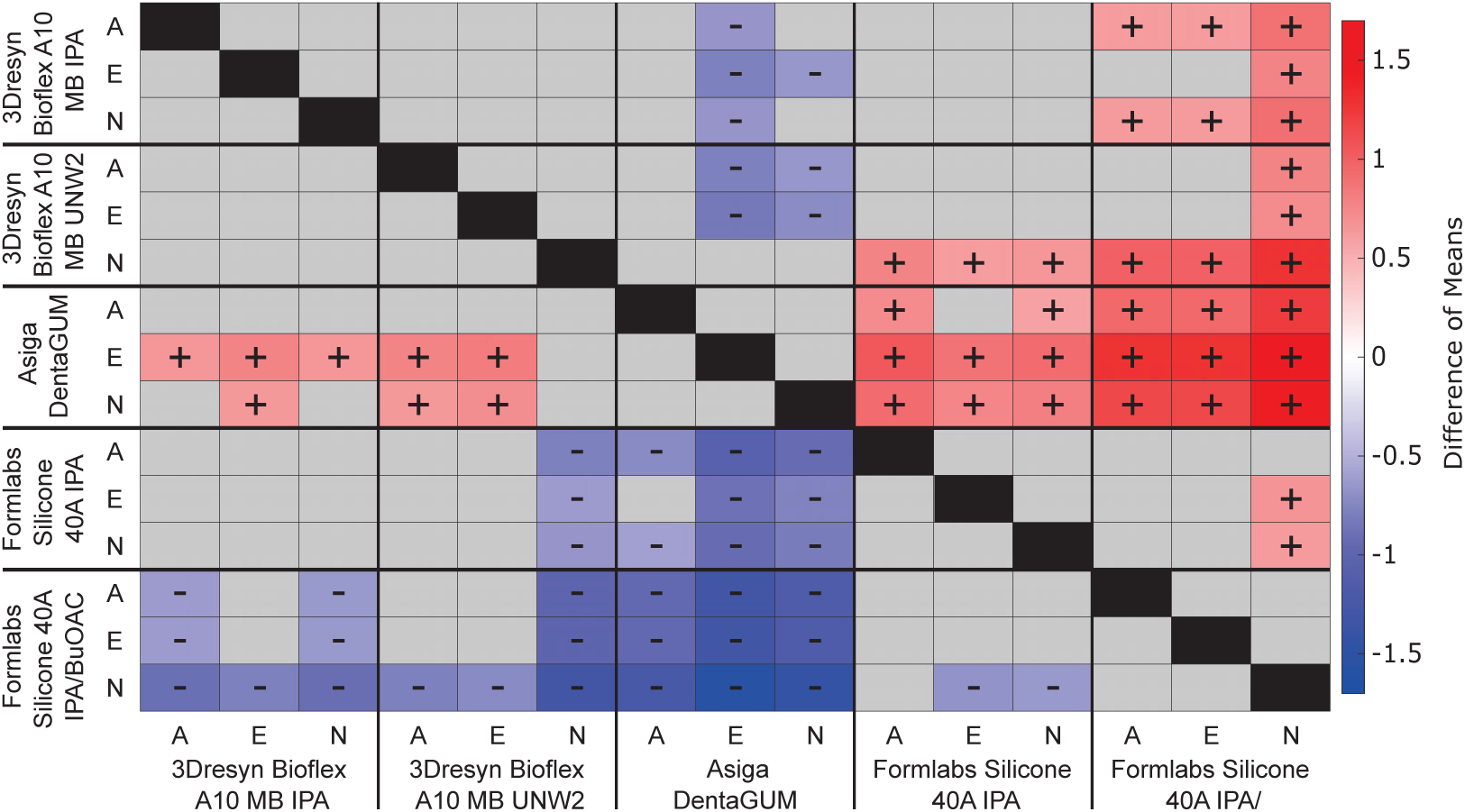
Comparison of the mean quality score for the entire sample for elastomeric resins. The colors and symbols represent the difference of means as calculated by subtracting the average quality of the left, vertical axis from the bottom, horizontal axis (positive value: red +, negative value: blue -). Grey boxes represent features with no significant difference. Sterilization factor: N - nonsterile, A - autoclave, E - ethanol/UV.

**Figure 18:**
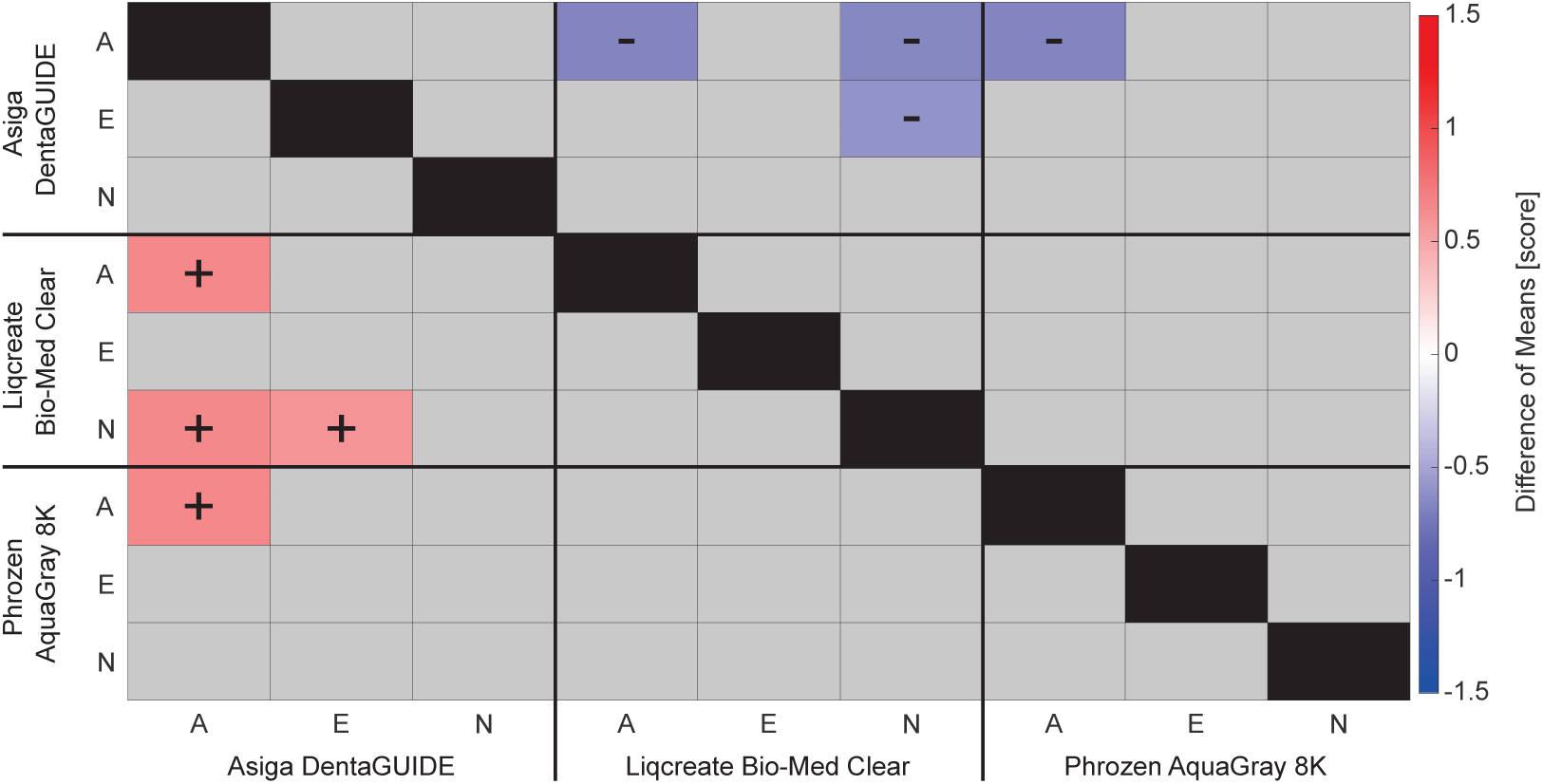
Comparison of the mean quality score for the entire sample for rigid resins. The colors and symbols represent the raw difference of means as calculated by subtracting the average quality of the left, vertical axis from the bottom, horizontal axis (positive value: red +, negative value: blue -). Grey boxes represent features with no significant difference. Sterilization factor: N - nonsterile, A - autoclave, E - ethanol/UV.

#### 3.3.1. Elastomeric Resin Print Fidelity

##### 3Dresyn Bioflex A10 MB

For all print fidelity features assessed, the 3Dresyn Bioflex A10 MB samples, both UNW2-treated and IPA-treated, showed no significant difference between the sterile and nonsterile samples (Figure C.36-C.43). However, the treatment type was found to cause significant differences. UNW2 post-treatment resulted in a significantly higher mean than IPA post-treatment in the response of the column with the smallest features and the quality of quadrant 2. (Table C.18, C.21) This was not surprising as the manufacturer notes that IPA and ethanol exposure may result in increased brittleness and haziness in the printed product (3dresyns).

##### Formlabs Silicone 40A

For samples post-treated with IPA/BuOAc, ethanol/UV-sterilization was found to significantly affect the row with the largest feature compared to nonsterile samples (Figure C.38). However, there was no significant difference between the two sterilization techniques or between autoclave-sterilized and nonsterile samples for any of the scored features. For the samples post-treated with IPA, there were no significant differences as a result of sterilization treatment.

To compare the two post-treatment types, the scores for nonsterile samples were compared. IPA-only post-treated samples were significantly different than those post-treated with IPA/BuOAc in the response of 6 out of 9 features and scored closer to the ground truth for all responses measured.(Figure C.36 - C.40 and Table C.15, C.16, C.17. C.18, C.19, C.20).

To compare with the manufacturer’s reported feature resolution, the row and column locations of the smallest resolvable features were analyzed. When manually assessing which column contained the smallest resolvable feature, samples post-treated with IPA or IPA/BuOAc achieved a mean score of 4.3 and 3.6, respectively. Similarly, for the row with the smallest resolvable feature, samples post-treated with IPA or IPA/BuOAc achieved a mean score of 4.2 and 3.4, respectively. The smallest designed dimensions for the features within column and row 3 were 0.4mm long, while the smallest feature size within column and row 4 was 0.3mm long (Figure 2). This is in line with the manufacturer’s reported achievable feature resolution of up to 0.3 mm when using their own printers (Formlabs 3 series) and post-printing systems (Form Wash and Form Cure) (Formlabs, 2024d).

##### Asiga DentaGUM

For all print fidelity features assessed, Asiga DentaGUM showed no significant difference between the sterile and nonsterile samples. Additionally, there was no significant difference between the two sterilization techniques. When considering only the resin type, Asiga DentaGUM was significantly closer to the ground truth scores than the other elastomeric resins (Tables C.15-C.23).

#### 3.3.2. Rigid Resin Print Fidelity

##### Asiga DentaGUIDE

For all print fidelity features assessed, Asiga DentaGUIDE showed no significant difference between the sterile and nonsterile samples, nor was there a significant difference between the two sterilization techniques.

No differences between the autoclave and nonsterile samples were expected since the manufacturer reports that DentaGUIDE is compatible with autoclave sterilization (Asiga, 2024a). However, the use of ethanol and UV sterilization was not reported. Since there were no significant differences between the ethanol/UV-sterilized and the autoclave-sterilized samples and no significant differences between the ethanol/UV-sterilized and nonsterile samples, the combination of ethanol and UV could also be a viable method for sterilizing Asiga DentaGuide without loss of print fidelity.

##### Liqcreate Bio-Med Clear

Liqcreate Bio-Med Clear, similarly showed no significant difference between the sterile and nonsterile samples or between the two sterilization techniques. The findings for autoclave sterilization are aligned with the manufacturer information (Liqcreate, 2024), and our findings further indicate that the combination of ethanol and UV may also be a viable method for sterilizing Liqcreate Bio-Med Clear.

When considering only the resin type, Liqcreate Bio-Med Clear received significantly closer to the ground truth than the other rigid resins for 5 of the 9 features (Table C.16, C.18, C.21, C.22, C.23). These features were: the column and row with the smallest feature, and quality in quadrants 2, 3, and 4.

##### Phrozen AquaGray 8K

There was no significant difference between ethanol/UV-sterilized and nonsterilized samples, but the ethanol/UV-sterilized samples were significantly different from the autoclaved samples. The ethanol/UV sterilized samples were further away from the ground truth than the autoclaved samples in 5 of 9 features: the column and row with the smallest feature and the quality of quadrants 2, 3, and 4 (Figure C.45, C.47, C.49, C.50, C.51). The formulation of Phrozen AquaGray 8K contains methacrylate which has been shown to chemically degrade when submerged in 70% ethanol (Phrozen, 2023; André et al., 2018). This could be a factor as to why autoclave-sterilized samples were closer to the ground truth.

When comparing the autoclaved samples with the nonsterile samples, only the quality of quadrant 4 was significantly different. For this feature, autoclave-sterilized samples were closer to ground truth with a mean value of 2.94 versus the nonsterile samples’ mean value of 2.26. When assessing the mean values, it is important to note that all manual scores are discrete integers as this is an inherent limitation from manually determining quality using Likert-type assessments. Since both means were between 2 and 3, this indicated a similar level of quality (Figure C.51).

#### 3.3.3. Print Fidelity Limitations

Print fidelity and the achievable minimum print size are dependent on the resin’s and the 3D printer’s properties. Different 3D printers have varying minimum resolutions based on differing technology, such as SLA, DLP, or LCD (refer to Section 1), and the possible power output can also impact the resin curing procedure during printing and result in variable final print quality. The Phrozen Sonic Mini 8K is capable of 1.1 mW/cm^2^ (measured on the UVA channel of an ams AS7331 UV sensor) while similar printers from Formlabs are capable of 16 mW/cm^2^ (Formlabs). This difference in power output could affect resin curing, leading to uncured resin becoming trapped inside each sample and poorer quality prints. To combat this, a longer cure time between printing layers can be used, but this would result in more time required to complete each layer, leading to longer print times.

For some samples print quality is lower along the edges with the largest features as seen from the responses to the row and column features (C.14). This could be a result of incidental damage during standard post-processing or diffusion of reactive species during printing. For all 3D printed products, parts must be removed from the build plate by scraping. However, these parts are not fully cured at the time of removal, potentially causing them to be softer and more prone to damage, particularly for features close to the edge of a sample. Furthermore, resins with low crosslinking density or step-growth polymerization may experience significant diffusion or convection of reactive species at the edges of the parts. Incomplete polymer chains may be washed away from the edges by viscous interactions with the resin during the raising and lowering of the build plate between printing layers. Therefore, when designing a part, the location of critical features must be considered to provide buffer space and preserve critical features.

Not only did the printing and post-processing impact the final print fidelity of a sample, but the sterilization procedure also impacted some of the resins in this study. Since sterilized samples must undergo extreme environmental conditions (high heat and pressure in the autoclaving process or long exposure to harsh chemicals in alcohol-based methods), the impacts of sterilization on these samples were not surprising. In particular, although alcohols, such as ethanol, are commonly used for sterilization prior to use with biological tissues, resin manufacturers also commonly recommend alcohols for washing and post-processing 3D-printable resins (Liqcreate, 2024; Phrozen, b). The alcohols are used to remove excess, uncured resins that may cling to the surface or stay in the cavities of a 3D-printed product. However, when submerged in 70% ethanol, cured resins containing methacrylate have been shown to chemically degrade (André et al., 2018). Methacrylate was found in the formulations of several resins included in this study (Phrozen AquaGray 8K (Phrozen, 2023), Liqcreate Bio-Med Clear (Liqcreate, 2023), Asiga DentaGUM (ASIGA, 2023b), and Asiga DentaGUIDE (ASIGA, 2023a)). Therefore, the ethanol exposure in combination with the presence of methacrylate could potentially cause leaching and explain the degradation of the samples qualitatively observed (Figure 16). To investigate this hypothesis in the future, the amount of methacrylate remaining in ethanol after sterilization could be measured. Additionally, if ethanol or autoclave sterilization is not compatible with a given resin, alternative sterilization techniques, such as gamma irradiation or plasma treatment, could be explored and assessed for their impacts on sample feature fidelity (Dai et al., 2016).

## 4. Conclusions

To identify suitable resins for use with low-cost, accessible manufacturing systems for biohybrid actuators composed of C2C12, this study investigated the properties of six commercially available resins with varying degrees of biocompatibility and material properties when printed using a Phrozen Sonic Mini 8K and post-cured using the Phrozen Curing Station. These results should be interpreted in the context of this low-cost manufacturing approach and not broadly to all printers as printing and curing power is likely to affect these findings. In this analysis, cytotoxicity was used as a proxy for biocompatibility. For the rigid resins, Asiga DentaGUIDE had the highest CalAM:EthD-1 ratio. Of the elastomeric resins, Formlabs Silicone 40A with either IPA or IPA/BuOAc post-treatments was the only elastomeric resin observed to be compatible with C2C12 cells for use in biohybrid actuators when manufactured as described here. Future work is needed to assess these resins more broadly for use with alternative printers and curing stations, particularly to assess the amount of uncured monomers and leachants after extended periods submerged in culture media.

From the mechanical analysis, the rigid resin groups were, in general, more greatly affected by the different sterilization techniques, with many of the soft resins showing no significant difference in their mechanical properties. Both Phrozen AquaGray 8K and Liqcreate Bio-Med Clear became much less stiff in tension after sterilization, and Phrozen AquaGray 8K became much more ductile. Asiga DentaGUIDE was more stable in its properties, with only the compressive modulus being significantly different. While the mean properties before and after sterilization were not very different for the two Formlabs Silicone post-treatment groups, the IPA/BuOAc post-treated samples saw a much larger property variability compared to the IPA-only group, suggesting that only IPA should be used for post-treating these parts.

Overall, print fidelity is highly dependent on only the resin type. The fidelity of rigid resin samples was not significantly impacted by the sterilization method, as compared to their nonsterile counterparts and the ground truth. However, sterilization impacted the fidelity of the elastomeric samples. Asiga DentaGUM samples were least affected, as compared to their nonsterile counterparts, but still deviated from the ground truth.

Based on the findings reported here, Asiga DentaGUIDE or Formlabs Silicone 40A are potential candidates for printing biohybrid base components using low-cost 3D-printing hardware. Furthermore, their mechanical properties were minimally impacted by sterilization. However, the print fidelity of Formlabs Silicone 40A samples was significantly affected by sterilization, so the minimum feature size of a biohybrid actuator design must be carefully considered if this resin will be used. Together, these findings provide a basis for researchers to select candidate materials for use in biohybrid robotics with low-cost lab-based manufacturing approaches, with a goal of broadening access to biohybrid robotics research.

## Supporting information

Graphical Abstract

## Acknowledgements

The authors would like to acknowledge that elements of the graphical abstract was created with Biorender.com.

## Author Statement

**Ashlee S. Liao:** Conceptualization, Methodology, Software, Validation, Formal Analysis, Investigation, Resources, Data Curation, Writing - Original Draft, Writing - Review & Editing, Visualization, Supervision, Project Administration, Funding Acquisition **Kevin Dai:** Conceptualization, Methodology, Validation, Investigation, Resources, Writing - Review & Editing, Funding Acquisition **Alaeddin Burak Irez:** Conceptualization, Methodology, Validation, Investigation, Data Curation, Writing - Original Draft, Writing - Review & Editing, Funding Acquisition **Anika Sun:** Methodology, Software, Validation, Formal Analysis, Investigation, Data Curation, Writing Original Draft, Writing - Review & Editing, Visualization **Michael Bennington:** Methodology, Software, Formal Analysis, Data Curation, Writing Original Draft, Writing - Review & Editing, Visualization **Saul Schaffer:** Methodology, Investigation, Writing - Review & Editing, Funding Acquisition **Bhavya Chopra:** Methodology, Investigation, Writing - Review & Editing **Ji Min Seok:** Methodology, Investigation, Writing - Review & Editing **Rebekah Adams:** Investigation, Writing - Review & Editing **Yongjie Jessica Zhang:** Writing - Review & Editing, Supervision, Funding Acquisition **Victoria A. Webster-Wood:** Conceptualization, Methodology, Validation, Formal Analysis, Resources, Data Curation, Writing - Review & Editing, Visualization, Supervision, Project Administration, Funding Acquisition

## Data Availability

All data, CAD files, and code associated with this work can be found in the associated online Zenodo repository:

https://doi.org/10.5281/zenodo.14014330

## 4.1. Generative AI statement

During the preparation of this work, the authors used the Grammarly Chrome Extension in order to improve grammar, spelling, and clarity. No Generative AI features were used to create new text. After using this tool, the authors reviewed and edited content further as needed and took full responsibility for the content of the publication.

