## Appendix for "Cytotoxicity and Characterization of 3D-Printable Resins Using a Low-Cost Printer for Muscle-based Biohybrid Devices"

### Appendix A. Mechanical Testing Sample Geometries

Table A.11: Average geometries of mechanical testing tensile samples. The gauge length is defined by the distance between the jaws that secured the sample in the test setup prior to loading. N is the sample size. For the original CAD geometries used to print the samples, refer to Figure 1 and the supplementary materials.

| Sterilization & Resin | | | N | Gauge Length | Thickness | Width |
|---|---|---|---|---|---|---|
| **Autoclave** | *Rigid* | DentaGUIDE | 7 | 6.23 ± 1.27 | 0.71 ± 0.03 | 1.06 ± 0.07 |
| | | AquaGray 8K | 6 | 7.44 ± 0.34 | 0.72 ± 0.07 | 1.12 ± 0.04 |
| | | Bio-Med Clear | 6 | 5.51 ± 0.64 | 0.77 ± 0.04 | 1.10 ± 0.07 |
| | *Elastomeric* | Bioflex A10 MB IPA | 6 | 29.14 ± 1.41 | 1.47 ± 0.04 | 2.94 ± 0.05 |
| | | Bioflex A10 MB UNW2 | 6 | 29.02 ± 1.06 | 1.51 ± 0.06 | 3.10 ± 0.09 |
| | | DentaGUM | 6 | 26.23 ± 1.00 | 1.51 ± 0.03 | 3.02 ± 0.04 |
| | | Silicone 40A IPA | 6 | 28.25 ± 1.27 | 1.51 ± 0.03 | 2.94 ± 0.04 |
| | | Silicone 40A IPA/BuOAc | 6 | 26.34 ± 1.06 | 1.21 ± 0.04 | 2.86 ± 0.04 |
| **Ethanol/UV** | *Rigid* | DentaGUIDE | 6 | 7.59 ± 0.54 | 0.72 ± 0.02 | 1.00 ± 0.05 |
| | | AquaGray 8K | 6 | 7.01 ± 0.41 | 0.74 ± 0.09 | 1.11 ± 0.02 |
| | | Bio-Med Clear | 6 | 7.32 ± 0.40 | 0.73 ± 0.06 | 1.14 ± 0.08 |
| | *Elastomeric* | Bioflex A10 MB IPA | 6 | 26.32 ± 0.70 | 1.48 ± 0.05 | 2.93 ± 0.03 |
| | | Bioflex A10 MB UNW2 | 6 | 28.33 ± 0.92 | 1.45 ± 0.06 | 2.98 ± 0.08 |
| | | DentaGUM | 6 | 25.57 ± 0.65 | 1.50 ± 0.04 | 3.00 ± 0.01 |
| | | Silicone 40A IPA | 6 | 28.30 ± 1.10 | 1.53 ± 0.04 | 2.89 ± 0.06 |
| | | Silicone 40A IPA/BuOAc | 6 | 25.91 ± 0.59 | 1.21 ± 0.04 | 2.87 ± 0.03 |
| **Nonsterile** | *Rigid* | DentaGUIDE | 4 | 7.02 ± 0.23 | 0.69 ± 0.03 | 0.98 ± 0.03 |
| | | AquaGray 8K | 5 | 7.10 ± 0.58 | 0.72 ± 0.05 | 1.09 ± 0.05 |
| | | Bio-Med Clear | 5 | 7.23 ± 0.37 | 0.73 ± 0.07 | 1.05 ± 0.04 |
| | *Elastomeric* | Bioflex A10 MB IPA | 6 | 26.65 ± 0.79 | 1.43 ± 0.03 | 2.95 ± 0.07 |
| | | Bioflex A10 MB UNW2 | 6 | 29.12 ± 1.64 | 1.50 ± 0.04 | 3.00 ± 0.06 |
| | | DentaGUM | 6 | 26.29 ± 0.75 | 1.50 ± 0.04 | 3.03 ± 0.02 |
| | | Silicone 40A IPA | 6 | 27.02 ± 1.02 | 1.57 ± 0.02 | 2.90 ± 0.04 |
| | | Silicone 40A IPA/BuOAc | 6 | 25.57 ± 0.66 | 1.21 ± 0.04 | 2.87 ± 0.06 |

Table A.12: Average geometries of mechanical testing compression samples. The gauge length is defined by the distance between the jaws that secured the sample in the test setup prior to loading. N is the sample size. For the original CAD geometries used to print the samples, refer to Figure 1 and the supplementary materials.

| Sterilization | & | Resin | N | Gauge Length | Diameter |
|---|---|---|---|---|---|
| **Autoclave** | *Rigid* | DentaGUIDE | 6 | 4.99 ± 0.06 | 2.47 ± 0.05 |
| | | AquaGray 8K | 6 | 5.17 ± 0.04 | 2.49 ± 0.03 |
| | | Bio-Med Clear | 6 | 5.03 ± 0.01 | 2.49 ± 0.02 |
| | *Elastomeric* | Bioflex A10 MB IPA | 6 | 5.00 ± 0.02 | 11.51 ± 0.05 |
| | | Bioflex A10 MB UNW2 | 6 | 5.06 ± 0.04 | 11.73 ± 0.06 |
| | | DentaGUM | 6 | 5.03 ± 0.03 | 11.42 ± 0.03 |
| | | Silicone 40A IPA | 6 | 5.08 ± 0.05 | 11.51 ± 0.06 |
| | | Silicone 40A IPA/BuOAc | 6 | 4.71 ± 0.04 | 11.30 ± 0.04 |
| **Ethanol/UV** | *Rigid* | DentaGUIDE | 6 | 5.02 ± 0.02 | 2.47 ± 0.02 |
| | | AquaGray 8K | 6 | 5.13 ± 0.05 | 2.49 ± 0.02 |
| | | Bio-Med Clear | 6 | 5.06 ± 0.02 | 2.48 ± 0.03 |
| | *Elastomeric* | Bioflex A10 MB IPA | 6 | 4.99 ± 0.07 | 11.49 ± 0.05 |
| | | Bioflex A10 MB UNW2 | 6 | 5.01 ± 0.01 | 11.57 ± 0.05 |
| | | DentaGUM | 6 | 5.00 ± 0.01 | 11.34 ± 0.06 |
| | | Silicone 40A IPA | 6 | 5.04 ± 0.04 | 11.39 ± 0.13 |
| | | Silicone 40A IPA/BuOAc | 6 | 4.70 ± 0.05 | 11.29 ± 0.02 |
| **Nonsterile** | *Rigid* | DentaGUIDE | 6 | 5.01 ± 0.03 | 2.47 ± 0.03 |
| | | AquaGray 8K | 7 | 5.15 ± 0.08 | 2.48 ± 0.02 |
| | | Bio-Med Clear | 6 | 5.07 ± 0.12 | 2.60 ± 0.12 |
| | *Elastomeric* | Bioflex A10 MB IPA | 6 | 4.96 ± 0.04 | 11.48 ± 0.08 |
| | | Bioflex A10 MB UNW2 | 6 | 4.99 ± 0.03 | 11.52 ± 0.08 |
| | | DentaGUM | 6 | 5.01 ± 0.02 | 11.36 ± 0.02 |
| | | Silicone 40A IPA | 6 | 5.01 ± 0.01 | 11.46 ± 0.04 |
| | | Silicone 40A IPA/BuOAc | 6 | 4.67 ± 0.04 | 11.33 ± 0.04 |

### Appendix B. Material Stress-Strain Curves and Elastomer Model Fits

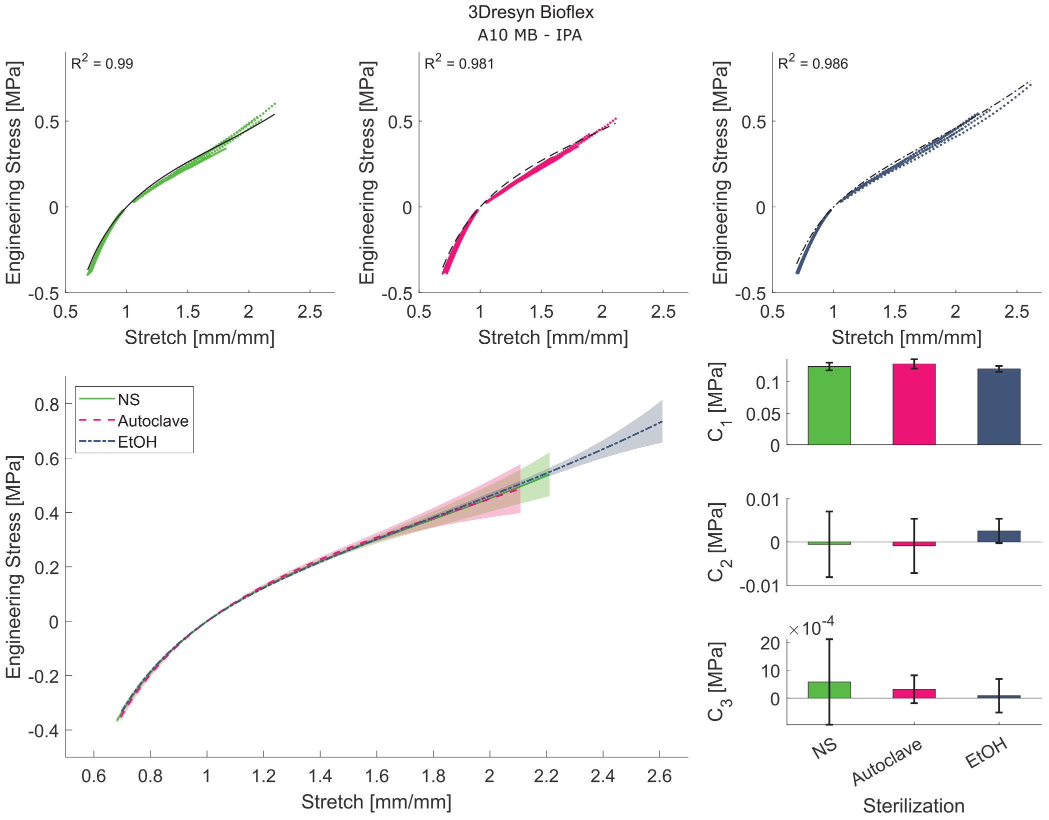


Figure B.19: Mechanical response of 3Dresyn Bioflex A10 MB post-treated with IPA for all sterilization groups. Top Row: Composite of all tensile and compressive stress-stretch for each sterilization technique (Left/green: nonsterile, Middle/pink: autoclave-sterilized, Right/blue: ethanol-sterilized) (stretch: $\lambda = 1 + \epsilon$). Black lines correspond to the mean Yeoh model fit determined from bootstrapped parameter estimation, and the $R^2$ value corresponds to the coefficient of determination of this mean model fit to all stress-stretch data. Bottom Left: Yeoh model response for the different sterilization groups. Lines indicate the mean model response, and the shaded region shows $\pm 1$ standard deviation. Bottom Right: Bar charts showing the bootstrapped model parameters $C_1$, $C_2$, and $C_3$. Bar height indicates mean parameter value, and error bars show $\pm 1$ standard deviation.

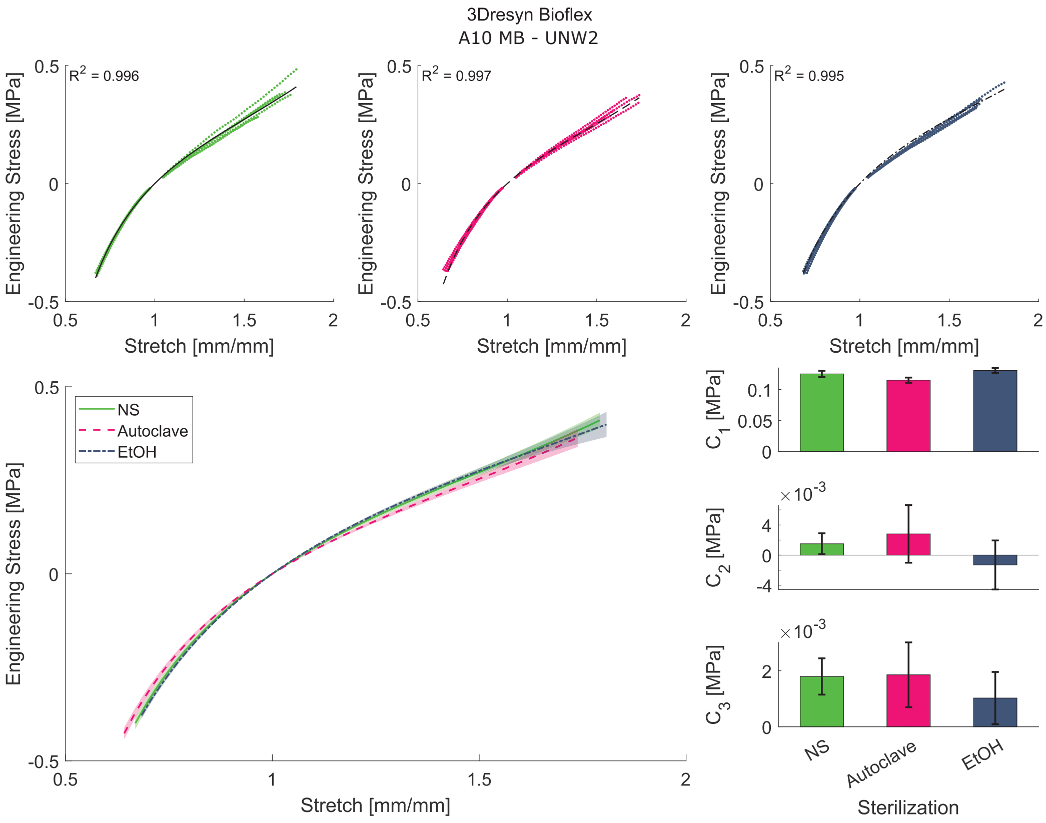


Figure B.20: Mechanical response of 3Dresyn Bioflex A10 MB post-treated with UNW2 for all sterilization groups. Layout and data follow the same layout as B.19. See caption for details.

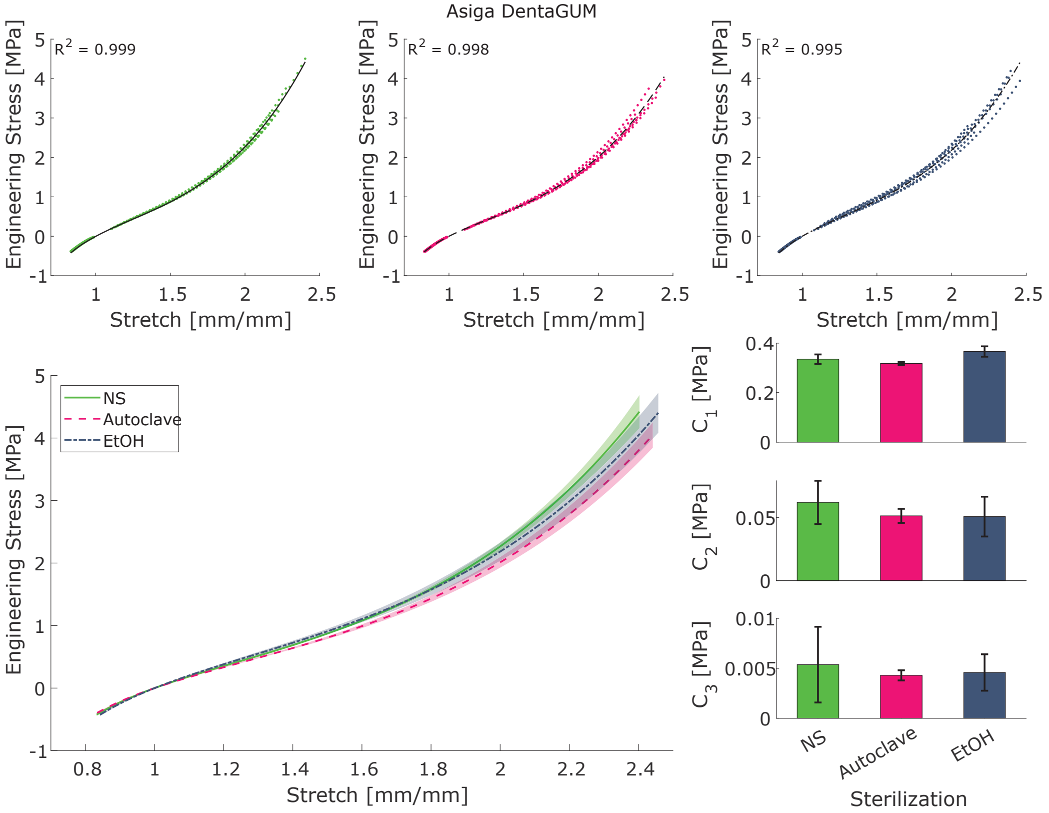


Figure B.21: Mechanical response of Asiga DentaGUM for all sterilization groups. Layout and data follow the same layout as B.19. See caption for details.

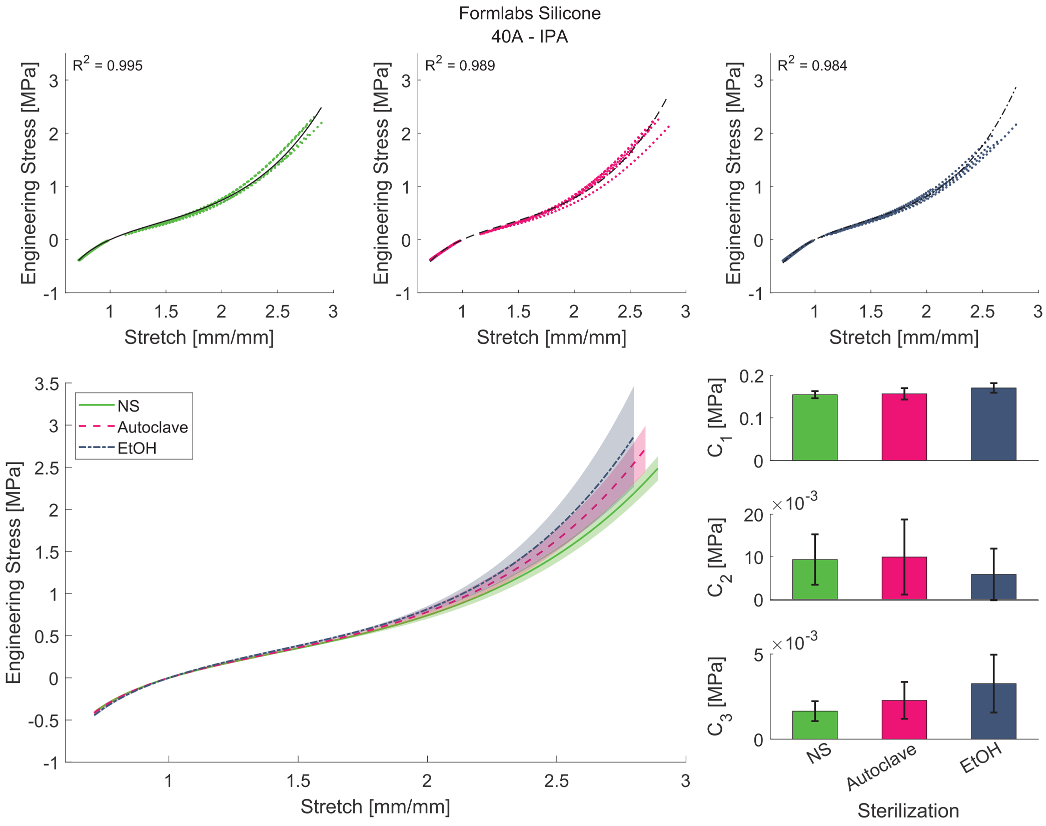


Figure B.22: Mechanical response of Formlabs Silicone post-treated with IPA for all sterilization groups. Layout and data follow the same layout as B.19. See caption for details.

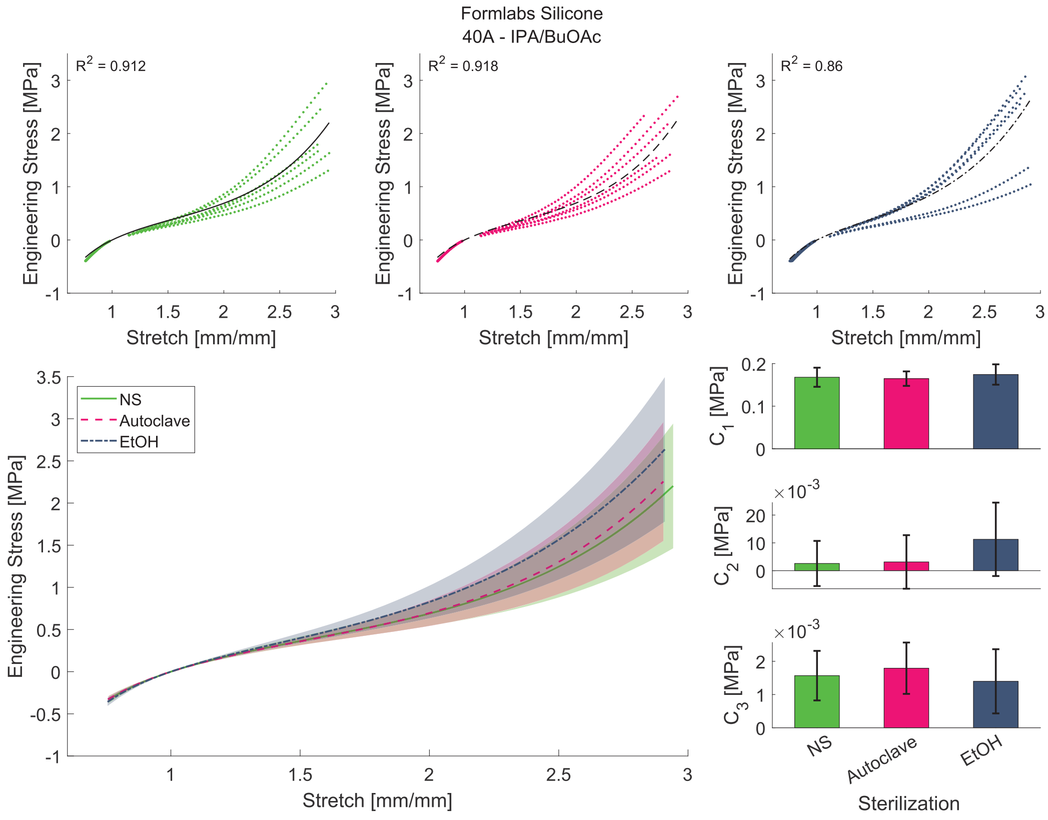


Figure B.23: Mechanical response of Formlabs Silicone post-treated with IPA+BuOAc for all sterilization groups. Layout and data follow the same layout as B.19. See caption for details.

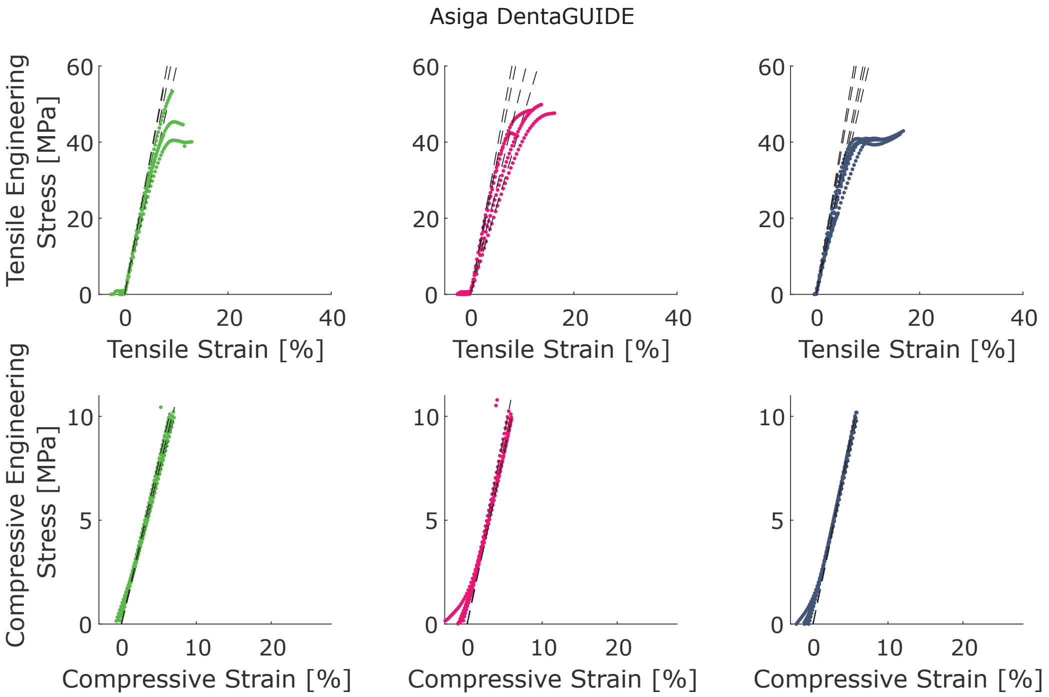


Figure B.24: Mechanical response of Asiga DentaGUIDE for all sterilization groups. Top Row: Tensile stress-strain curves. Dashed lines show the corresponding Hookean model approximation for each curve, the slope of which gives the Young's Modulus. Bottom Row: Compressive stress-strain curves. Here again, dashed lines show the Hookean model, this time with the slope giving the compressive modulus. Each column/color gives the data for a particular sterilization group (left/green: nonsterilize, middle/pink: autoclave-sterilized, right/blue: ethanol sterilized).

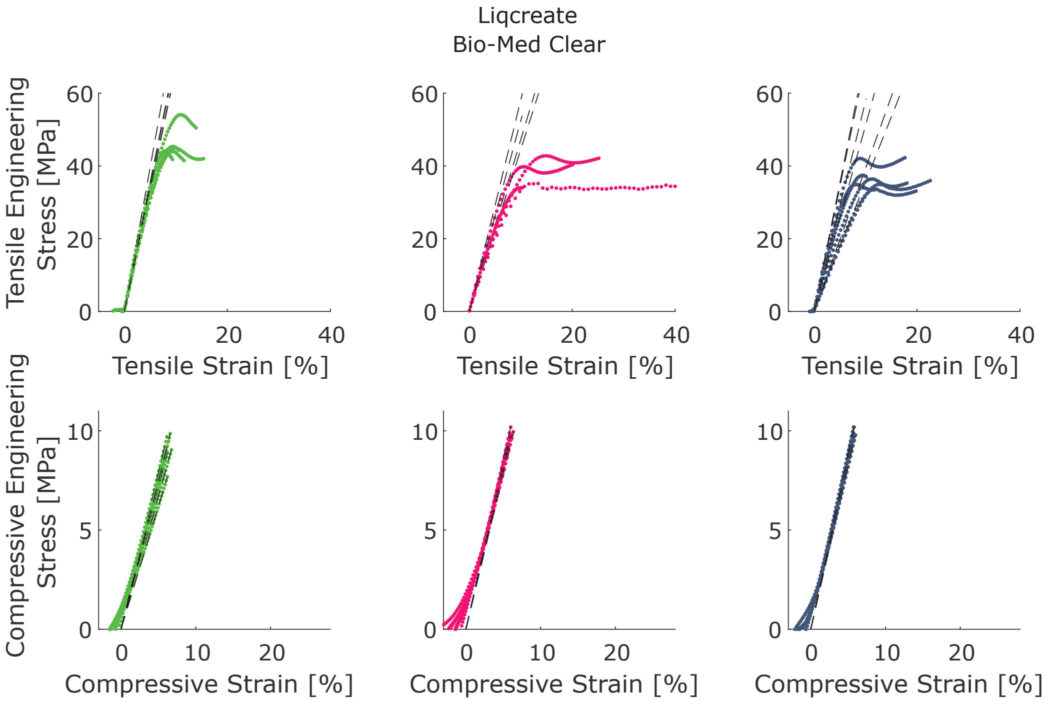


Figure B.25: Mechanical response of Liqcreate Bio-Med Clear for all sterilization groups. Layout and data follow the same layout as B.24. See caption for details.

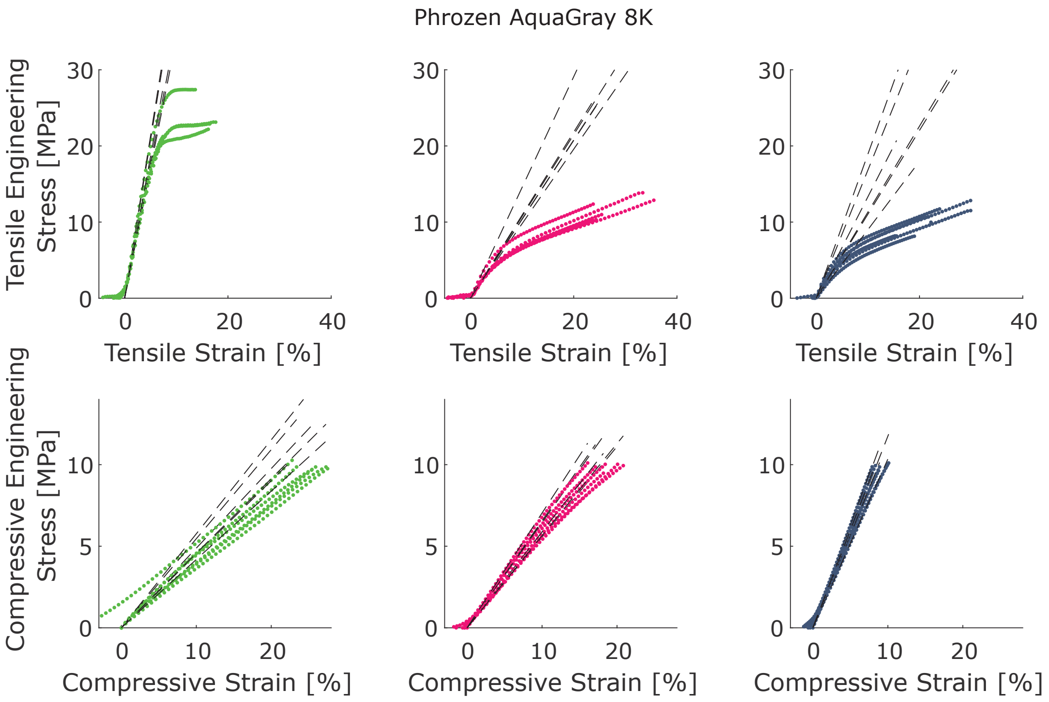


Figure B.26: Mechanical response of Phrozen AquaGray 8K for all sterilization groups. Layout and data follow the same layout as B.24. See caption for details.

### Appendix C. Mixed Effects Model fixed effects and variance components

Table C.13: Mixed effects models for assessing the variance contributed by the individual scorer on all manually assessed features for describing print fidelity. Var: variance component value; % of Total: Percentage of the total variation for each the variance component; SE Var: variance component standard error; Z: Z-value test statistic; $p$: probability value.

| Feature | Source | Var | % of Total | SE Var | Z | $p$ |
|---|---|---|---|---|---|---|
| Column Containing the Largest Feature | Scorer ID | 0.001 | 1.65% | 0.001 | 0.737 | 0.231 |
| | Error | 0.047 | 98.35% | 0.003 | 15.411 | 0.000 |
| | Total | 0.048 | | | | |
| Column Containing the Smallest Feature | Scorer ID | 0.237 | 42.70% | 0.239 | 0.992 | 0.161 |
| | Error | 0.318 | 57.30% | 0.021 | 15.411 | 0.000 |
| | Total | 0.556 | | | | |
| Row Containing the Largest Feature | Scorer ID | 0.003 | 5.81% | 0.003 | 0.911 | 0.181 |
| | Error | 0.048 | 94.19% | 0.003 | 15.411 | 0.000 |
| | Total | 0.051 | | | | |
| Row Containing the Smallest Feature | Scorer ID | 0.370 | 54.27% | 0.372 | 0.995 | 0.160 |
| | Error | 0.312 | 45.73% | 0.020 | 15.411 | 0.000 |
| | Total | 0.682 | | | | |
| Print Quality of the Entire Sample | Scorer ID | 0.289 | 52.99% | 0.290 | 0.995 | 0.160 |
| | Error | 0.256 | 47.01% | 0.017 | 15.411 | 0.000 |
| | Total | 0.545 | | | | |
| Print Quality of Quadrant 1 | Scorer ID | 0.397 | 59.66% | 0.398 | 0.996 | 0.160 |
| | Error | 0.268 | 40.34% | 0.017 | 15.411 | 0.000 |
| | Total | 0.665 | | | | |
| Print Quality of Quadrant 2 | Scorer ID | 0.054 | 13.93% | 0.056 | 0.964 | 0.167 |
| | Error | 0.334 | 86.07% | 0.022 | 15.411 | 0.000 |
| | Total | 0.388 | | | | |
| Print Quality of Quadrant 3 | Scorer ID | 0.024 | 7.30% | 0.026 | 0.929 | 0.176 |
| | Error | 0.309 | 92.70% | 0.020 | 15.411 | 0.000 |
| | Total | 0.333 | | | | |
| Print Quality of Quadrant 4 | Scorer ID | 0.140 | 29.44% | 0.142 | 0.986 | 0.162 |
| | Error | 0.336 | 70.56% | 0.022 | 15.411 | 0.000 |
| | Total | 0.476 | | | | |

Table C.14: Mixed effects models for assessing the fixed effects of the resin type, sterilization method, and interaction between the resin and sterilization on the manually assessed features for describing print fidelity. The column and row locations range from 1-7, where (row 1, column 1) was designed to contain the largest features and (row 7, column 7) was designed to contain the smallest features (Figure 2). For all analyses, the numerators for the degrees of freedom were 7, 2, and 14 for the resin, sterilization, and interaction between sterilization and resin sources, respectively. The denominator for the degrees of freedom was 475 for all sources. F: F-value test statistic; $p$: probability value.

| Feature | Source | F | $p$ |
|---|---|---|---|
| Column Containing the Largest Feature | Resin | 82.79 | 0.000 |
| | Sterilization | 1.33 | 0.266 |
| | Sterilization*Resin | 1.28 | 0.218 |
| Column Containing the Smallest Feature | Resin | 187.43 | 0.000 |
| | Sterilization | 2.72 | 0.067 |
| | Sterilization*Resin | 3.05 | 0.000 |
| Row Containing the Largest Feature | Resin | 84.79 | 0.000 |
| | Sterilization | 1.03 | 0.358 |
| | Sterilization*Resin | 2.44 | 0.002 |
| Row Containing the Smallest Feature | Resin | 197.43 | 0.000 |
| | Sterilization | 1.68 | 0.188 |
| | Sterilization*Resin | 2.32 | 0.004 |
| Print Quality of the Entire Sample | Resin | 175.95 | 0.000 |
| | Sterilization | 1.44 | 0.238 |
| | Sterilization*Resin | 2.29 | 0.005 |
| Print Quality of Quadrant 1 | Resin | 195.64 | 0.000 |
| | Sterilization | 6.10 | 0.002 |
| | Sterilization*Resin | 1.72 | 0.049 |
| Print Quality of Quadrant 2 | Resin | 148.78 | 0.000 |
| | Sterilization | 1.82 | 0.163 |
| | Sterilization*Resin | 2.32 | 0.004 |
| Print Quality of Quadrant 3 | Resin | 155.08 | 0.000 |
| | Sterilization | 2.51 | 0.083 |
| | Sterilization*Resin | 2.76 | 0.001 |
| Print Quality of Quadrant 4 | Resin | 189.63 | 0.000 |
| | Sterilization | 2.92 | 0.055 |
| | Sterilization*Resin | 2.87 | 0.000 |

Table C.15: Post-hoc Bonferroni pairwise comparisons for the manual assessment of the column containing the largest feature between each type of factor per source, as listed in the mixed effects model for all resins (C.14), with the exception of the interactions between sterilization and resin type (C.36,C.44). N is the number of samples per factor. Factors that share a Group letter per type of source are not significantly different.

| **Source** (bold)<br>Factor | N | Mean | Group | |
|---|---|---|---|---|
| **Sterilization** | | | | |
| Autoclave | 147 | 1.1 | A | |
| Nonsterile | 207 | 1.1 | A | |
| Ethanol/UV | 147 | 1.1 | A | |
| **Resin** | | | | |
| Formlabs Silicone 40A IPA/BuOAC | 61 | 1.7 | A | |
| 3dresyns Bioflex A10 MB IPA | 63 | 1.1 | | B |
| Formlabs Silicone 40A IPA | 62 | 1.1 | | B |
| 3dresyns Bioflex A10 MB UNW2 | 63 | 1.0 | | B |
| Asiga DentaGUM | 63 | 1.0 | | B |
| Asiga DentaGUIDE | 60 | 1.0 | | B |
| Liqcreate Bio-Med Clear | 63 | 1.0 | | B |
| Phrozen AquaGray 8K | 66 | 1.0 | | B |

Table C.16: Post-hoc Bonferroni pairwise comparisons for the manual assessment of the column containing the smallest feature between each type of factor per source, as listed in the mixed effects model for all resins (C.14), with the exception of the interactions between sterilization and resin type (C.37,C.45). N is the number of samples per factor. Factors that share a Group letter per type of source are not significantly different.

| **Source** (bold)<br>Factor | N | Mean | Group | | | | | |
|---|---|---|---|---|---|---|---|---|
| **Sterilization** | | | | | | | | |
| Autoclave | 147 | 5.0 | A | | | | | |
| NonSterile | 207 | 5.0 | A | | | | | |
| Ethanol/UV | 147 | 4.9 | A | | | | | |
| **Resin** | | | | | | | | |
| Liqcreate Bio-Med Clear | 63 | 6.5 | A | | | | | |
| Asiga DentaGUIDE | 60 | 5.9 | | B | | | | |
| Phrozen AquaGray 8K | 66 | 5.7 | | B | | | | |
| Asiga DentaGUM | 63 | 5.1 | | | C | | | |
| 3dresyns Bioflex A10 MB UNW2 | 63 | 4.6 | | | | D | | |
| Formlabs Silicone 40A IPA | 62 | 4.3 | | | | | E | |
| 3dresyns Bioflex A10 MB IPA | 63 | 4.1 | | | | | E | |
| Formlabs Silicone 40A IPA/BuOAC | 61 | 3.6 | | | | | | F |

Table C.17: Post-hoc Bonferroni pairwise comparisons for the manual assessment of the row containing the largest feature between each type of factor per source, as listed in the mixed effects model for all resins (C.14), with the exception of the interactions between sterilization and resin type (C.38,C.46). N is the number of samples per factor. Factors that share a Group letter per type of source are not significantly different.

| **Source** (bold)<br>Factor | N | Mean | Group | | |
|---|---|---|---|---|---|
| **Sterilization** | | | | | |
| Autoclave | 147 | 1.1 | A | | |
| Ethanol/UV | 147 | 1.1 | A | | |
| Nonsterile | 207 | 1.1 | A | | |
| **Resin** | | | | | |
| Formlabs Silicone 40A IPA/BuOAC | 61 | 1.8 | A | | |
| Formlabs Silicone 40A IPA | 62 | 1.1 | | B | |
| 3dresyns Bioflex A10 MB IPA | 63 | 1.1 | | B | C |
| 3dresyns Bioflex A10 MB UNW2 | 63 | 1.0 | | B | C |
| Asiga DentaGUIDE | 60 | 1.0 | | | C |
| Asiga DentaGUM | 63 | 1.0 | | | C |
| Liqcreate Bio-Med Clear | 63 | 1.0 | | | C |
| Phrozen AquaGray 8K | 66 | 1.0 | | | C |

Table C.18: Post-hoc Bonferroni pairwise comparisons for the manual assessment of the row containing the smallest feature between each type of factor per source, as listed in the mixed effects model for all resins (C.14), with the exception of the interactions between sterilization and resin type (C.39,C.47). N is the number of samples per factor. Factors that share a Group letter per type of source are not significantly different.

| **Source** (bold)<br>Factor | N | Mean | Group | | | | |
|---|---|---|---|---|---|---|---|
| **Sterilization** | | | | | | | |
| Autoclave | 147 | 5.0 | A | | | | |
| Nonsterile | 207 | 4.9 | A | | | | |
| Ethanol/UV | 147 | 4.9 | A | | | | |
| **Resin** | | | | | | | |
| Liqcreate Bio-Med Clear | 63 | 6.3 | A | | | | |
| Phrozen AquaGray 8K | 66 | 5.9 | | B | | | |
| Asiga DentaGUIDE | 60 | 5.7 | | B | | | |
| Asiga DentaGUM | 63 | 5.2 | | | C | | |
| 3dresyns Bioflex A10 MB UNW2 | 63 | 4.4 | | | | D | |
| Formlabs Silicone 40A IPA | 62 | 4.2 | | | | D | |
| 3dresyns Bioflex A10 MB IPA | 63 | 4.2 | | | | D | |
| Formlabs Silicone 40A IPA/BuOAC | 61 | 3.4 | | | | | E |

Table C.19: Post-hoc Bonferroni pairwise comparisons for the manual assessment of the quality of each sample between each type of factor per source, as listed in the mixed effects model for all resins (C.14), with the exception of the interactions between sterilization and resin type (17,18). N is the number of samples per factor. Factors that share a Group letter per type of source are not significantly different.

| **Source** (bold)<br>Factor | N | Mean | Group | | | | | |
|---|---|---|---|---|---|---|---|---|
| **Sterilization** | | | | | | | | |
| Nonsterile | 207 | 2.8 | A | | | | | |
| Autoclave | 147 | 2.7 | A | | | | | |
| Ethanol/UV | 147 | 2.7 | A | | | | | |
| **Resin** | | | | | | | | |
| Liqcreate Bio-Med Clear | 63 | 3.9 | A | | | | | |
| Phrozen AquaGray 8K | 66 | 3.8 | A | | | | | |
| Asiga DentaGUIDE | 60 | 3.4 | | B | | | | |
| Asiga DentaGUM | 63 | 2.8 | | | C | | | |
| 3dresyns Bioflex A10 MB UNW2 | 63 | 2.3 | | | | D | | |
| 3dresyns Bioflex A10 MB IPA | 63 | 2.2 | | | | D | E | |
| Formlabs Silicone 40A IPA | 62 | 2.0 | | | | | E | |
| Formlabs Silicone 40A IPA/BuOAC | 61 | 1.6 | | | | | | F |

Table C.20: Post-hoc Bonferroni pairwise comparisons for the manual assessment of the quality of quadrant 1 for each sample between each type of factor per source, as listed in the mixed effects model for all resins (C.14), with the exception of the interactions between sterilization and resin type (C.40,C.48). N is the number of samples per factor. Factors that share a Group letter per type of source are not significantly different.

| **Source** (bold)<br>Factor | N | Mean | Group | | | | | |
|---|---|---|---|---|---|---|---|---|
| **Sterilization** | | | | | | | | |
| Nonsterile | 207 | 3.3 | A | | | | | |
| Ethanol/UV | 147 | 3.1 | | B | | | | |
| Autoclave | 147 | 3.1 | | B | | | | |
| **Resin** | | | | | | | | |
| Phrozen AquaGray 8K | 66 | 4.5 | A | | | | | |
| Liqcreate Bio-Med Clear | 63 | 4.2 | A | | | | | |
| Asiga DentaGUIDE | 60 | 3.7 | | B | | | | |
| Asiga DentaGUM | 63 | 3.2 | | | C | | | |
| 3dresyns Bioflex A10 MB UNW2 | 63 | 2.9 | | | | D | | |
| 3dresyns Bioflex A10 MB IPA | 63 | 2.9 | | | | D | | |
| Formlabs Silicone 40A IPA | 62 | 2.3 | | | | | E | |
| Formlabs Silicone 40A IPA/BuOAC | 61 | 1.8 | | | | | | F |

Table C.21: Post-hoc Bonferroni pairwise comparisons for the manual assessment of the quality of quadrant 2 for each sample between each type of factor per source, as listed in the mixed effects model for all resins (C.14), with the exception of the interactions between sterilization and resin type (C.41,C.49). N is the number of samples per factor. Factors that share a Group letter per type of source are not significantly different.

| **Source** (bold)<br>Factor | N | Mean | Group | | | | | |
|---|---|---|---|---|---|---|---|---|
| **Sterilization** | | | | | | | | |
| Nonsterile | 207 | 2.3 | A | | | | | |
| Autoclave | 147 | 2.2 | A | | | | | |
| Ethanol/UV | 147 | 2.1 | A | | | | | |
| **Resin** | | | | | | | | |
| Liqcreate Bio-Med Clear | 63 | 3.6 | A | | | | | |
| Asiga DentaGUIDE | 60 | 3.0 | | B | | | | |
| Phrozen AquaGray 8K | 66 | 3.0 | | B | | | | |
| Asiga DentaGUM | 63 | 2.4 | | | C | | | |
| 3dresyns Bioflex A10 MB UNW2 | 63 | 1.8 | | | | D | | |
| 3dresyns Bioflex A10 MB IPA | 63 | 1.5 | | | | | E | |
| Formlabs Silicone 40A IPA | 62 | 1.3 | | | | | E | F |
| Formlabs Silicone 40A IPA/BuOAC | 61 | 1.1 | | | | | | F |

Table C.22: Post-hoc Bonferroni pairwise comparisons for the manual assessment of the quality of quadrant 3 for each sample between each type of factor per source, as listed in the mixed effects model for all resins (C.14), with the exception of the interactions between sterilization and resin type (C.42,C.50). N is the number of samples per factor. Factors that share a Group letter per type of source are not significantly different.

| **Source** (bold)<br>Factor | N | Mean | Group | | | | |
|---|---|---|---|---|---|---|---|
| **Sterilization** | | | | | | | |
| Autoclave | 147 | 2.2 | A | | | | |
| Nonsterile | 207 | 2.2 | A | | | | |
| Ethanol/UV | 147 | 2.1 | A | | | | |
| **Resin** | | | | | | | |
| Liqcreate Bio-Med Clear | 63 | 3.6 | A | | | | |
| Phrozen AquaGray 8K | 66 | 3.0 | | B | | | |
| Asiga DentaGUIDE | 60 | 2.9 | | B | | | |
| Asiga DentaGUM | 63 | 2.3 | | | C | | |
| 3dresyns Bioflex A10 MB UNW2 | 63 | 1.7 | | | | D | |
| 3dresyns Bioflex A10 MB IPA | 63 | 1.5 | | | | D | E |
| Formlabs Silicone 40A IPA | 62 | 1.4 | | | | | E |
| Formlabs Silicone 40A IPA/BuOAC | 61 | 1.2 | | | | | E |

Table C.23: Post-hoc Bonferroni pairwise comparisons for the manual assessment of the quality of quadrant 4 for each sample between each type of factor per source, as listed in the mixed effects model for all resins (C.14), with the exception of the interactions between sterilization and resin type (C.43,C.51). N is the number of samples per factor. Factors that share a Group letter per type of source are not significantly different.

| **Source** (bold)<br>Factor | N | Mean | Group |
|---|---|---|---|
| **Sterilization** | | | |
| Autoclave | 147 | 1.6 | A |
| Nonsterile | 207 | 1.6 | A |
| Ethanol/UV | 147 | 1.5 | A |
| **Resin** | | | |
| Liqcreate Bio-Med Clear | 63 | 3.2 | A |
| Asiga DentaGUIDE | 60 | 2.6 | B |
| Phrozen AquaGray 8K | 66 | 2.4 | B |
| Asiga DentaGUM | 63 | 1.8 | C |
| 3dresyns Bioflex A10 MB UNW2 | 63 | 0.8 | D |
| Formlabs Silicone 40A IPA | 62 | 0.8 | D E |
| 3dresyns Bioflex A10 MB IPA | 63 | 0.7 | D E |
| Formlabs Silicone 40A IPA/BuOAC | 61 | 0.5 | E |

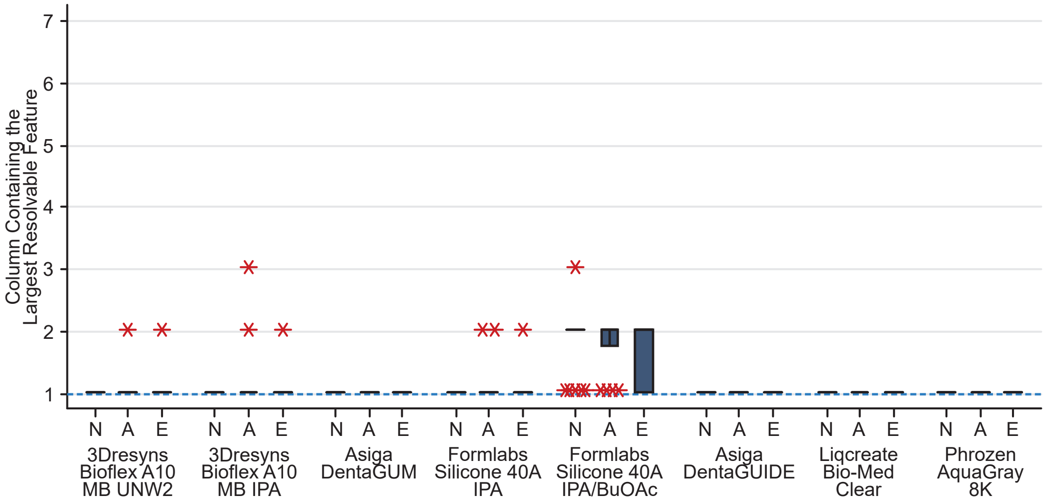


Figure C.27: The columns containing the largest resolvable features as determined by the manual assessment are represented as boxplots of the rigid (dark blue-gray) and elastomeric (light blue-gray) resin samples with one of the sterilization conditions: N - non-sterile, A - autoclave, E - ethanol/UV. The red asterisks (*) denote outliers. The upper and lower boundaries of the box represent the first and third quartiles. Whiskers extend from the interquartile boxes to the minimum and maximum values, excluding outliers. The blue dashed line represented the target value if the largest resolvable feature was located in the intended column based on the original CAD model (Figure 2).

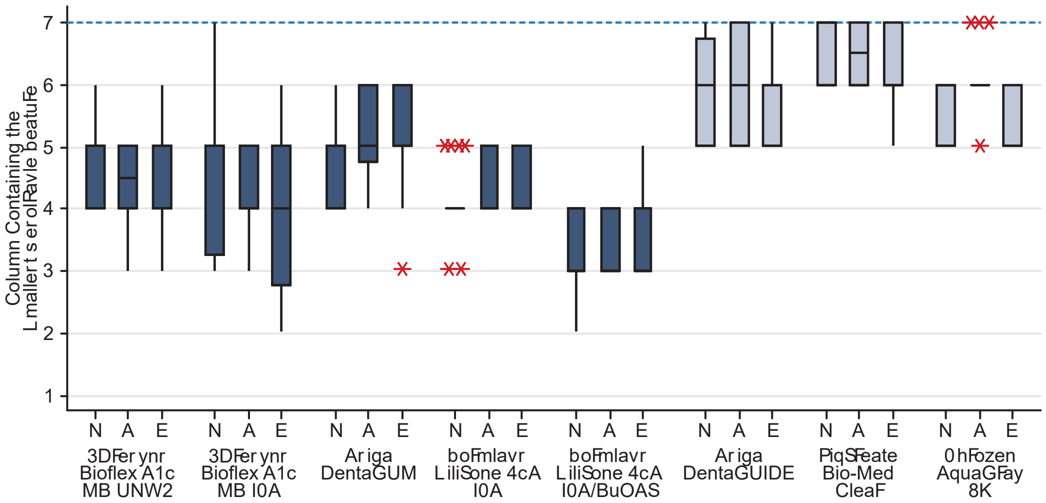


Figure C.28: The columns containing the smallest resolvable features as determined by the manual assessment are represented as boxplots of the rigid (dark blue-gray) and elastomeric (light blue-gray) resin samples with one of the sterilization conditions: N - non-sterile, A - autoclave, E - ethanol/UV. The red asterisks (*) denote outliers. The upper and lower boundaries of the box represent the first and third quartiles. Whiskers extend from the interquartile boxes to the minimum and maximum values, excluding outliers. The blue dashed line represented the target value if the smallest resolvable feature was located in the intended column based on the original CAD model (Figure 2).

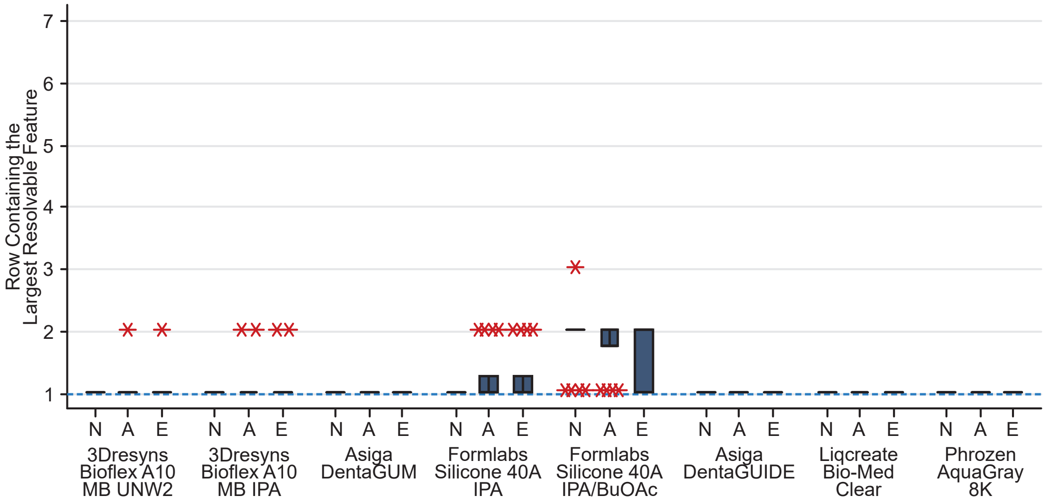


Figure C.29: The rows containing the largest resolvable features as determined by the manual assessment are represented as boxplots of the rigid (dark blue-gray) and elastomeric (light blue-gray) resin samples with one of the sterilization conditions: N - nonsterile, A - autoclave, E - ethanol/UV. The red asterisks (*) denote outliers. The upper and lower boundaries of the box represent the first and third quartiles. Whiskers extend from the interquartile boxes to the minimum and maximum values, excluding outliers. The blue dashed line represented the target value if the largest resolvable feature was located in the intended row based on the original CAD model (Figure 2).

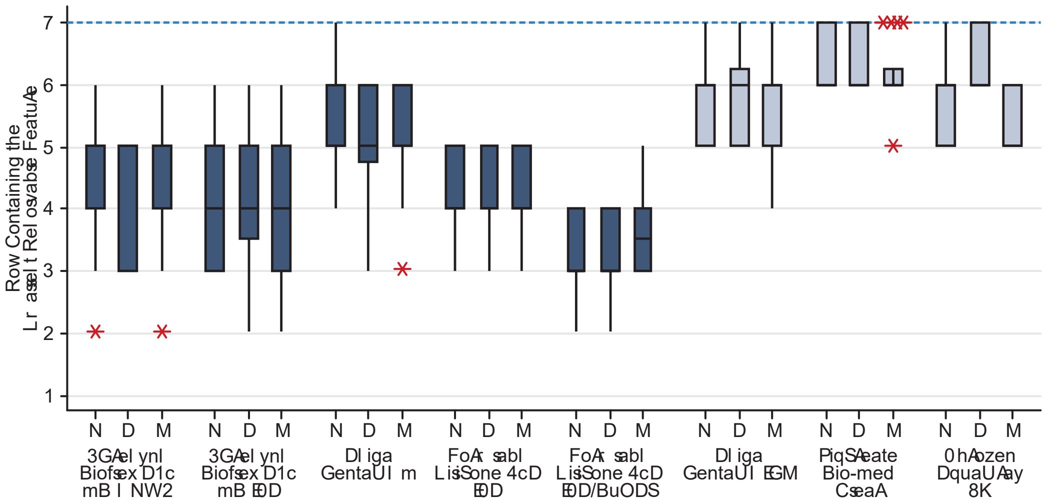

Figure C.30: The rows containing the smallest resolvable features as determined by the manual assessment are represented as boxplots of the rigid (dark blue-gray) and elastomeric (light blue-gray) resin samples with one of the sterilization conditions: N - non-sterile, A - autoclave, E - ethanol/UV. The red asterisks (*) denote outliers. The upper and lower boundaries of the box represent the first and third quartiles. Whiskers extend from the interquartile boxes to the minimum and maximum values, excluding outliers. The blue dashed line represented the target value if the smallest resolvable feature was located in the intended row based on the original CAD model (Figure 2).

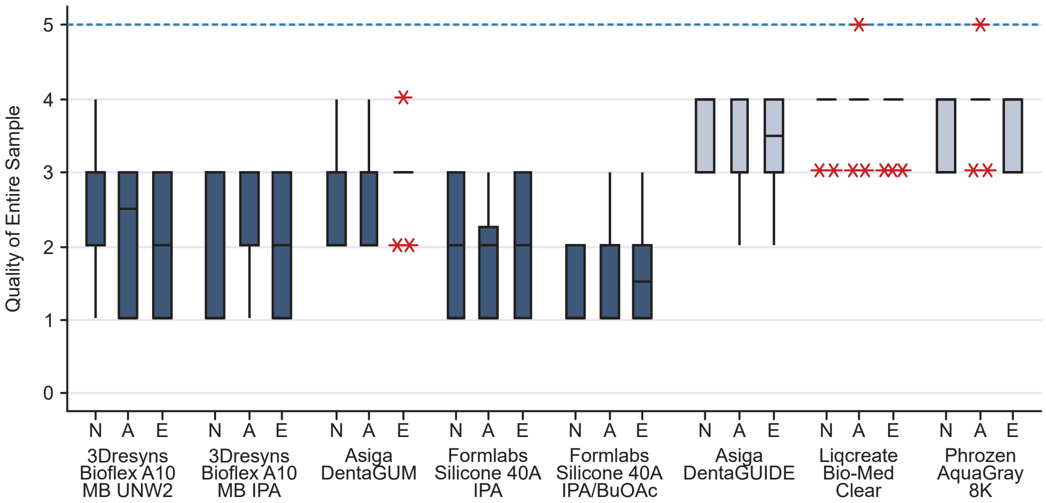


Figure C.31: The quality of the entire printed part as determined by the manual assessment are represented as boxplots of the rigid (dark blue-gray) and elastomeric (light blue-gray) resin samples with one of the sterilization conditions: N - nonsterile, A - autoclave, E - ethanol/UV. The red asterisks (*) denote outliers. The upper and lower boundaries of the box represent the first and third quartiles. Whiskers extend from the interquartile boxes to the minimum and maximum values, excluding outliers. The blue dashed line represented the score awarded if the sample qualitatively appeared equivalent to the original CAD model (Figure 2).

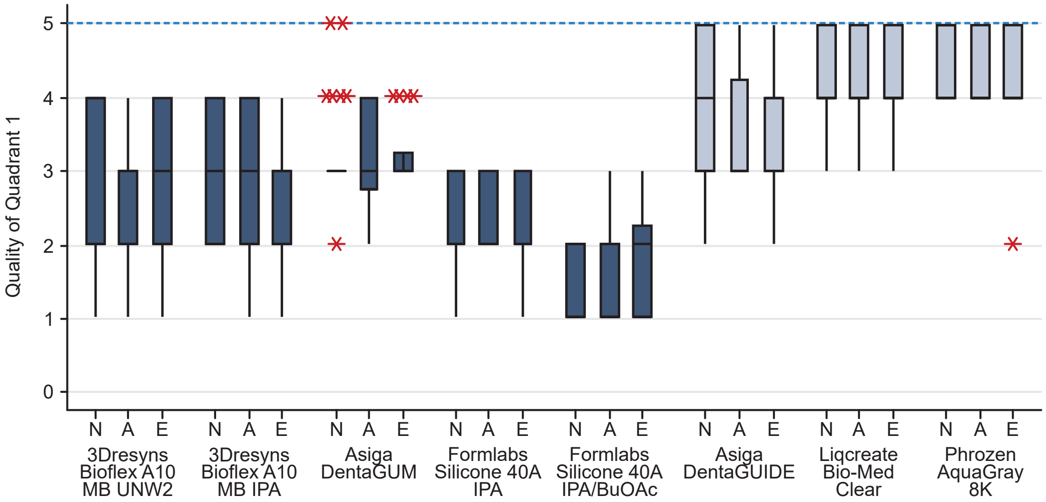


Figure C.32: The quality of the quadrant 1 area (Figure reffig:Fidelity sizingB) as determined by the manual assessment are represented as boxplots of the rigid (dark blue-gray) and elastomeric (light blue-gray) resin samples with one of the sterilization conditions: N - nonsterile, A - autoclave, E - ethanol/UV. The red asterisks (*) denote outliers. The upper and lower boundaries of the box represent the first and third quartiles. Whiskers extend from the interquartile boxes to the minimum and maximum values, excluding outliers. The blue dashed line represented the score awarded if the sample qualitatively appeared equivalent to the original CAD model (Figure 2).

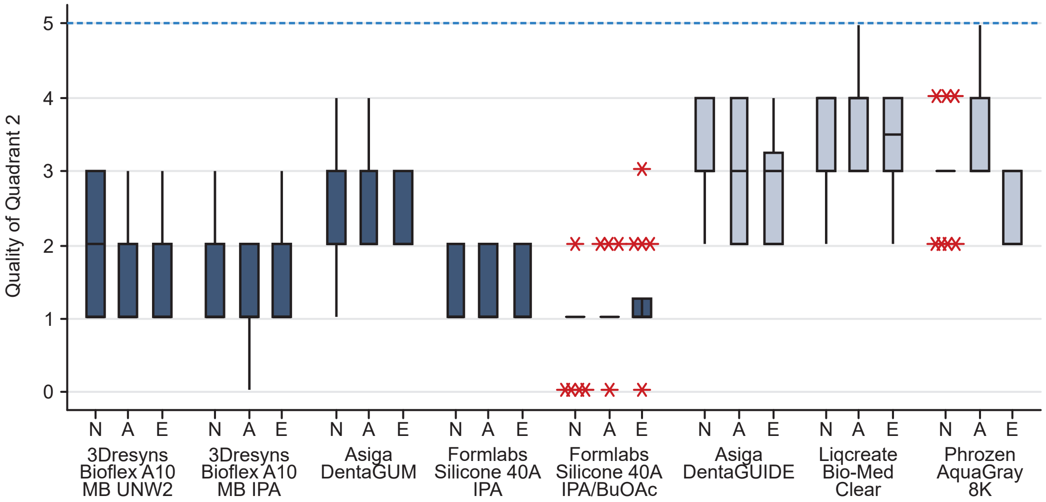


Figure C.33: The quality of the quadrant 2 area (Figure 2B) as determined by the manual assessment are represented as boxplots of the rigid (dark blue-gray) and elastomeric (light blue-gray) resin samples with one of the sterilization conditions: N - nonsterile, A - autoclave, E - ethanol/UV. The red asterisks (*) denote outliers. The upper and lower boundaries of the box represent the first and third quartiles. Whiskers extend from the interquartile boxes to the minimum and maximum values, excluding outliers. The blue dashed line represented the score awarded if the sample qualitatively appeared equivalent to the original CAD model (Figure 2).

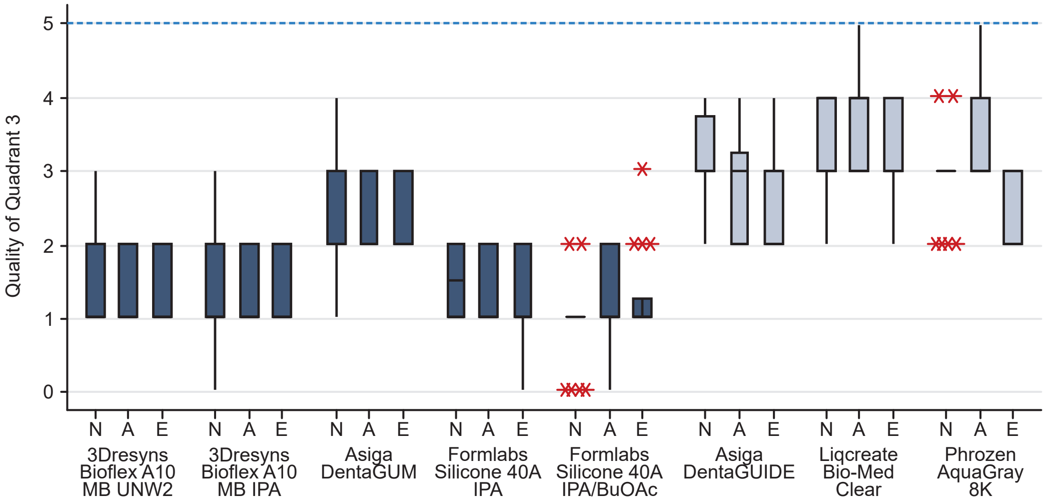


Figure C.34: The quality of the quadrant 3 area (Figure 2B) as determined by the manual assessment are represented as boxplots of the rigid (dark blue-gray) and elastomeric (light blue-gray) resin samples with one of the sterilization conditions: N - nonsterile, A - autoclave, E - ethanol/UV. The red asterisks (*) denote outliers. The upper and lower boundaries of the box represent the first and third quartiles. Whiskers extend from the interquartile boxes to the minimum and maximum values, excluding outliers. The blue dashed line represented the score awarded if the sample qualitatively appeared equivalent to the original CAD model (Figure 2).

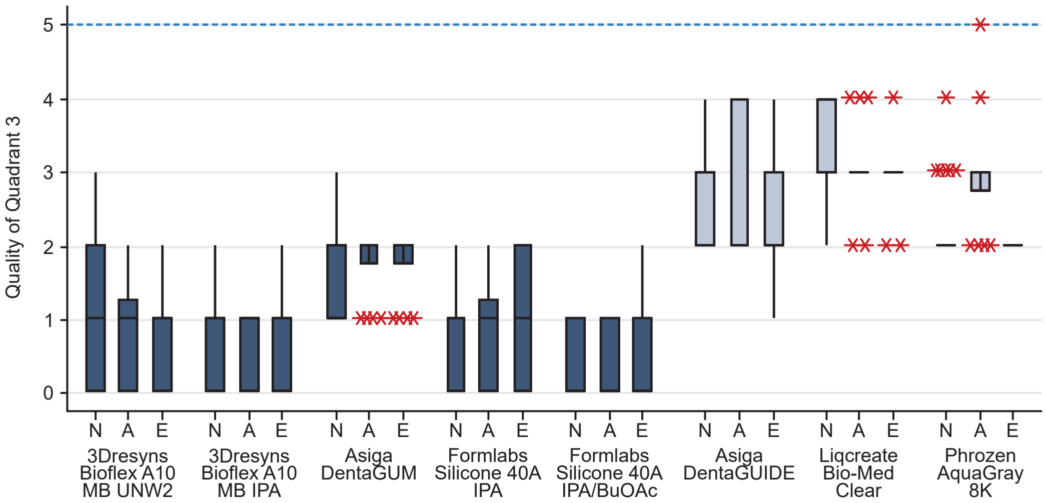


Figure C.35: The quality of the quadrant 4 area (Figure 2B) as determined by the manual assessment are represented as boxplots of the rigid (dark blue-gray) and elastomeric (light blue-gray) resin samples with one of the sterilization conditions: N - nonsterile, A - autoclave, E - ethanol/UV. The red asterisks (*) denote outliers. The upper and lower boundaries of the box represent the first and third quartiles. Whiskers extend from the interquartile boxes to the minimum and maximum values, excluding outliers. The blue dashed line represented the score awarded if the sample qualitatively appeared equivalent to the original CAD model (Figure 2).

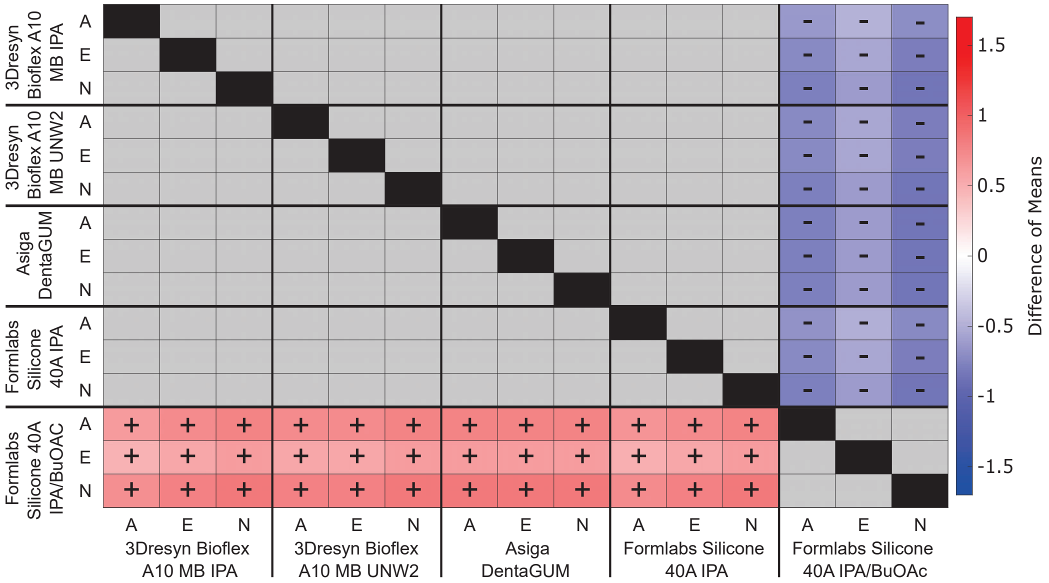


Figure C.36: Comparison of the mean quality score for the column containing the largest feature for the elastomeric resin samples. The colors and symbols represent the difference of means as calculated by subtracting the average quality of the left, vertical axis from the bottom, horizontal axis (positive value: red +, negative value: blue -). Grey boxes represent features with no significant difference. Sterilization factor: N - nonsterile, A - autoclave, E - ethanol/UV.

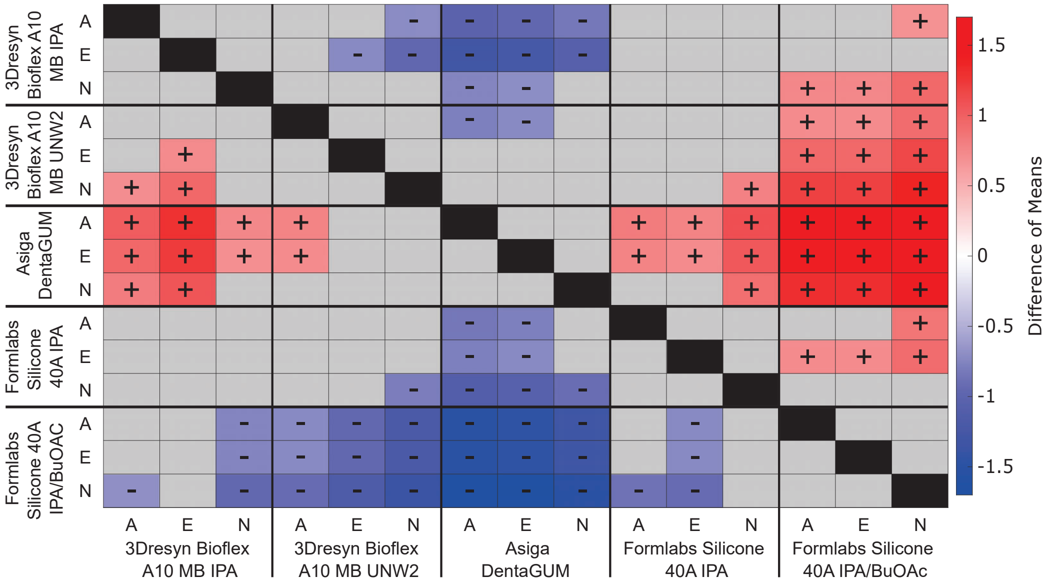


Figure C.37: Comparison of the mean quality score for the column containing the smallest feature for the elastomeric resin samples. The colors and symbols represent the difference of means as calculated by subtracting the average quality of the left, vertical axis from the bottom, horizontal axis (positive value: red +, negative value: blue -). Grey boxes represent features with no significant difference. Sterilization factor: N - nonsterile, A - autoclave, E - ethanol/UV.

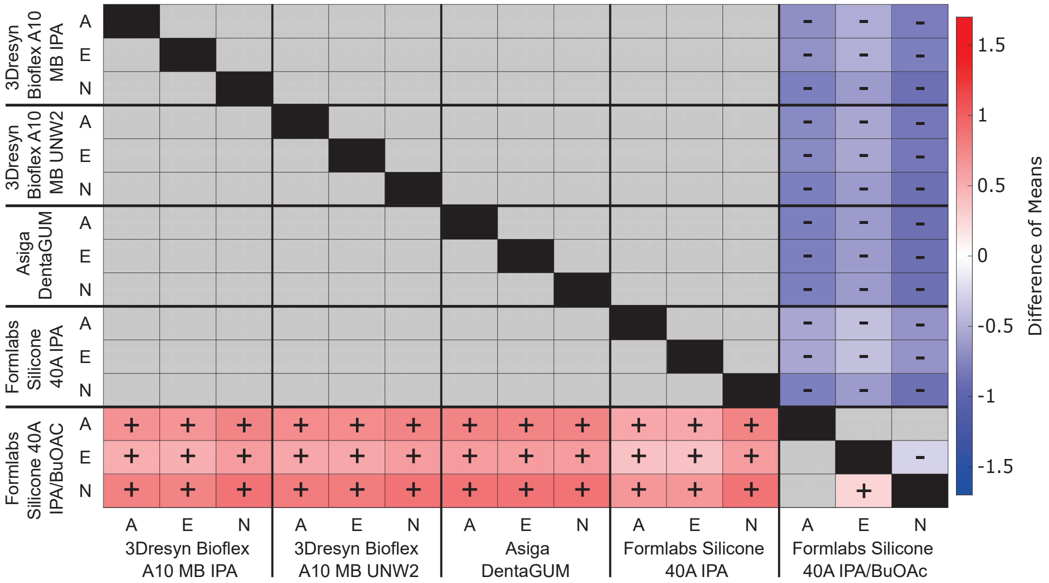


Figure C.38: Comparison of the mean quality score for the row containing the largest feature for the elastomeric resin samples. The colors and symbols represent the difference of means as calculated by subtracting the average quality of the left, vertical axis from the bottom, horizontal axis (positive value: red +, negative value: blue -). Grey boxes represent features with no significant difference. Sterilization factor: N - nonsterile, A - autoclave, E - ethanol/UV.

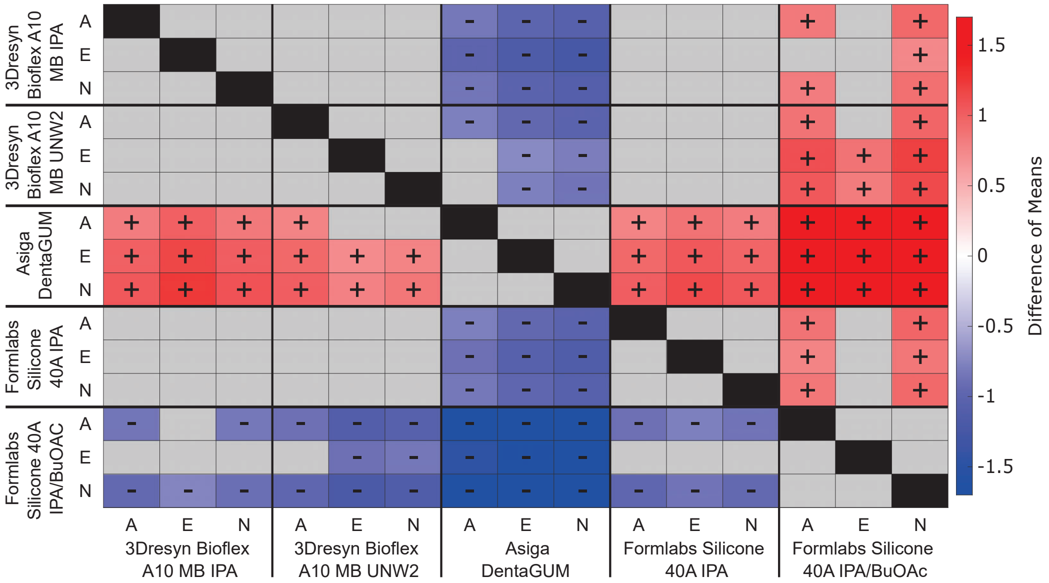


Figure C.39: Comparison of the mean quality score for the row containing the smallest feature for the elastomeric resin samples. The colors and symbols represent the difference of means as calculated by subtracting the average quality of the left, vertical axis from the bottom, horizontal axis (positive value: red +, negative value: blue -). Grey boxes represent features with no significant difference. Sterilization factor: N - nonsterile, A - autoclave, E - ethanol/UV.

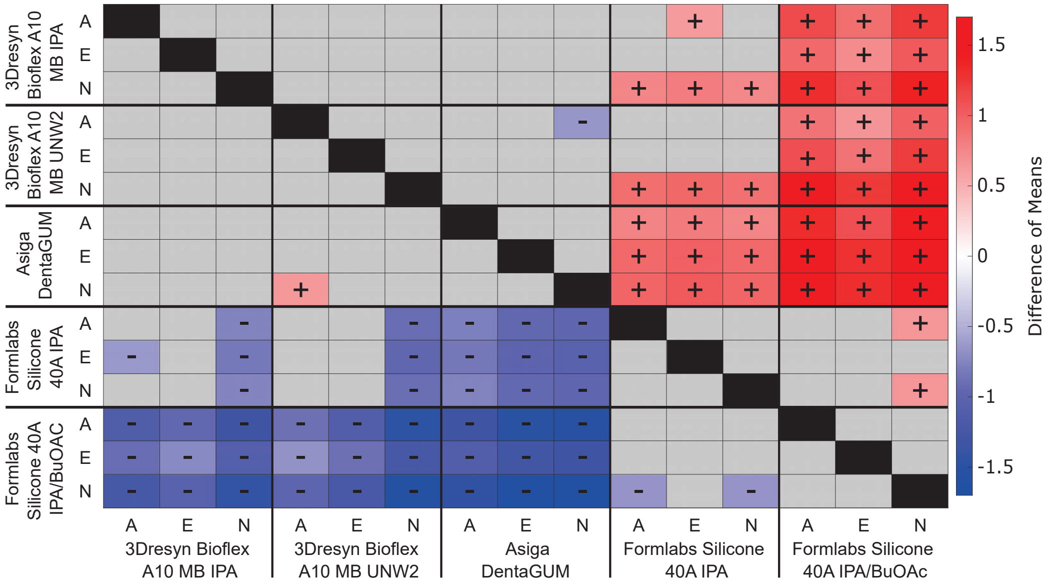


Figure C.40: Comparison of the mean quality score for quadrant 1 for the elastomeric resin samples. The colors and symbols represent the difference of means as calculated by subtracting the average quality of the left, vertical axis from the bottom, horizontal axis (positive value: red +, negative value: blue -). Grey boxes represent features with no significant difference. Sterilization factor: N - nonsterile, A - autoclave, E - ethanol/UV.

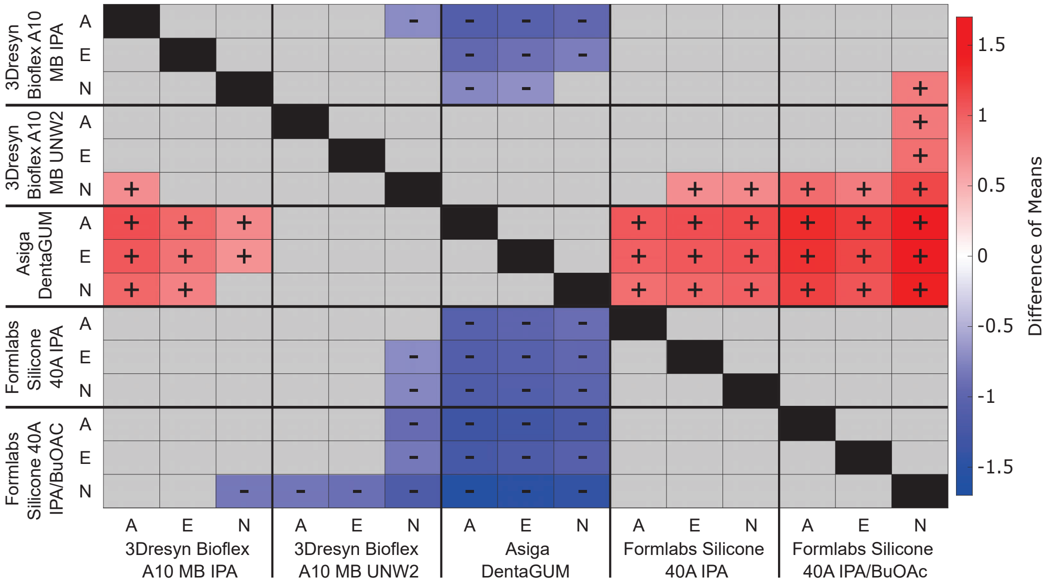


Figure C.41: Comparison of the mean quality score for quadrant 2 for the elastomeric resin samples. The colors and symbols represent the difference of means as calculated by subtracting the average quality of the left, vertical axis from the bottom, horizontal axis (positive value: red +, negative value: blue -). Grey boxes represent features with no significant difference. Sterilization factor: N - nonsterile, A - autoclave, E - ethanol/UV.

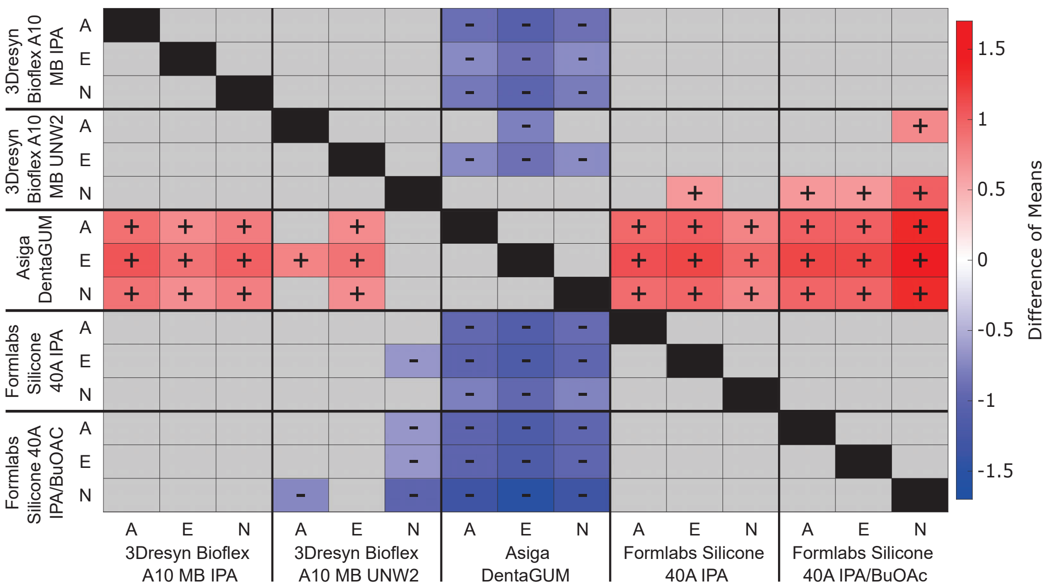


Figure C.42: Comparison of the mean quality score for quadrant 3 for the elastomeric resin samples. The colors and symbols represent the difference of means as calculated by subtracting the average quality of the left, vertical axis from the bottom, horizontal axis (positive value: red +, negative value: blue -). Grey boxes represent features with no significant difference. Sterilization factor: N - nonsterile, A - autoclave, E - ethanol/UV.

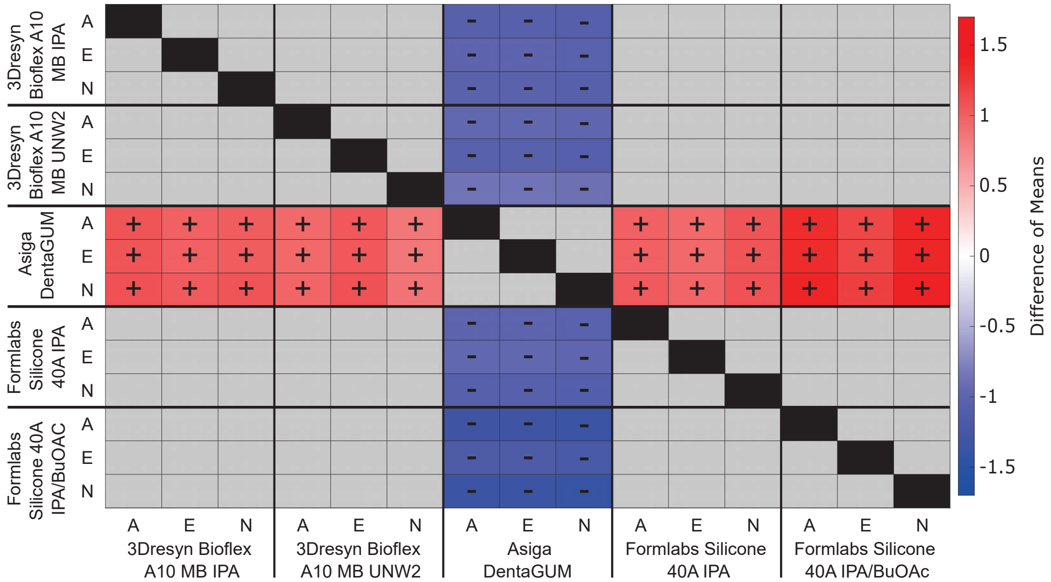


Figure C.43: Comparison of the mean quality score for quadrant 4 for the elastomeric resin samples. The colors and symbols represent the difference of means as calculated by subtracting the average quality of the left, vertical axis from the bottom, horizontal axis (positive value: red +, negative value: blue -). Grey boxes represent features with no significant difference. Sterilization factor: N - nonsterile, A - autoclave, E - ethanol/UV.

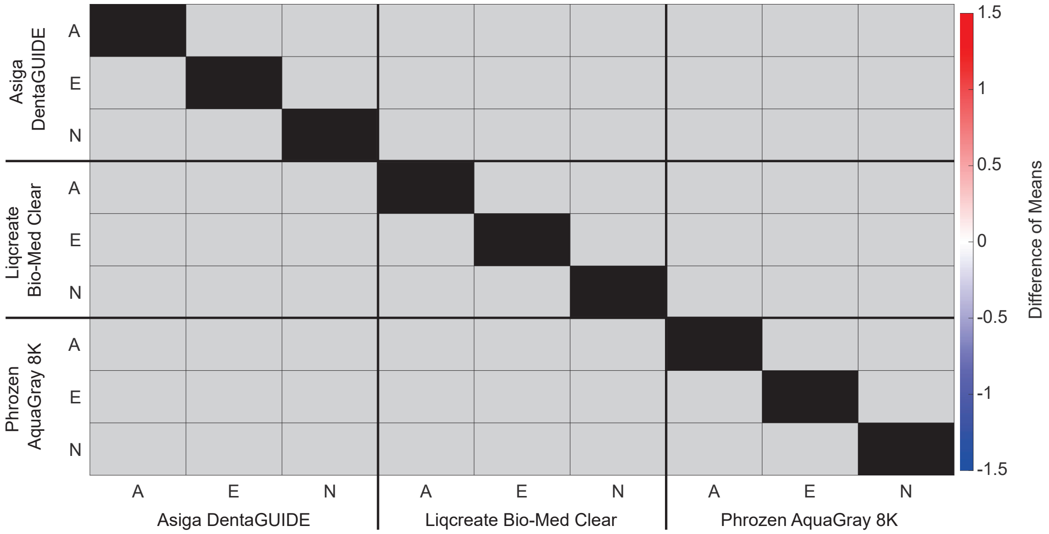


Figure C.44: Comparison of the mean quality score for the column containing the largest feature for the rigid resin samples. The colors and symbols represent the difference of means as calculated by subtracting the average quality of the left, vertical axis from the bottom, horizontal axis (positive value: red +, negative value: blue -). Grey boxes represent features with no significant difference. Sterilization factor: N - nonsterile, A - autoclave, E - ethanol/UV.

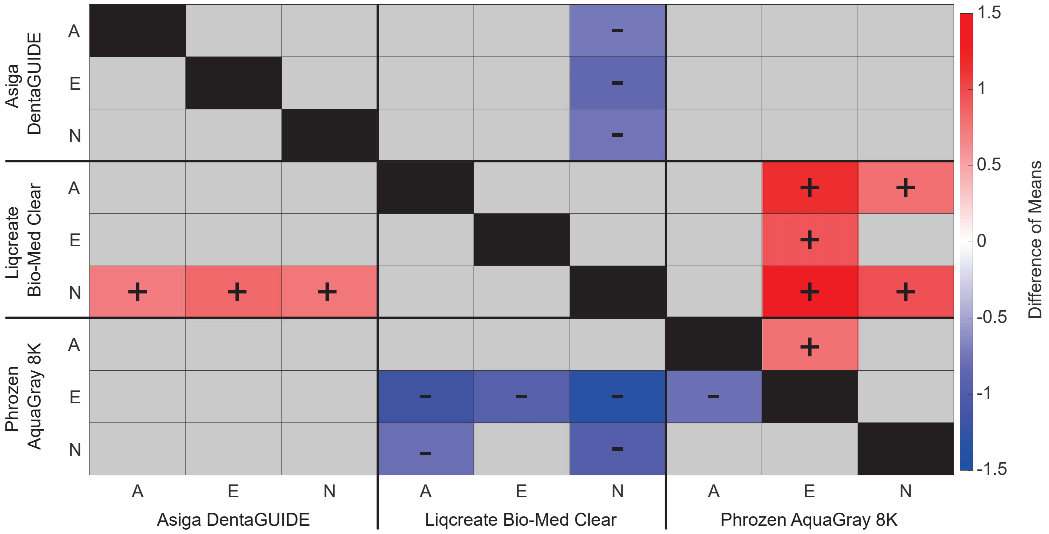


Figure C.45: Comparison of the mean quality score for the column containing smallest feature for the rigid resin samples. The colors and symbols represent the difference of means as calculated by subtracting the average quality of the left, vertical axis from the bottom, horizontal axis (positive value: red +, negative value: blue -). Grey boxes represent features with no significant difference. Sterilization factor: N - nonsterile, A - autoclave, E - ethanol/UV.

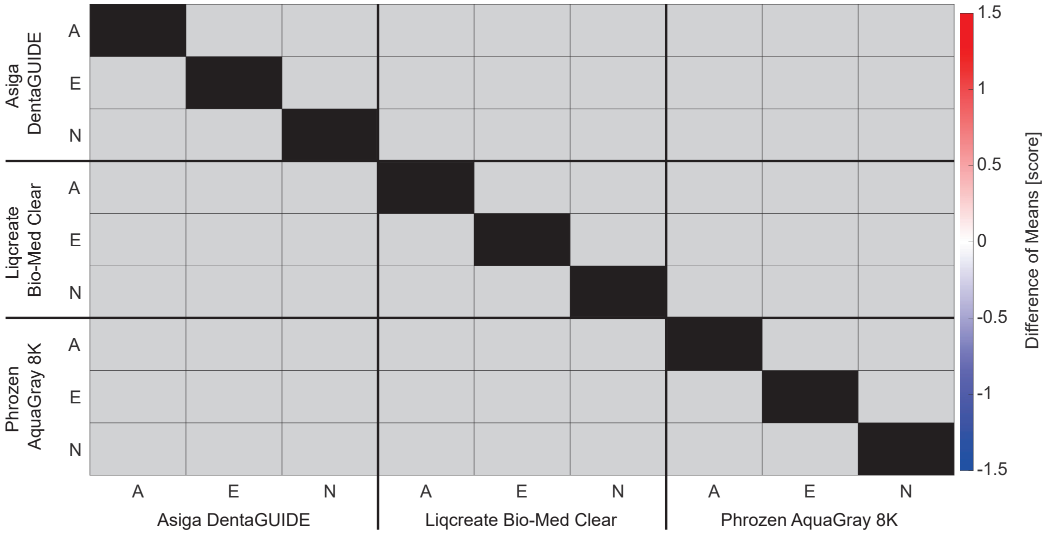


Figure C.46: Comparison of the mean quality score for the row containing the largest feature for the rigid resin samples for elastomeric resins. The colors and symbols represent the difference of means as calculated by subtracting the average quality of the left, vertical axis from the bottom, horizontal axis (positive value: red +, negative value: blue -). Grey boxes represent features with no significant difference. Sterilization factor: N - nonsterile, A - autoclave, E - ethanol/UV.

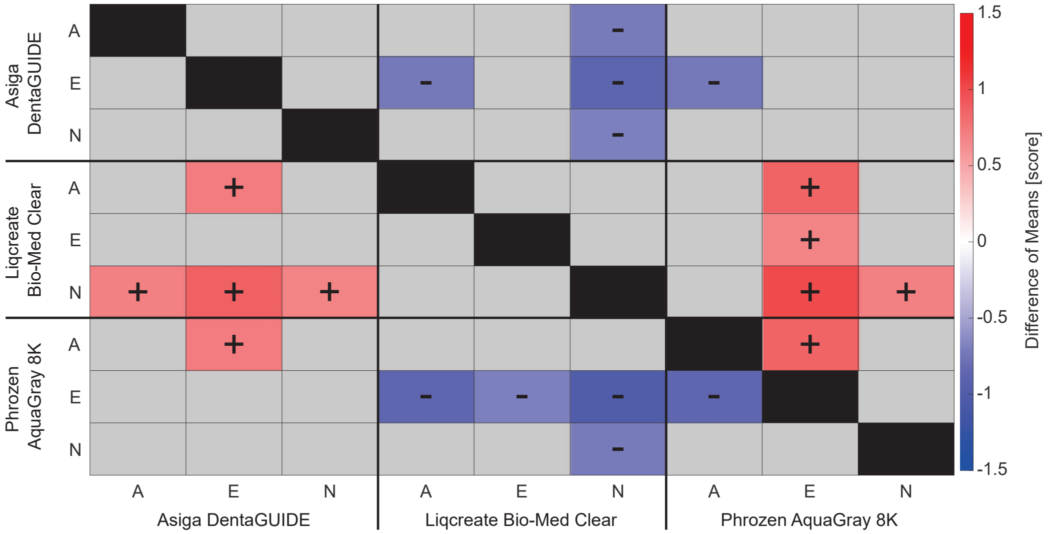


Figure C.47: Comparison of the mean quality score for the row containing the smallest feature for the rigid resins. The colors and symbols represent the difference of means as calculated by subtracting the average quality of the left, vertical axis from the bottom, horizontal axis (positive value: red +, negative value: blue -). Grey boxes represent features with no significant difference. Sterilization factor: N - nonsterile, A - autoclave, E - ethanol/UV.

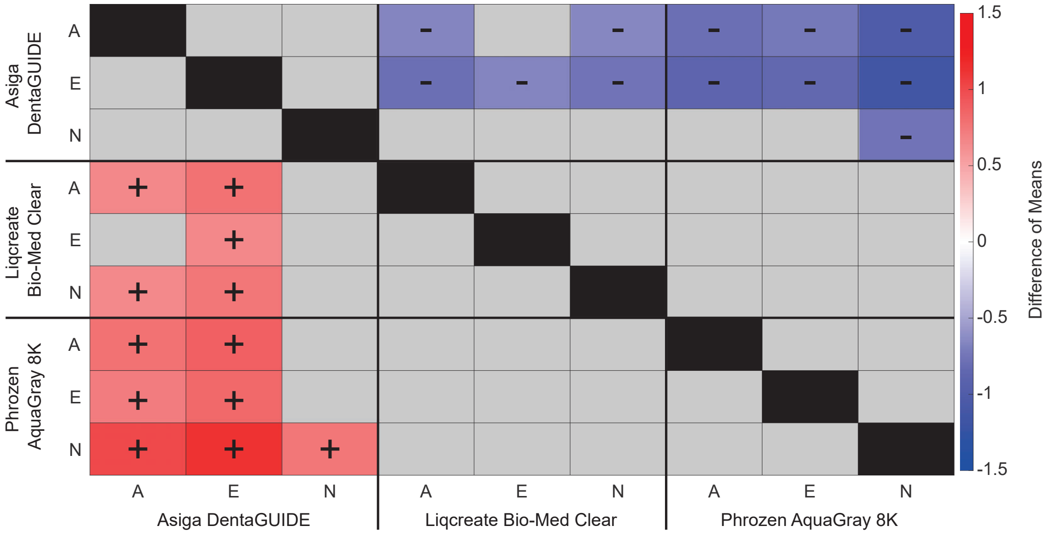


Figure C.48: Comparison of the mean quality score for quadrant 1 for the rigid resins. The colors and symbols represent the difference of means as calculated by subtracting the average quality of the left, vertical axis from the bottom, horizontal axis (positive value: red +, negative value: blue -). Grey boxes represent features with no significant difference. Sterilization factor: N - nonsterile, A - autoclave, E - ethanol/UV.

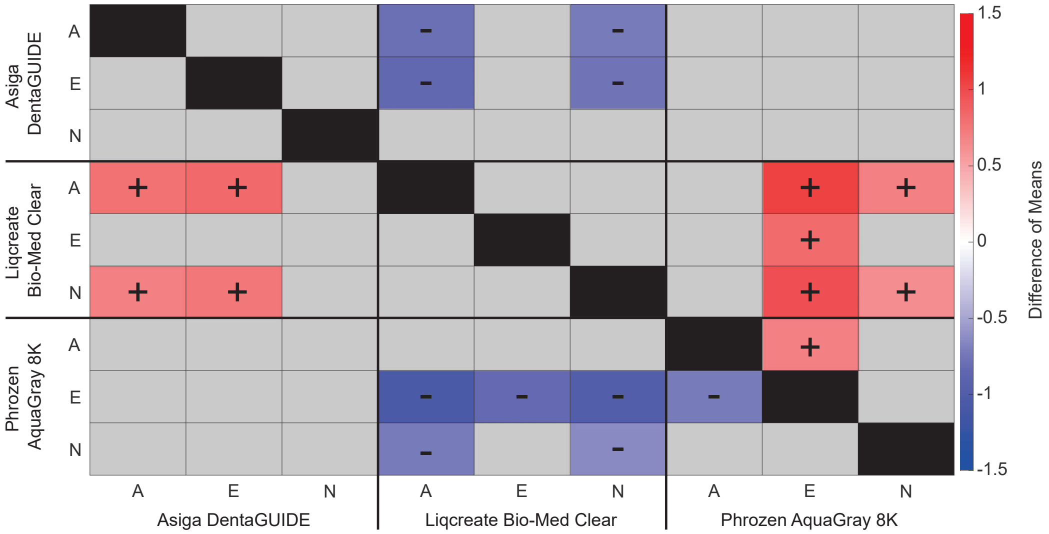


Figure C.49: Comparison of the mean quality score for quadrant 2 for the rigid resins. The colors and symbols represent the difference of means as calculated by subtracting the average quality of the left, vertical axis from the bottom, horizontal axis (positive value: red +, negative value: blue -). Grey boxes represent features with no significant difference. Sterilization factor: N - nonsterile, A - autoclave, E - ethanol/UV.

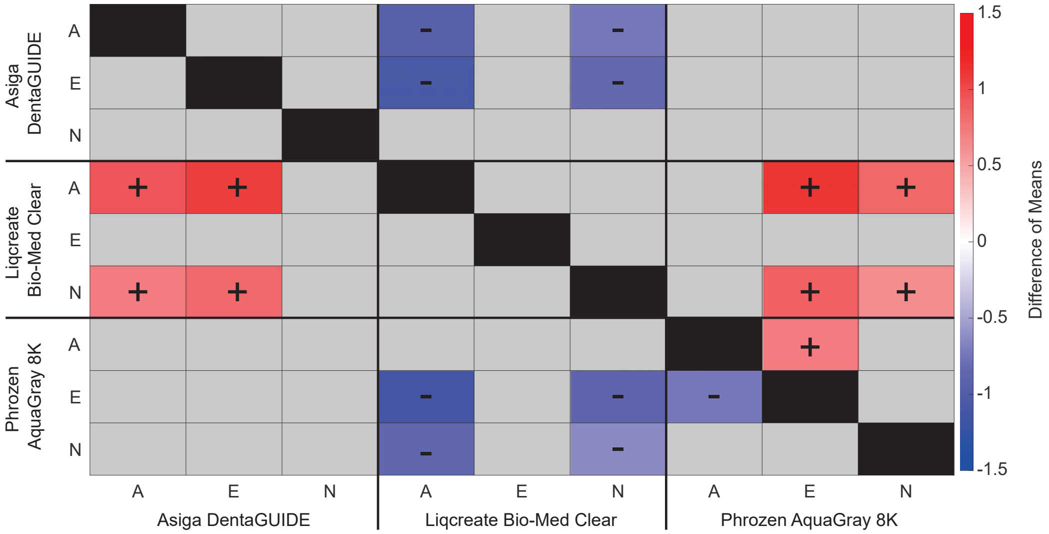


Figure C.50: Comparison of the mean quality score for quadrant 3 for the rigid resins. The colors and symbols represent the difference of means as calculated by subtracting the average quality of the left, vertical axis from the bottom, horizontal axis (positive value: red +, negative value: blue -). Grey boxes represent features with no significant difference. Sterilization factor: N - nonsterile, A - autoclave, E - ethanol/UV.

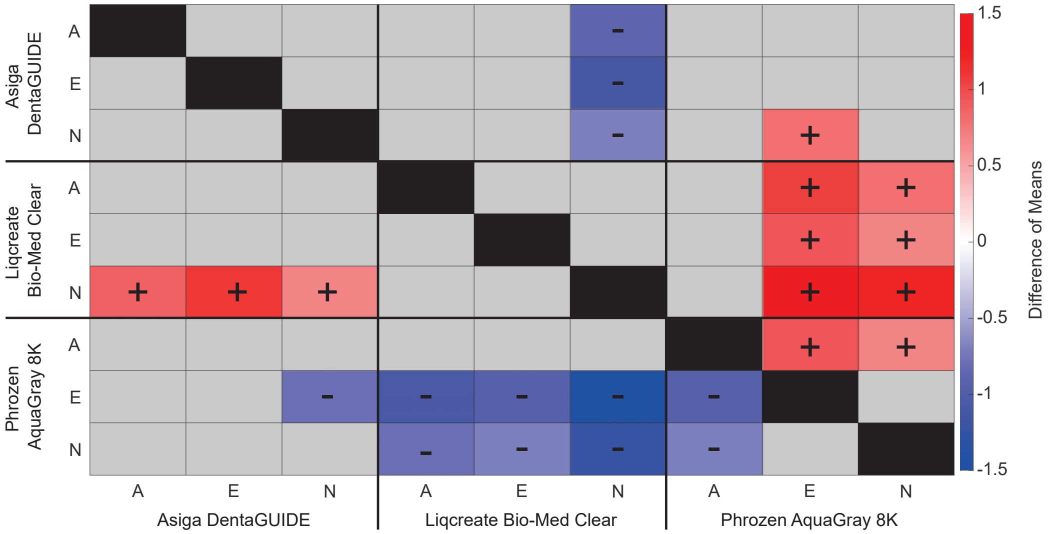


Figure C.51: Comparison of the mean quality score for quadrant 4 for the rigid resins. The colors and symbols represent the difference of means as calculated by subtracting the average quality of the left, vertical axis from the bottom, horizontal axis (positive value: red +, negative value: blue -). Grey boxes represent features with no significant difference. Sterilization factor: N - nonsterile, A - autoclave, E - ethanol/UV.
