## Supplementary material for "Cytotoxicity and Characterization of 3D-Printable Resins Using a Low-Cost Printer for Muscle-based Biohybrid Devices": Graphical Abstract

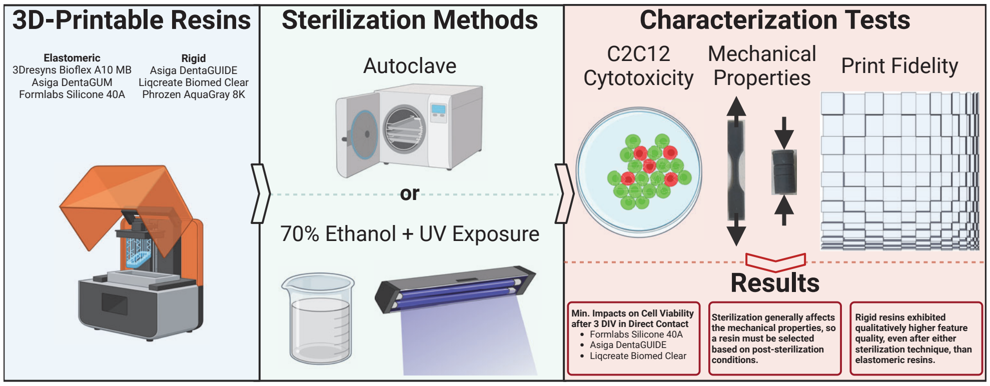
